# Pairwise Linkage Disequilibrium Estimation for Polyploids

**DOI:** 10.1101/2020.08.03.234476

**Authors:** David Gerard

**Affiliations:** Department of Mathematics and Statistics, American University, Washington, DC, 20016, USA

**Keywords:** composite, correlation, haplotypic, genotype likelihoods, LD, polyploidy

## Abstract

Many tasks in statistical genetics involve pairwise estimation of linkage disequilibrium (LD). The study of LD in diploids is mature. However, in polyploids, the field lacks a comprehensive characterization of LD. Polyploids also exhibit greater levels of genotype uncertainty than diploids, and yet no methods currently exist to estimate LD in polyploids in the presence of such genotype uncertainty. Furthermore, most LD estimation methods do not quantify the level of uncertainty in their LD estimates. Our paper contains three major contributions. (i) We characterize haplotypic and composite measures of LD in polyploids. These composite measures of LD turn out to be functions of common statistical measures of association. (ii) We derive procedures to estimate haplotypic and composite LD in polyploids in the presence of genotype uncertainty. We do this by estimating LD directly from genotype likelihoods, which may be obtained from many genotyping platforms. (iii) We derive standard errors of all LD estimators that we discuss. We validate our methods on both real and simulated data. Our methods are implemented in the R package ldsep, available on the Comprehensive R Archive Network https://cran.r-project.org/package=ldsep.

## 1 Introduction

Linkage disequilibrium (LD), the statistical association between alleles at different loci, is a fundamental quantity in statistical genetics. Estimates of LD have applications in association mapping [Devlin and Risch, 1995, Jorde, 1995, Xiong and Guo, 1997, Mackay and Powell, 2007, Farh et al., 2015, Gur et al., 2017], analyses using summary statistics from genome-wide association studies (GWAS) [Yang et al., 2012, Benner et al., 2017, Zhu and Stephens, 2017], genomic prediction [Wientjes et al., 2013, Sun et al., 2016], and population genetic studies [Slatkin, 2008, Zhu et al., 2015, Van Wyngaarden et al., 2017, Griffiths et al., 2019], among other tasks [Sved and Hill, 2018].

Many of these LD tasks are now being applied to polyploid organisms. Polyploids, organisms with more than two copies of their genome, are ubiquitous in the plant kingdom [Barker et al., 2016], predominant in agriculture [Udall and Wendel, 2006], and important drivers of evolution [Soltis et al., 2014]. As such, researchers started applying polyploid LD estimates to genotype imputation [Clark et al., 2019, Matias et al., 2019], GWAS [Barreto et al., 2019, Ferrão et al., 2020], genomic prediction [Ramstein et al., 2016, de Bem Oliveira et al., 2019, de C. Lara et al., 2019], and various other applications.

Characterizing and estimating LD in diploids is a mature field. Since the various measures of LD were proposed [Lewontin and Kojima, 1960, Lewontin, 1964, Hill and Robertson, 1968] and the basic strategies of estimation implemented [see Weir, 1996, for example], researchers have proposed extensions spanning many directions. Methods have been created to estimate LD using genotype data, rather than haplotype data, under the assumption of Hardy-Weinberg equilibrium (HWE) [Hill, 1974, Weir and Cockerham, 1979, Hui and Burt, 2020]. Composite measures of LD have been defined that are estimable using genotype data even when HWE is violated [Cockerham and Weir, 1977, Weir, 1979, Hamilton and Cole, 2004, Zaykin, 2004]. Procedures have been devised to estimate LD in the presence of genotype uncertainty [Li, 2011, Maruki and Lynch, 2014, Bilton et al., 2018, Fox et al., 2019]. Regularization procedures have been suggested to improve LD estimates [Wen and Stephens, 2010].

The research on characterizing and estimating LD in polyploids is much more limited. To date, there have basically been three approaches to estimating LD in polyploids. (i) Researchers assume they have known haplotypes (through phasing of known genotypes or otherwise) and then estimate LD using the empirical haplotype frequencies [Simko et al., 2006, Bradbury et al., 2007, Shen et al., 2016]. (ii) Researchers construct two-way tables of known genotypes and run categorical tests of association, such as Fisher’s exact test [Raboin et al., 2008, Julier, 2009, Huang et al., 2020]. (iii) Finally, researchers use standard statistical measures of association, such as the squared Pearson correlation, on the known genotypes between two loci [Björn et al., 2010, Ramstein et al., 2016, Vos et al., 2017, Sharma et al., 2018, de Bem Oliveira et al., 2019].

There are many well-developed methods to obtain phased haplotypes in diploids [Scheet and Stephens, 2006, Browning and Browning, 2007, Li et al., 2010, Swarts et al., 2014], and so method (i) above is an appealing strategy as it allows the study of association directly at the haplotypic level. However, similar advances in polyploids are relatively infant and are just now emerging in force [Su et al., 2008, Shen et al., 2016, Zheng et al., 2016, Mollinari and Garcia, 2019]. An additional limitation is that these phasing approaches usually require access to a reference genome, which is not always available or necessary in many modern next-generation sequencing pipelines [Lu et al., 2013].

A greater concern is that all of the polyploid LD estimation approaches listed above assume genotypes are known without error. In polyploids, this assumption is incorrect. Even though there have been gains in the accuracy of genotyping methods [Voorrips et al., 2011, Serang et al., 2012, Mollinari and Serang, 2015, Maruki and Lynch, 2017, Schmitz Carley et al., 2017, Blischak et al., 2018, Gerard et al., 2018, Clark et al., 2019, Gerard and Ferrão, 2019, Zych et al., 2019] the issue of genotype uncertainty in polyploids is still severe and much more so than in diploids. In Gerard et al. [2018], we found that genotyping error rates in diploids can be reduced to less than 0.05 at a read-depth of around 5×, but similar error rates could not be achieved for hexaploids in some scenarios until one had read-depths in the many-thousands, an unrealistic scenario for most applied researchers.

In this paper, we provide various methods to estimate LD. After reviewing measures of haplotypic LD in Section 2.1, in Section 2.2 we derive a procedure to estimate haplotypic LD in autopolyploids in the presence of genotype uncertainty under the assumption of HWE. We do this by estimating LD directly using genotype likelihoods. In allopolyploids, organisms that exhibit partial preferential pairing, or populations that violate the random-mating assumption, these estimates are not appropriate. Thus, in Section 2.3 we define various “composite” measures of LD that generalize the composite measures proposed in Cockerham and Weir [1977]. These measures turn out to be functions of the statistical moments of the genotypes. In Section 2.4 we provide methods to estimate these composite measures of LD in the presence of genotype uncertainty by directly using genotype likelihoods. To reduce the number of parameters we estimate, we propose a novel and flexible class of distributions over the genotypes. We validate our methods both in simulations (Section 3.1) and on real data (Section 3.3). All analyses are reproducible (https://doi.org/10.6084/m9.figshare.12765803) and all methods are available in the ldsep R package on the Comprehensive R Archive Network (https://cran.r-project.org/package=ldsep).

## 2 Materials and Methods

### Notation

The following contains the notation conventions used throughout this manuscript. We will denote scalars by non-bold letters (*a* or *A*), vectors by bold lower-case letters (***a***), and matrices by bold upper-case letters (***A***). We let Latin letters denote haplotypic measures of LD (*D, D*′, and *r*), while we let Greek letters denote composite measures of LD (Δ, Δ′, and *ρ*). Estimates of population parameters will be denoted with a hat 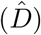.

### 2.1 Measures of haplotypic LD

The original definitions of LD quantify the statistical association between alleles located at different loci on the same haplotype. In diploids, since only a single haplotype is present in a gamete, such associations are sometimes referred to as “gametic LD” [Hedrick et al., 1978]. However, in autopolyploids, a single gamete might contain multiple haplotypes (on the multiple homologues), and so a more appropriate term for such LD in polyploids might be “haplotypic”. For this manuscript, we will use the terminology “haplotypic LD” to refer to what is known in the diploid literature as “gametic LD”.

Various measures of haplotypic LD have been proposed in the literature, each possessing relative strengths and weaknesses for interpreting the association between loci [Hedrick, 1987, Devlin and Risch, 1995]. In this section, we briefly review these common measures of haplotypic LD.

Perhaps the three most commonly used measures are the LD coefficient *D* [Lewontin and Kojima, 1960], the standardized LD coefficient *D*′ [Lewontin, 1964], and the Pearson correlation *r* [Hill and Robertson, 1968]. In this manuscript, we will explore these measures of LD only in the context of biallelic loci, and leave for future work the task of generalizing multiallelic loci methods to polyploids from diploids [Thomson and Single, 2014, Okada, 2018, e.g.]. To define these terms, let A and a be the reference and alternative alleles, respectively, at locus 1. Similarly let B and b be the reference and alternative alleles at locus 2. The LD coefficient is defined to be the difference between the true haplotype frequency and the haplotype frequency under independence:

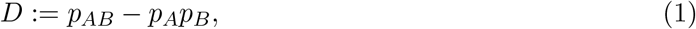

where *p*_*AB*_ denotes the frequency of haplotype AB, *p*_*A*_ denotes the allele frequency of A, and *p*_*B*_ denotes the allele frequency of B. Note that this definition necessarily implies that

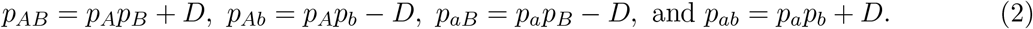

The range of *D* is constrained by the allele frequencies [Lewontin, 1964] by

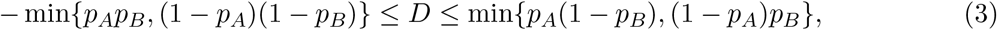

and so Lewontin [1964] suggested the standardized LD coefficient:

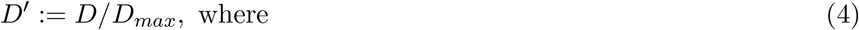

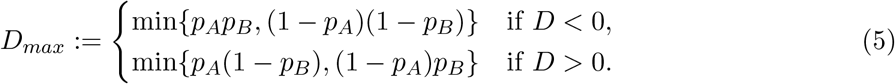

The *D*′ coefficient is free to vary between -1 and 1, though it still depends on the loci-specific allele frequencies [Lewontin, 1988]. Finally, the Pearson correlation between loci is

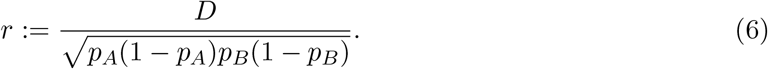

It is common to report not *D, D*′, and *r*, but rather their absolute values or their squares. This is because the sign of the LD between loci depends on the mostly arbitrary labels of the alternative and reference alleles.

### 2.2 Pairwise haplotypic LD estimation in autopolyploids under HWE

In this section, we consider estimating haplotypic LD from a population of autopolyploid individuals under HWE in a species that exhibits only bivalent pairing (though we note that the following methods are rather robust to the bivalent assumption, Section S12). We will begin by deriving the joint distribution of genotypes at two loci conditional on the LD coefficient and the allele frequencies at both loci. These genotype distributions will then be used, along with user-provided genotype likelihoods, to develop a procedure to estimate LD in the presence of genotype uncertainty. Genotype likelihoods, the probability of the data given the genotypes, may be obtained from many genotyping programs (at least up to a proportional constant which does not depend on any parameters, which may be used in what follows; see Section S14) [McKenna et al., 2010, Gerard et al., 2018, Clark et al., 2019, Gerard and Ferrão, 2019, e.g.]. Maximum likelihood theory will be used to derive standard errors of these estimators.

Let *X*_*iAB*_, *X*_*iAb*_, *X*_*iaB*_, and *X*_*iab*_ be the number of AB, Ab, aB, and ab haplotypes (respectively) in individual *i*. If each autopolyploid individual is *K*-ploid, then under HWE we have

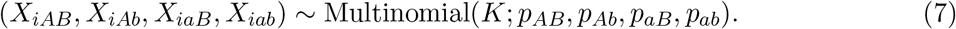

Researchers typically do not have access to individual haplotypes as in (7). Rather, most methods assume that researchers have available the dosages of each allele, which we call the genotypes. Let *G*_*iA*_ := *X*_*iAB*_ + *X*_*iAb*_ and *G*_*iB*_ := *X*_*iAB*_ + *X*_*iaB*_ be the number of reference alleles at loci 1 and 2, respectively, in individual *i*. Then we may sum over the *X*’s to obtain the joint distribution of *G*_*iA*_ and *G*_*iB*_ (Supplementary Section S1).

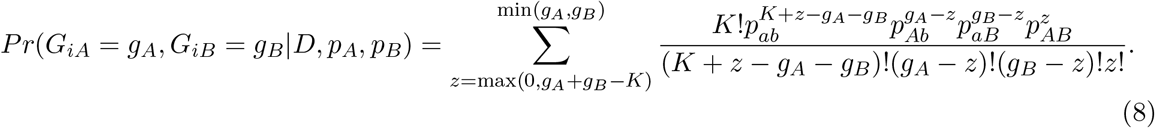

Equation (8) is conditional on *D, p*_*A*_, and *p*_*B*_ due to the relations in (2). Equation (8) contains the distribution of the genotypes under HWE, conditional on *D* and allele frequencies, and generalizes the formulas in Table 1 of Hill [1974] to autopolyploids.

**Table 1:** Summary of LD estimators. The 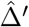 estimators all estimate 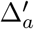 (19), *not* 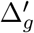 (17).

| Estimators | Type | Input | Notes |
| --- | --- | --- | --- |
| $\hat{D}_g, \hat{D}'_g, \hat{r}_g$ | Haplotypic | Genotypes | |
| $\hat{D}_{gl}, \hat{D}'_{gl}, \hat{r}_{gl}$ | Haplotypic | Genotype Likelihoods | A Dirichlet(2, 2, 2, 2) prior is placed on the haplotype frequencies. |
| $\hat{\Delta}_{mom}, \hat{\Delta}'_{mom}, \hat{\rho}_{mom}$ | Composite | Genotypes | Can accept continuous genotypes as input. |
| $\hat{\Delta}_{gc}, \hat{\Delta}'_{gc}, \hat{\rho}_{gc}$ | Composite | Genotype Likelihoods | Uses the general categorical class of genotype distributions. |
| $\hat{\Delta}_{pn}, \hat{\Delta}'_{pn}, \hat{\rho}_{pn}$ | Composite | Genotype Likelihoods | Uses the proportional bivariate normal class of genotype distributions. |

If the genotypes were known without error, then we could estimate *D* (along with *p*_*A*_ and *p*_*B*_) by maximum likelihood estimation (MLE) using the following log-likelihood:

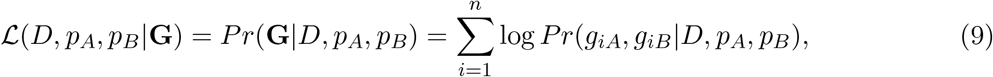

where **G** = (**g**_1_, …, **g**_*n*_) and **g**_*i*_ = (*g*_*iA*_, *g*_*iB*_) are the observed genotypes for individual *i*. Since *D*′ and *r* are both functions of *D, p*_*A*_, and *p*_*B*_ (4)–(6), this would also yield the MLEs of *D*′ and *r*. We have implemented maximizing (9) in our software using gradient ascent. We call the resulting MLEs 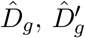 and 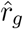 for “genotypes”.

However the genotypes are not known without error [Gerard et al., 2018]. Maruki and Lynch [2014] and Bilton et al. [2018] accounted for this in diploid sequencing data by adaptively estimating the sequencing error rate at each locus. However, their models ignore important features concerning polyploid sequencing data, particularly allele bias and overdispersion [Gerard et al., 2018]. Li [2011] and Fox et al. [2019] take the more modular approach of allowing the input of genotype likelihoods for diploids derived from any genotyping software. Letting the input be genotype likelihoods allows the use of different genotyping platforms and software that may account for different features of the data. This also results in greater generalizability since different data types (e.g., microarrays [Fan et al., 2003], next-generation sequencing [Baird et al., 2008, Elshire et al., 2011], or mass spectrometry [Oeth et al., 2009]) can all result in genotype likelihoods that may then be fed into these LD estimation applications. We thus take this approach and consider estimating LD from genotype likelihoods.

We will now describe a procedure to estimate LD while integrating over genotype uncertainty using genotype likelihoods. Let us denote the data for individual *i* at loci 1 and 2 by *y*_*iA*_ and *y*_*iB*_, respectively. Let *p*(*y*_*iA*_|*g*_*A*_) and *p*(*y*_*iB*_|*g*_*B*_) be the probabilities of the data at loci 1 and 2 given genotypes *g*_*A*_ and *g*_*B*_. These probabilities are the genotype likelihoods and are assumed to be provided by the user. All of the results in this manuscript use the genotype likelihoods from the updog software [Gerard et al., 2018, Gerard and Ferrão, 2019], but we do not require this in the methods below and any genotyping software may be used as long as it returns genotype likelihoods [McKenna et al., 2010, Clark et al., 2019, e.g.]. The log-likelihood of *D, p*_*A*_, and *p*_*B*_ given the data is:

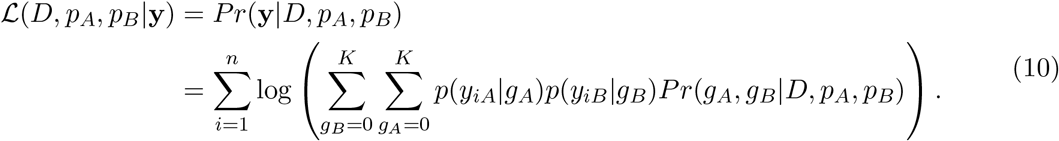

We developed an expectation-maximization (EM) algorithm [Dempster et al., 1977] to maximize (10) and estimate haplotype frequencies and, thus, LD (Section S2). For diploids (*K* = 2), this estimation procedure reduces down to that proposed in Li [2011] and implemented in the ngsLD software [Fox et al., 2019] (Figure S1). However, we found it more efficient to maximize likelihood (10) using gradient ascent. Details of this optimization are also presented in Section S2.

Standard errors are important for hypothesis testing [Brown, 1975], read-depth suggestions [Maruki and Lynch, 2014], and hierarchical shrinkage (a statistical technique to borrow strength between SNPs and improve estimation performance) [Dey and Stephens, 2018]. However, most LD estimation methods do not return standard error estimates. In Section S8, we provide a description for how to obtain such estimates using standard maximum likelihood theory. Our software returns these standard errors.

### 2.3 Composite measures of LD

The methods in Section 2.2 were developed under the assumption of HWE in autopolyploids. The goal in this section is to develop measures of LD that are (i) valid measures of association when this autopolyploid HWE assumption is violated (e.g. in the case of allopolyploids), (ii) estimable even when only genotype information is provided, and (iii) reduced to the haplotypic measures of LD (Section 2.1) when HWE is satisfied in autopolyploids. These measures will be called “composite” measures of LD, as they account for associations not just at the haplotypic level.

The easiest composite measure to obtain that satisfies our goals is that which generalizes *r* (6). Let the random variables *G*_*A*_ and *G*_*B*_ be the genotypes of a randomly selected *K*-ploid individual at loci 1 and 2. Then, quite simply, the composite measure of correlation is the Pearson correlation between genotypes:

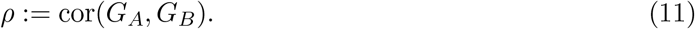

We prove in Section S3 that *ρ* equals *r* when HWE is satisfied in autopolyploids.

In diploids, Cockerham and Weir [1977] (attributing their result to Peter Burrows’ unpublished work) suggested a composite measure of LD which they defined as the sum of haplotypic LD and non-haplotypic LD. This sum depends only on genotype frequencies, not the haplotype frequencies, and is thus estimable even under general violations from HWE. Under HWE, this composite LD measure was equal to *D*, and so Weir [1979] suggested its use also when HWE was satisfied. Let *q*_*ij*_ denote the probability of a randomly selected individual containing genotype *i ∈* {0, 1, 2} on locus 1 and *j ∈* {0, 1, 2} on locus 2. Then the coefficient as defined in Cockerham and Weir [1977] is

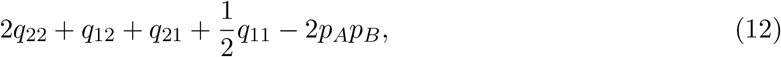

Where 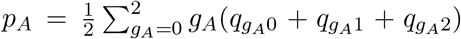 and 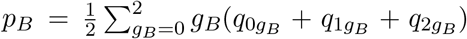 are the frequencies of the A allele at locus 1 and B allele at locus 2, respectively. It is easy to show that (12) is actually 1*/*2 the covariance of the genotypes [Weir, 2008, Rogers and Huff, 2009, Section S4]. This immediately suggests a generalized composite LD coefficient for polyploids:

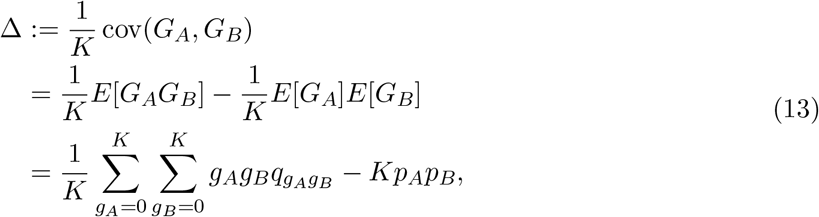

Where

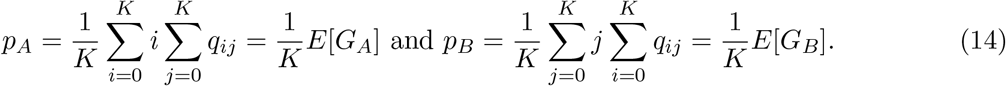

In Section S3, we show that under HWE in autopolyploids, Δ (13) equals *D* (1). We provide a decomposition of Δ in Section S7 that generalizes the decomposition in Cockerham and Weir [1977].

We will now discuss how to define a composite measure that generalizes *D*′ for polyploids. Recall that *D*′ = *D/D*_*max*_, where *D*_*max*_ is the maximum value of *D* while fixing the observed allele frequencies at both loci. We could define Δ′ = Δ*/*Δ_*max*_, where Δ_*max*_ is the maximum value of Δ. However, it is not immediately clear what observed property of the genotype distributions we should fix during this maximization. When we are finding *D*′, fixing the allele frequencies can be considered both (i) fixing the distributions of alleles and (ii) fixing the expected numbers of alleles. And so, for our generalization, we will consider in turn fixing both (i) the marginal probabilities of genotypes and (ii) the expected genotypes, while finding the maximum value of Δ.

This first approach generalizes that of Zaykin [2004] and Hamilton and Cole [2004] from diploids to polyploids. They found the maximum value of Δ in closed-form for diploids while conditioning on the marginal distributions of genotypes at both loci. For our generalization, we note that we can formulate this maximization problem as a linear program [Nocedal and Wright, 2006] and so can solve it efficiently for any ploidy level. Specifically, we seek to find the *q*_*ij*_’s that solve the following maximization problem:

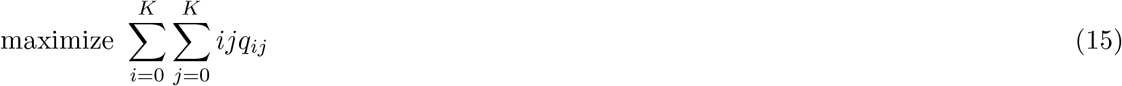

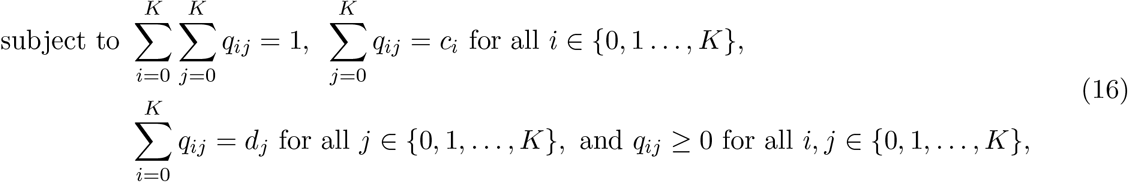

where *c*_*i*_ and *d*_*j*_ are the provided marginal probabilities of genotype *i* at locus 1 and genotype *j* at locus 2, respectively. Since (15)–(16) formulates a standard linear program, many out-of-the-box optimization software packages may be used to solve it [Berkelaar et al., 2016, e.g.]. The optimal values of the *q*_*ij*_’s may then be used to find the maximum value of Δ using (13), which can then be used to normalize Δ. If Δ is negative then we instead minimize (15) and normalize Δ by the absolute value of the minimum. Specifically, let Δ_*m*_ be the maximum of value of (13) when Δ > 0, or the absolute value of the minimum of (13) when Δ *<* 0, where we use constraints (16). Then we define

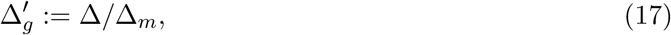

where the subscript is for “genotype frequency”. See Section S6 for a discussion on a different algorithm to obtain Δ_*m*_.

For diploids, Zaykin [2004] noted that normalizing by the maximum covariance conditional on the marginal distributions of genotypes at each locus did not result in *D*′ when when the population was in HWE. The same is true for polyploids, though we note that under HWE there appears to be a simple piecewise-linear relationship between *D*′ and 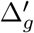 (Figure S2). As such, Zaykin [2004] in diploids also recommended the second approach of maximizing the covariance conditional on the expected genotype. We could also formulate these bounds as a linear program. However, we can actually represent the bounds on Δ (13) in closed-form (Theorem S2):

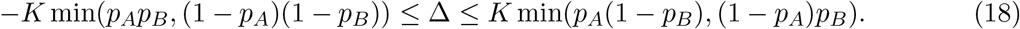

Equation (18) would suggest dividing Δ by *K* min(*p*_*A*_*p*_*B*_, (1−*p*_*A*_)(1−*p*_*B*_)) when Δ *<* 0, and dividing Δ by *K* min(*p*_*A*_(1 − *p*_*B*_), (1 − *p*_*A*_)*p*_*B*_) when Δ > 0. This would result in a composite measure of LD that is bounded within [−1, 1]. However, the resulting measure would still not equal *D*′ when HWE is satisfied. Thus, we prefer the following composite measure of LD

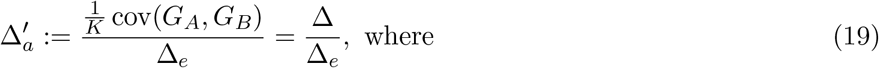

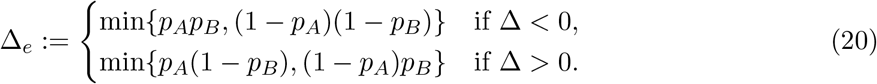

We prove in Section S3 that this 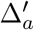 (for “allele frequency”) is equal to *D*′ under HWE in autopolyploids but is still estimable when HWE is violated. For the rest of this manuscript, we will only consider Δ 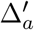 and not 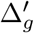.

Though a perceived disadvantage of (19) would be that it is constrained to fall within [−*K, K*] rather than [−1, 1], we find it compelling that 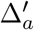 can be viewed as a direct generalization of *D*′ and is equal to *D*′ when HWE is satisfied in autopolyploids. However, a researcher may always divide (19) by *K* to obtain a measure that is constrained to be within [−1, 1].

### 2.4 Estimating composite measures of LD

When genotypes are known, it would be appropriate to use the sample moments of genotypes to estimate *ρ* (11), Δ (13), and 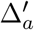 (19). When genotypes are not known, it is natural to plug-in a good estimator of the genotypes, such as the posterior means, into the sample moments. The typical methods of covariance and correlation standard errors may then be used (Section S10). This is fast and so the sample correlation of posterior genotypes has been used in the literature [Clark et al., 2019, Fox et al., 2019] (though not with the capability of returning standard errors). We will denote such estimators by 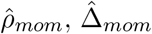, and 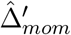 for “moment-based”.

However, as we will see in Section 3.1, using the posterior mean in such moment-based estimators results in LD estimates that are biased low. We can explain this by the law of total covariance:

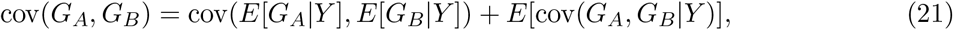

where *Y* contains the data. The covariance of posterior means is only the first term on the right hand side of (21), and so does not account for the posterior covariance of genotypes. Thus, the covariance of the posterior means is necessarily a biased estimator for Δ.

When genotypes are not known we can still estimate these composite LD measures by first estimating the *q*_*ij*_’s in (13) using maximum likelihood and then using these estimates to obtain the MLEs for *ρ*, Δ, and 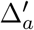. The likelihood to be maximized is

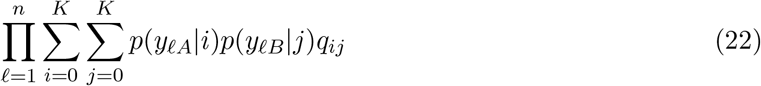

One EM step to maximize (22) consists of

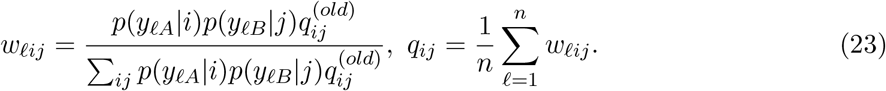

We will call the resulting MLEs 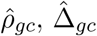,and 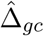 for “general categorical”, as the *q*_*ij*_’s are allowed to vary over the space of general categorical distributions. Descriptions of obtaining standard errors for these estimators are provided in Section S8.

Equation (22) uses a general categorical distribution with support over the possible genotype pairs. This distribution over genotype pairs is the most flexible possible, but yields (*K* + 1)^2^ − 1 parameters to estimate. Adding constraints over the space of possible genotype distributions can improve estimation performance as long as the genotype frequencies follow these constraints. Gerard and Ferrão [2019] introduced the proportional normal distribution, a very flexible class of distributions with support over genotypes at one locus. We now generalize this distribution to include support over the genotypes at two loci. Let ***µ*** *∈* ℝ^2^ and let **Σ** *∈* ℝ^2×2^ be a positive definite matrix. Then the proportional bivariate normal distribution with support over {0, 1, …, *K*}^2^ is:

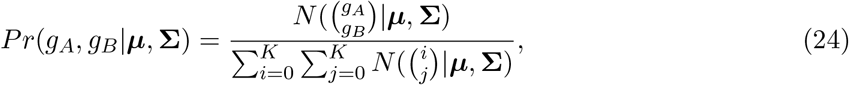

where *N* (***x***|***µ*, Σ**) is the bivariate normal density evaluated at ***x*** with mean ***µ*** and covariance matrix **Σ**. This distribution, though seemingly *ad hoc*, can be seen as a generalization of the distribution of genotypes under HWE (Figure S3). Though its use for modeling genotypes when HWE is violated can be rationalized by the flexible shapes of genotype distributions it is capable of representing (Figure S4).

Using the proportional bivariate normal distribution, the log-likelihood to be maximized is:

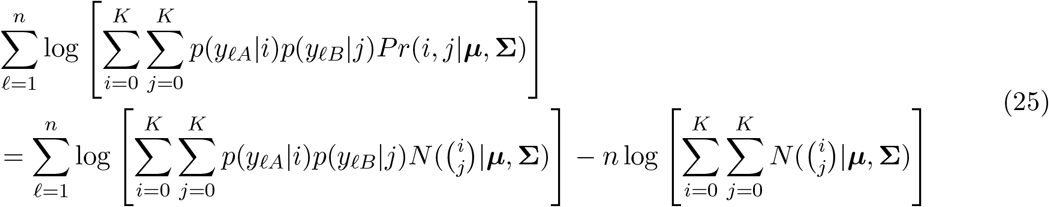

We can maximize (25) using gradient ascent. Asymptotic standard errors may be obtained using standard results from maximum likelihood theory (Section S8). We will denote the resulting MLEs by 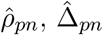and 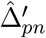 for “proportional normal”.

A summary of the various estimators we have proposed in this paper are presented in Table 1.

## 3 Results

### 3.1 Pairwise LD simulations under HWE

In this section we run simulations where genotypes are generated for autopolyploids that exhibit bivalent pairing under HWE. Under HWE for autopolyploids, both the haplotypic (Section 2.2) and the composite (Section 2.4) estimators are valid estimators of haplotypic LD.

These are the steps of a single simulation replication. Given the major allele frequencies (*p*_*A*_ and *p*_*B*_) and Pearson correlation *r* (6), the haplotype frequencies (*p*_*AB*_, *p*_*Ab*_, *p*_*aB*_, *p*_*ab*_) are uniquely identified. For individual *i ∈* {1, 2, …, *n* = 100} at a given ploidy level *K*, we simulated the number of each haplotype they contained (*X*_*iAB*_, *X*_*iAb*_, *X*_*iaB*_, *X*_*iab*_) given the haplotype frequencies using (7). Genotypes were calculated by *G*_*iA*_ := *X*_*iAB*_ + *X*_*iAb*_ and *G*_*iB*_ := *X*_*iAB*_ + *X*_*iaB*_. Given these genotypes, read-counts were simulated using `updog’s rflexdog()` function at a specified read-depth, a 0.01 sequencing error rate, no allele bias, and an overdispersion value of 0.01. Updog was then used to generate genotype likelihoods, posterior mode genotypes, and posterior mean genotypes. These outputs were fed into `ldsep` to provide the estimators listed in Table 1. The parameters that varied within the simulation were: the read depth *∈* {1, 5, 10, 50, 100}, the ploidy *K ∈* {2, 4, 6, 8}, the major allele frequencies (*p*_*A*_, *p*_*B*_) *∈* {(0.5, 0.5), (0.5, 0.75), (0.9, 0.9)}, and the Pears<u>o</u>n correlation *r ∈* {0, 0.5, 0.9}. When *p*_*A*_ = 0.5 and *p*_*B*_ = 0.75, *r* is constrained to be less than 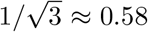, and so in this scenario the *r* = 0.9 setting was omitted. Each unique combination of parameters was replicated 200 times.

The conclusions for estimating *r*^2^ and *ρ*^2^ when *p*_*A*_ = *p*_*B*_ = 0.5 are presented in Figures 1, S5, and S6. Mean-squared error performance for estimating *D* and *D*′ when *p*_*A*_ = *p*_*B*_ = 0.5 are presented in Figures S7 and S8, respectively. Results for other scenarios are similar and are available on Figshare (https://doi.org/10.6084/m9.figshare.12765803). We see that the moment-based estimators of composite LD 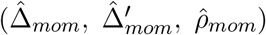 have a strong bias toward 0 until a large read-depth is attained (Figure S5). This bias makes these estimators look like they perform very well when LD is close to zero, but they consequently perform very poorly for large levels of LD. Furthermore, maximum likelihood estimation of haplotypic LD using genotype likelihoods 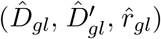 generally has better mean squared error (MSE) and bias performance in all scenarios compared to maximum likelihood estimation of haplotypic LD using posterior mode genotypes 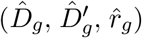 (Figures S5 and 1). Maximum likelihood estimation using the genotype likelihoods is also generally unbiased (Figure S5). It pays for this lower bias by having a larger standard error (Figure S6), but it also produces lower MSE in non-zero LD regimes (Figure 1). Using genotype likelihoods is mostly important in settings of high ploidy. All methods behave rather similarly in diploids, but the genotype likelihood approaches perform much better at higher ploidy levels in high LD regimes (Figure 1). Finally, even though these data were simulated under HWE, estimators of composite measures of LD using genotype likelihoods generally perform as well as the estimators of haplotypic LD when estimating *r*^2^ (Figure 1). The HWE assumption helps when estimating *D* and *D*′ (Figures S7 and S8).

**Figure 1:**
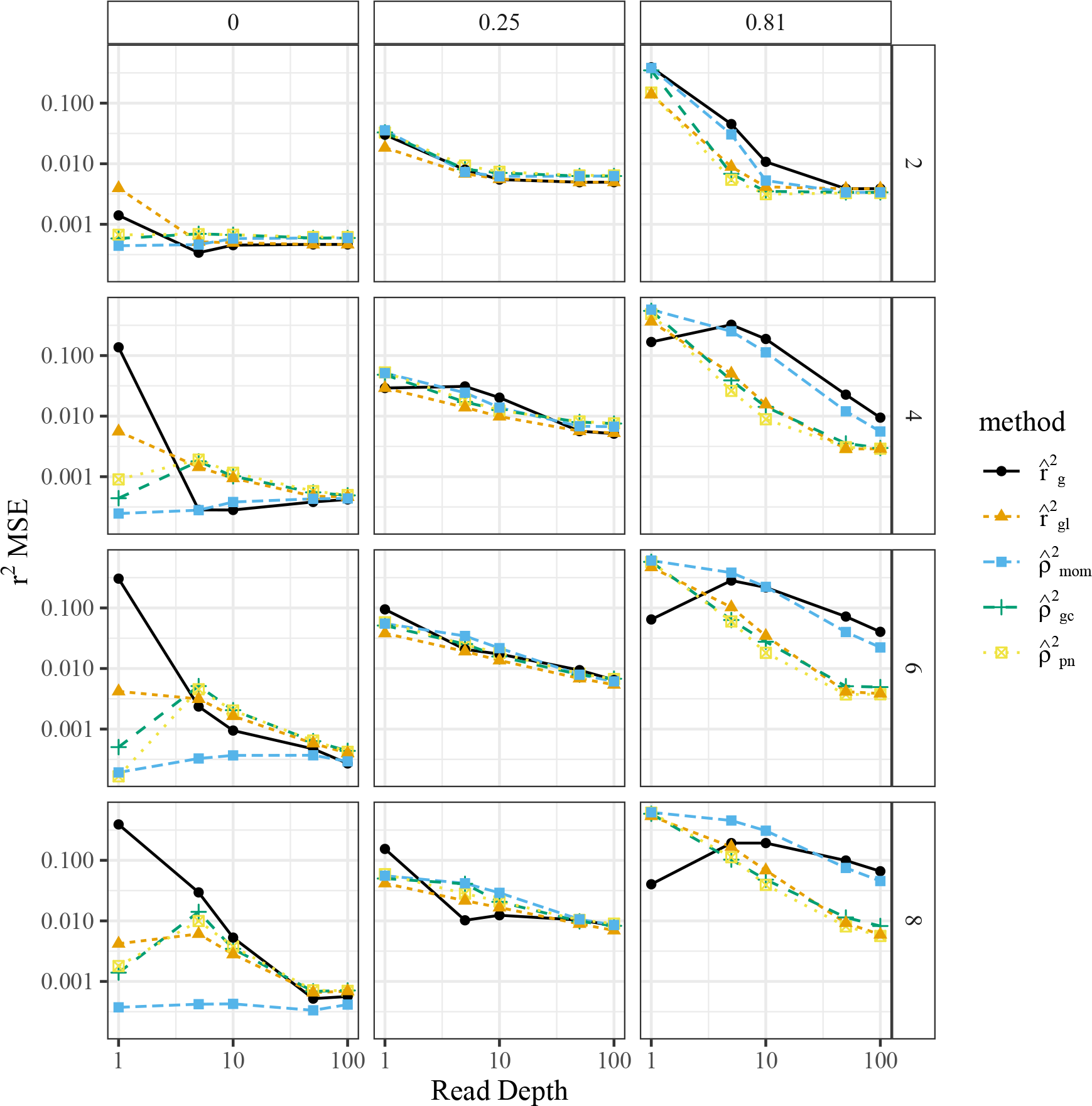
Mean-squared error of *r*^2^ estimators (*y*-axis) stratified by read-depth (*x*-axis), estimation method (color), ploidy (row-facets), and true *r*^2^ (column-facets) for the simulations from Section 3.1. Except in low-LD regimes, the MLE using genotype likelihoods has the smallest MSE. The moment-based estimator has a lower MSE in low-LD regimes because of its strong bias toward 0. Simulations were performed with *p*_*A*_ = 0.5 and *p*_*B*_ = 0.5.

Figures 2 and S9 highlight that our standard errors for the LD estimators are generally accurate except when the read-depth is 1. Computation time for each method is presented in Figure S11. We see there that ploidy is the major cause of computation time increases and that genotype likelihood methods tend to be much slower than methods that use estimated genotypes. However, all methods take less than half a second on average.

**Figure 2:**
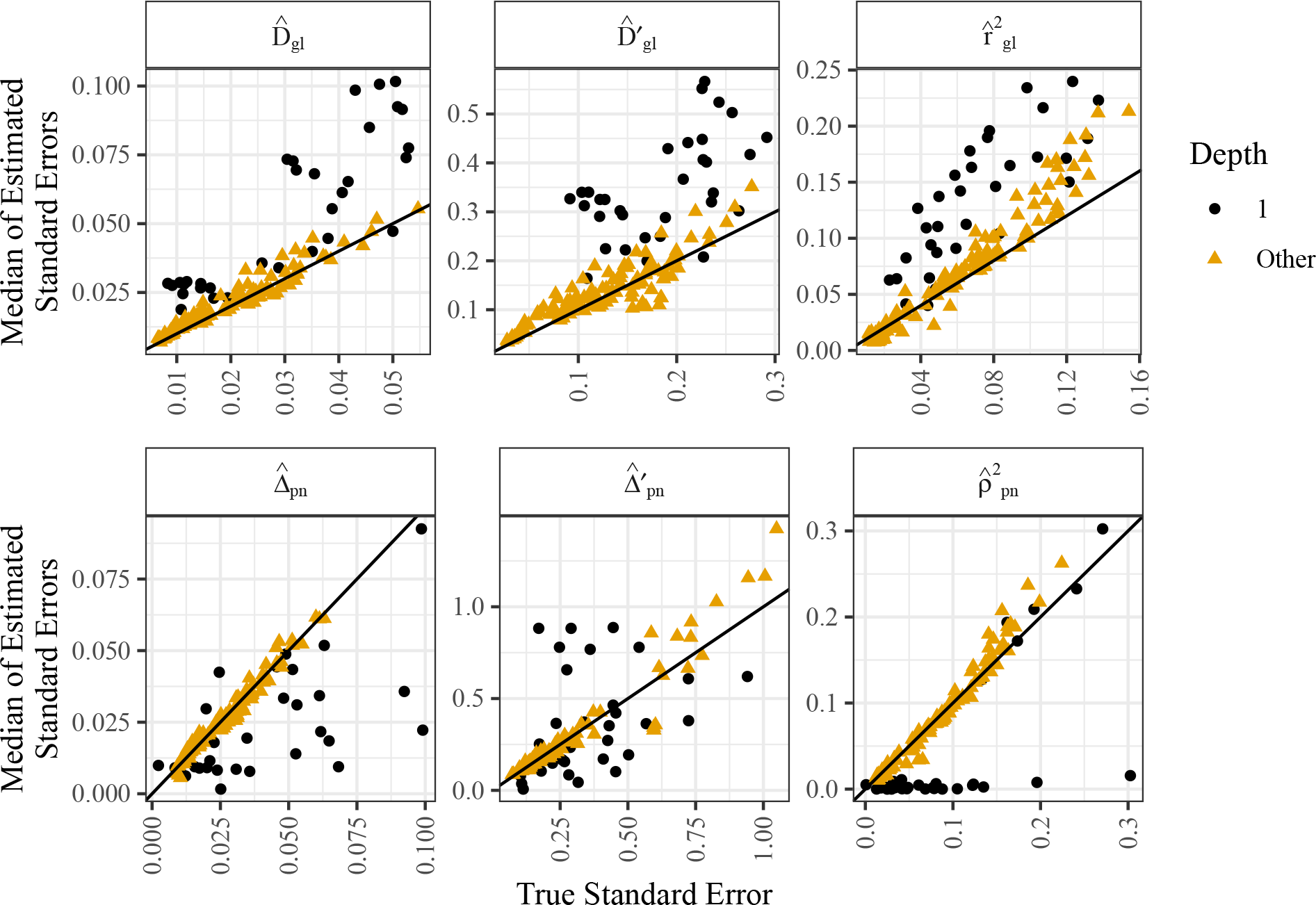
Standard errors (*x*-axis) of haplotypic (first row) and composite (second row) LD estimators when using genotype likelihoods from the simulations in Section 3.1. The *y*-axis contains the median of the estimated standard errors. Each point is a different simulation setting. The line is the *y* = *x* line and points above that line indicate that the estimated standard errors are typically larger than the true standard errors. Standard errors are reasonably unbiased except when the read-depth is 1 (color and shape).

### 3.2 Additional simulations under deviations from HWE

Additional simulation results, evaluating the performance of the various LD estimators when HWE is violated, are presented in Sections S11 and S12. The simulations in Section S11 generate genotypes under the proportional bivariate normal distribution in order to observe general deviations from HWE. The simulations in Section S12 generate genotypes using the PedigreeSim software [Voorrips and Maliepaard, 2012] under specific deviations from HWE, namely partial preferential pairing and double reduction. We see in both sections that the composite measures of LD perform better when the assumptions of HWE are not appropriate. Though the haplotypic methods are robust to instances of double reduction and moderate levels of preferential pairing.

### 3.3 LD estimates using data from *Solanum tuberosum*

We evaluated all pairwise LD estimators discussed in this paper on the genotyping-by-sequencing data from Uitdewilligen et al. [2013b]. These data come from a diversity panel of autotretraploid *Solanum tuberosum* (2*n* = 4*x* = 48). We randomly selected two regions of size 40 SNPs each, obtained their genotype estimates and genotype likelihoods using `updog` [Gerard et al., 2018, Gerard and Ferrão, 2019], and filtered out SNPs that were mostly monoallelic (had one dosage with an estimated prior probability of more than 0.9). These genotypes and genotype likelihoods were then used to estimate the LD between all SNPs within and between both regions. Though these regions were arbitrarily chosen, the results we present below are robust to the region selection and the reader is encouraged to change the random seed in our reproducible scripts on Figshare (https://doi.org/10.6084/m9.figshare.12765803).

A pairs plot comparing the estimates of *r*^2^ (or *ρ*^2^) between pairs of SNPs within each region is presented in Figure 3, and between pairs of SNPs between regions in Figure 4. We would expect well-performing methods to (i) have high LD estimates within each region and (ii) low LD estimates between regions. The estimators that do not use genotype likelihoods,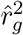 and 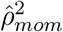, generally produce the smallest LD estimates within each region (Figure 3), indicating that using genotype likelihoods generally produced better results. This is without sacrificing LD estimation between regions, where all methods generally produced small LD estimates (Figure 4). Generally, the composite methods produced larger LD estimates within regions. This might be because these data come from a diversity panel, and so they are likely not in HWE (Figure S32). Heatmaps of LD estimates of *r*^2^ or *ρ*^2^ are presented in Figures S26–S30.

**Figure 3:**
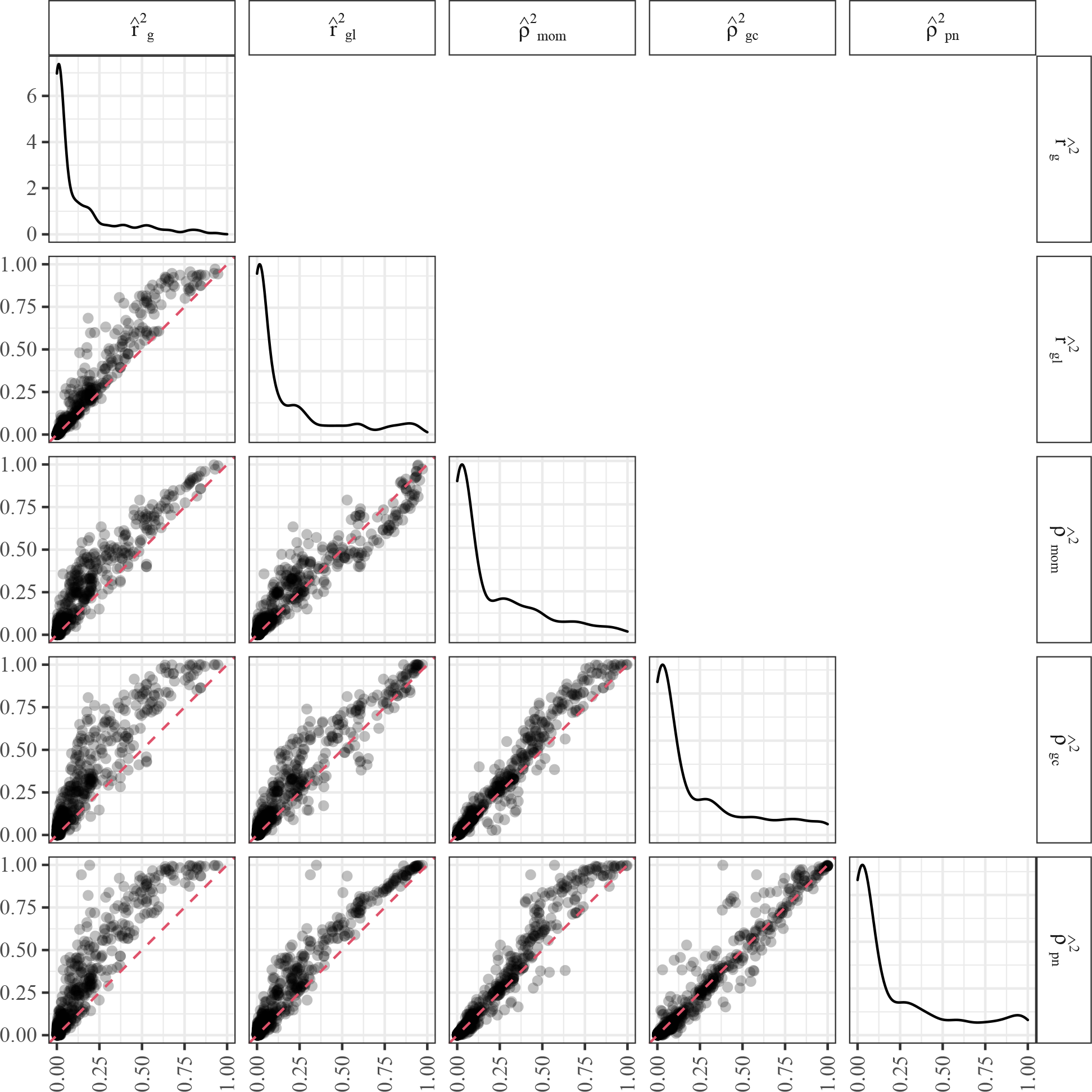
Pairs plot of within-region LD estimates of SNPs obtained from two regions from the *Solanum tuberosum* data of Uitdewilligen et al. [2013b]. Each point is a pair of SNPs. The red dashed line is the *y* = *x* line, and any points above this line indicates the estimator on the *y*-axis produced a larger LD estimate than the estimator on the *x*-axis. We would generally expect better-performing estimators to produce larger LD estimates within each region. The diagonal plots are density plots for LD estimates.

**Figure 4:**
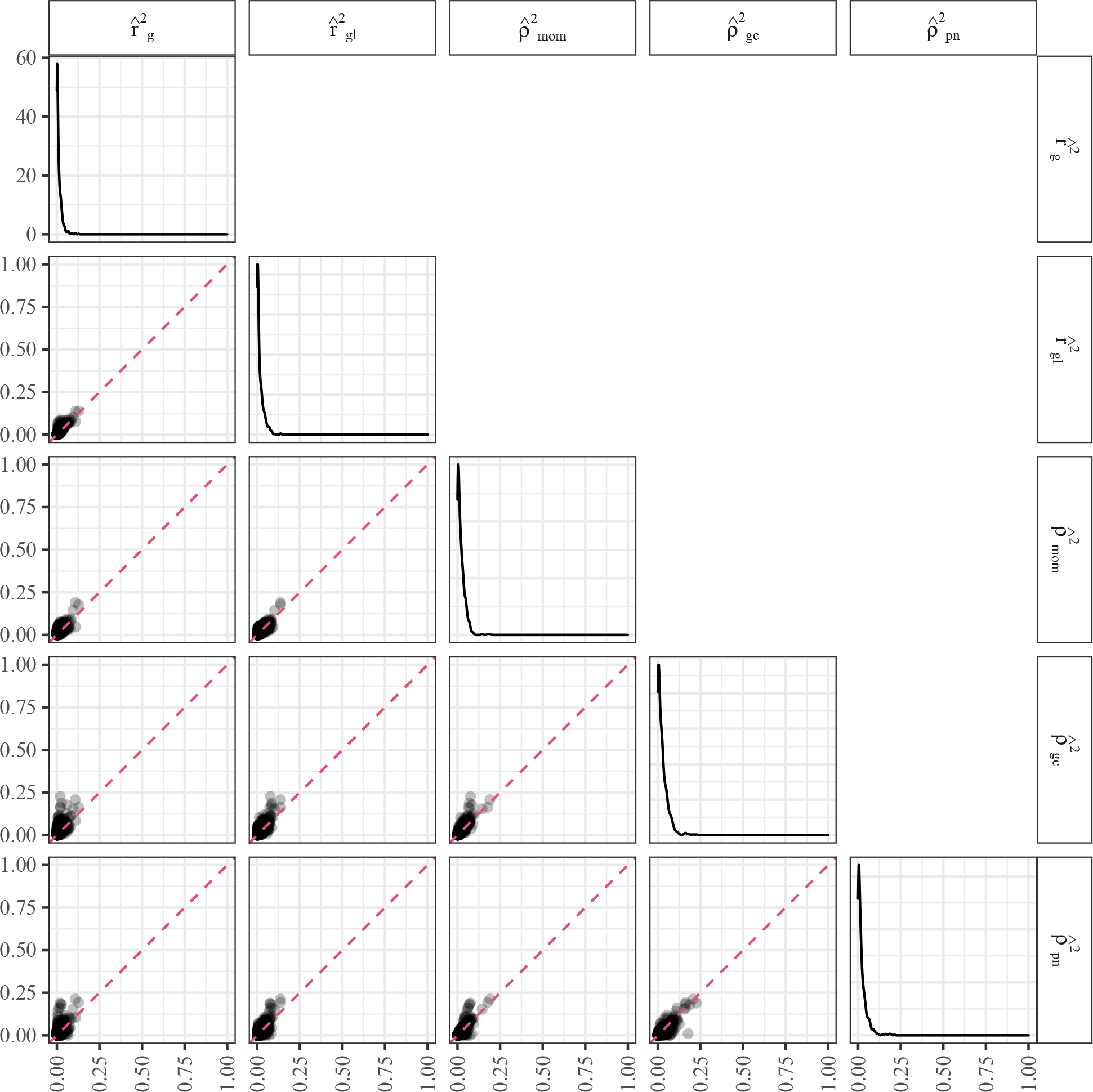
Pairs plot of between-region LD estimates of SNPs obtained from two regions from the *Solanum tuberosum* data of Uitdewilligen et al. [2013b]. Each point is a pair of SNPs. The red dashed line is the *y* = *x* line, and any points above this line indicates the estimator on the *y*-axis produced a larger LD estimate than the estimator on the *x*-axis. The diagonal plots are density plots for LD estimates. We would generally expect most LD estimates to be near 0 between regions.

Additional empirical evaluations using a dataset with a lower read-depth [McAllister and Miller, 2016] are presented in Section S13.

## 4 Discussion

In this manuscript, we reviewed haplotypic measures of LD and then derived generalizations of Burrows’ composite measures of LD to polyploids. For the composite generalizations of Lewontin‘s *D*′, this involved deriving novel bounds on the covariance between genotypes. We provided a collection of methods to estimate both haplotypic LD and composite LD in the presence of genotype uncertainty by directly using genotype likelihoods. For composite LD, this involved developing a novel class of distributions over the genotypes. We validated our methods both in simulations and on real data.

We found that not accounting for genotype uncertainty results in underestimating LD, particularly between pairs of SNPs in high LD. Our LD estimation procedures, using genotype likelihoods to account for this genotype uncertainty, help reduce this bias. This has the potential to impact many downstream applications of LD. For example, pairwise LD is often used to estimate effective population size [Hill, 1981, Waples, 2006], and attenuation bias of LD estimates would result in over-estimating the effective population size. Using our LD estimators should correct this bias. Pairwise LD estimates are also often used to estimate the extent LD decays with genetic distance. This has applications in determining the density of markers necessary to successfully map a quantitative trait locus [Vos et al., 2017]. Attenuation bias of LD estimates would result in believing LD decays faster than in reality, and thus result in overestimating the number of required markers. Using our LD estimators should provide more accurate LD decay estimates. Another example comes from new approaches that use summary statistics to run multiple SNP GWAS [Zhu and Stephens, 2017]. These create likelihoods that critically depend on accurate estimates of LD, and using poor LD estimates would result in decreased performance.

Our estimation procedures contain the following assumptions. First, we assume that the genotype likelihoods are well-calibrated. That is, misspecification of the genotype likelihood would result in bias in the LD estimates that use these genotype likelihoods. Accurately calibrating genotype likelihoods is a non-trivial problem in polyploids due to the many features of polyploid sequencing data that make modeling difficult [Gerard et al., 2018]. Such calibration is particularly difficult at very low sequencing depth, which might explain why most methods behave similarly on the data from McAllister and Miller [2016] (Section S13). This is an ongoing research problem and improved calibration will likely improve our LD estimates. Our model further assumes that the probability of the data given the genotypes at both loci are independent. This is also an assumption of Li [2011], Maruki and Lynch [2014], Bilton et al. [2018], Fox et al. [2019] and, as mentioned by Bilton et al. [2018] could be violated if SNPs were close enough to each other to be located on the same sequencing reads. As long as SNPs are on separate reads, this assumption is mostly reasonable. We also assume that individuals were sampled randomly from the population of interest. Bias in the sampling scheme could result in biased LD estimates, which might be influenced by the structure of any subpopulation being overly represented in the sample.

All of our haplotypic LD estimators assume that we are working with a population of autopolyploid individuals under HWE that exhibit polysomic inheritance and bivalent non-preferential pairing. Though we note that these methods are rather robust to the bivalent pairing assumption (Section S12). However, all of our composite LD estimators are applicable more generally to violations in both the random mating and the mode of inheritance assumptions. In particular, our composite estimators are applicable for allopolyploids, for autopolyploids that exhibit double reduction, and for polyploids have have intermediate inheritance modes, variously labeled segmental allopolyploidy [Stebbins, 1947], partial preferential pairing [Wu et al., 1992], incomplete polysomy [Guimarães et al., 1997], heterosomy [Roux and Pannell, 2015], or mixosomy [Soltis et al., 2016]. In Section S7, we provide an interpretable decomposition of Δ in the case of allopolyploids. For general violations of HWE, for intermediate forms of polyploidy, or for cases of extensive double reduction, the interpretation is more difficult. However, because these composite measures of LD are functions of usual statistical measures of association (the correlation and covariance), one should interpret these composite measures as one would usually interpret such statistical measures of association: as describing the strength of the linear association between dosages at two loci.

We briefly used covariance shrinkage in this manuscript (Section S13). Shrinkage is a statistical technique to reduce variability and improve estimation performance by adjusting estimates toward some known or estimated value. Typically, the direction and strength of the shrinkage is derived by assuming a common distribution of the parameters, where the estimation of said distribution is weighted inversely to the variance of the estimates. And so instances of shrinkage are sometimes termed “borrowing strength” because low variance estimators provide information on how much to shrink high variance estimators to improve overall performance. Covariance shrinkage has a long history, dating back at least to James and Stein [1961] and has shown great promise in improving covariance estimates in high dimensional regimes, including in genomic studies [Schäfer and Strimmer, 2005]. Yet, most LD estimators do not use any form of regularization or shrinkage (with the exception of Wen and Stephens [2010]). An off-the-shelf approach that we have used in this manuscript is “adaptive shrinkage” on the Fisher-*z* transformed correlation matrix [Stephens, 2016, Dey and Stephens, 2018] (Figures S38–S41). This shrinkage estimator is not purpose-built for LD estimation, and so there are many avenues for improvement (e.g., by accounting for known physical locations of SNPs). Research on LD shrinkage estimation is now more accessible with the accurate standard errors that we derived in this manuscript.

## Acknowledgments

We would like to thank Dr. Y. Samuel Wang, at the University of Chicago, for providing a simpler proof to Theorem S1. Our original approach with duality theory was much too cumbersome. We would also like to thank Dr. Luís Felipe Ventorim Ferrão, at the University of Florida, for his insightful comments on a draft of this manuscript.

## Data accessibility

Supplementary file S1 contains additional theoretical considerations, derivations, simulations, and supplementary figures. All methods discussed in this manuscript are implemented in the ldsep package, available on the Comprehensive R Archive Network (https://cran.r-project.org/package=ldsep) under a GPL-3 license. Scripts to reproduce the results of this research are available on Figshare (https://doi.org/10.6084/m9.figshare.12765803). All datasets used in this manuscript are publicly available [Uitdewilligen et al., 2013a, McAllister and Miller, 2017] and may be downloaded from:

- https://doi.org/10.5061/dryad.05qs7
- https://doi.org/10.1371/journal.pone.0062355.s004
- https://doi.org/10.1371/journal.pone.0062355.s007
- https://doi.org/10.1371/journal.pone.0062355.s009
- https://doi.org/10.1371/journal.pone.0062355.s010

## Author contributions

David Gerard developed the methodology, built the software, implemented the study, and wrote the manuscript.

## Notes

https://cran.r-project.org/package=ldsep

https://doi.org/10.6084/m9.figshare.12765803

### Competing Interest Statement

The authors have declared no competing interest.

### Summary of Updates

(i) The terminology "gametic LD" has been changed to "haplotypic LD". A discussion of this choice is presented at the start of Section 2.1. (ii) The Discussion has been improved to include more applied aspects of the methodology. Specifically, we now discuss the applications, assumptions, and interpretations of the various LD estimators introduced in this manuscript. (iii) We have included new simulations to explore the effects of interpretable deviations from Hardy-Weinberg equilibrium, namely preferential pairing and double reduction. These are provided in Section S12 of the Supplementary Material.

## References

N. A. Baird, P. D. Etter, T. S. Atwood, M. C. Currey, A. L. Shiver, Z. A. Lewis, E. U. Selker, W. A. Cresko, and E. A. Johnson. Rapid SNP discovery and genetic mapping using sequenced RAD markers. PLOS ONE, 3(10):1–7, 10 2008. doi: 10.1371/journal.pone.0003376.

M. S. Barker, N. Arrigo, A. E. Baniaga, Z. Li, and D. A. Levin. On the relative abundance of autopolyploids and allopolyploids. New Phytologist, 210(2):391–398, 2016. doi: 10.1111/nph.13698.

F. Z. Barreto, J. R. B. F. Rosa, T. W. A. Balsalobre, M. M. Pastina, R. R. Silva, H. P. Hoffmann, A. P. de Souza, A. A. F. Garcia, and M. S. Carneiro. A genome-wide association study identified loci for yield component traits in sugarcane (saccharum spp.). PLOS ONE, 14(7):1–22, 07 2019. doi: 10.1371/journal.pone.0219843.

C. Benner, A. S. Havulinna, M.-R. Järvelin, V. Salomaa, S. Ripatti, and M. Pirinen. Prospects of fine-mapping trait-associated genomic regions by using summary statistics from genome-wide association studies. The American Journal of Human Genetics, 101(4):539–551, 2017. ISSN 0002-9297. doi: 10.1016/j.ajhg.2017.08.012.

M. Berkelaar, K. Eikland, and P. Notebaert. lp solve 5.5.2.5, open source (mixed-integer) linear programming system. Software, 2016. URL http://lpsolve.sourceforge.net/5.5/. xLast Accessed July, 6 2020.

T. P. Bilton, J. C. McEwan, S. M. Clarke, R. Brauning, T. C. van Stijn, S. J. Rowe, and K. G. Dodds.Linkage disequilibrium estimation in low coverage high-throughput sequencing data. Genetics, 209(2):389–400, 2018. ISSN 0016-6731. doi: 10.1534/genetics.118.300831.

B. Björn, M. J. Paulo, K. Kowitwanich, M. Sengers, R. G. Visser, H. J. van Eck, and F. A. van Eeuwijk. Population structure and linkage disequilibrium unravelled in tetraploid potato. Theoretical and Applied Genetics, 121(6):1151–1170, 2010. doi: 10.1007/s00122-010-1379-5.

P. D. Blischak, L. S. Kubatko, and A. D. Wolfe. SNP genotyping and parameter estimation in polyploids using low-coverage sequencing data. Bioinformatics, 34(3):407–415, 2018. doi: 10.1093/bioinformatics/btx587.

P. J. Bradbury, Z. Zhang, D. E. Kroon, T. M. Casstevens, Y. Ramdoss, and E. S. Buckler. TASSEL: software for association mapping of complex traits in diverse samples. Bioinformatics, 23(19): 2633–2635, 06 2007. ISSN 1367-4803. doi: 10.1093/bioinformatics/btm308.

A. Brown. Sample sizes required to detect linkage disequilibrium between two or three loci. Theoretical Population Biology, 8(2):184 –201, 1975. ISSN 0040-5809. doi: 10.1016/0040-5809(75)90031-3.

S. R. Browning and B. L. Browning. Rapid and accurate haplotype phasing and missing-data inference for whole-genome association studies by use of localized haplotype clustering. The American Journal of Human Genetics, 81(5):1084–1097, 2007. doi: 10.1086/521987.

L. V. Clark, A. E. Lipka, and E. J. Sacks. polyRAD: Genotype calling with uncertainty from sequencing data in polyploids and diploids. G3: Genes, Genomes, Genetics, 9(3):663–673, 2019. doi: 10.1534/g3.118.200913.

C. C. Cockerham and B. S. Weir. Digenic descent measures for finite populations. Genetical Research, 30(2):121–147, 1977. doi: 10.1017/S0016672300017547.

de Bem Oliveira, M. F. R. Resende L. F. V. Ferrão, R. R. Amadeu, J. B. Endelman, M. Kirst, A. S. G. Coelho, and P. R. Muñoz. Genomic prediction of autotetraploids; influence of relationship matrices, allele dosage, and continuous genotyping calls in phenotype prediction. G3: Genes, Genomes, Genetics, 9(4):1189–1198, 2019. doi: 10.1534/g3.119.400059.

L. A. de C. Lara, M. F. Santos, L. Jank, L. Chiari, M. d. M. Vilela, R. R. Amadeu, J. P. R. dos Santos, G. d. S. Pereira, Z.-B. Zeng, and A. A. F. Garcia. Genomic selection with allele dosage in Panicum maximum jacq. G3: Genes, Genomes, Genetics, 9(8):2463–2475, 2019. doi: 10.1534/g3.118.200986.

P. Dempster, N. M. Laird, and D. B. Rubin. Maximum likelihood from incomplete data via the EM algorithm. Journal of the Royal Statistical Society: Series B (Methodological), 39(1):1–22, 1977. doi: 10.1111/j.2517-6161.1977.tb01600.x.

Devlin and N. Risch. A comparison of linkage disequilibrium measures for fine-scale mapping. Genomics, 29(2):311–322, 1995. doi: 10.1006/geno.1995.9003.

K. K. Dey and M. Stephens. CorShrink: Empirical Bayes shrinkage estimation of correlations, with applications. bioRxiv, 2018. doi: 10.1101/368316.

R. J. Elshire, J. C. Glaubitz, Q. Sun, J. A. Poland, K. Kawamoto, E. S. Buckler, and S. E. Mitchell. A robust, simple genotyping-by-sequencing (GBS) approach for high diversity species. PLOS ONE, 6(5):1–10, 05 2011. doi: 10.1371/journal.pone.0019379.

J.-B. Fan, A. Oliphant, R. Shen, B. G. Kermani, F. García, K. L. Gunderson, M. S. T. Hansen, F. Steemers, S. L. Butler, P. Deloukas, L. Galver, S. Hunt, C. McBride, M. Bibikova, T. Rubano, Chen E. Wickham, D. Doucet, W. Chang, D. Campbell, B. Zhang, S. Kruglyak, D. Bentley, J. Haas, P. Rigault, L. Zhou, J. R. Stuelpnagel, and M. S. Chee. Highly parallel SNP genotyping. Cold Spring Harbor Symposia on Quantitative Biology, 68:69–78, 2003. doi: 10.1101/sqb.2003.68.69.

K.-H. Farh, A. Marson, J. Zhu, M. Kleinewietfeld, W. J. Housley, S. Beik, N. Shoresh, H. Whitton, R. J. Ryan, A. A. Shishkin, M. Hatan, M. J. Carrasco-Alfonso, D. Mayer, C. J. Luckey, N. A. Patsopoulos, P. L. De Jager, V. K. Kuchroo, C. B. Epstein, M. J. Daly, D. A. Hafler, and B. E. Bernstein. Genetic and epigenetic fine mapping of causal autoimmune disease variants. Nature, 518(7539):337–343, 2015. doi: 10.1038/nature13835.

F. V. Ferrão, T. S. Johnson, J. Benevenuto, P. P. Edger, T. A. Colquhoun, and P. R. Munoz. Genome-wide association of volatiles reveals candidate loci for blueberry flavor. New Phytologist, 226(6):1725–1737, 2020. doi: 10.1111/nph.16459.

E. A. Fox, A. E. Wright, M. Fumagalli, and F. G. Vieira. ngsLD: evaluating linkage disequilibrium using genotype likelihoods. Bioinformatics, 35(19):3855–3856, 03 2019. ISSN 1367-4803. doi:10.1093/bioinformatics/btz200.

D. Gerard and L. F. V. Ferrão. Priors for genotyping polyploids. Bioinformatics, 36(6):1795–1800, 11 2019. ISSN 1367-4803. doi: 10.1093/bioinformatics/btz852. bioRxiv: 751784.

D. Gerard, L. F. V. Ferrão, A. A. F. Garcia, and M. Stephens. Genotyping polyploids from messy sequencing data. Genetics, 210(3):789–807, 2018. ISSN 0016-6731. doi: 10.1534/ge-netics.118.301468.

A. G. Griffiths, R. Moraga, M. Tausen, V. Gupta, T. P. Bilton, M. A. Campbell, R. Ashby, I. Nagy, A. Khan, A. Larking, C. Anderson, B. Franzmayr, K. Hancock, A. Scott, N. W. Ellison, M. P. Cox, T. Asp, T. Mailund, M. H. Schierup, and S. U. Andersen. Breaking free: The genomics of allopolyploidy-facilitated niche expansion in white clover. The Plant Cell, 31(7):1466–1487, 2019. ISSN 1040-4651. doi: 10.1105/tpc.18.00606.

C. T. Guimarães, G. R. Sills, and B. W. S. Sobral. Comparative mapping of Andropogoneae: Saccharum L. (sugarcane) and its relation to sorghum and maize. Proceedings of the National Academy of Sciences, 94(26):14261–14266, 1997. ISSN 0027-8424. doi: 10.1073/pnas.94.26.14261.

A. Gur, G. Tzuri, A. Meir, U. Sa’ar, V. Portnoy, N. Katzir, A. A. Schaffer, L. Li, J. Burger, and Y. Tadmor. Genome-wide linkage-disequilibrium mapping to the candidate gene level in melon (Cucumis melo). Scientific reports, 7(1):1–13, 2017. doi: 10.1038/s41598-017-09987-4.

D. C. Hamilton and D. E. C. Cole. Standardizing a composite measure of linkage disequilibrium. Annals of Human Genetics, 68(3):234–239, 2004. doi: 10.1046/j.1529-8817.2004.00056.x.

P. Hedrick, S. Jain, and L. Holden. Multilocus systems in evolution. In M. K. Hecht, W. C. Steere, and B. Wallace, editors, Evolutionary Biology, Bolume 11, pages 101–184. Springer, 1978. doi:10.1007/978-1-4615-6956-5_3.

P. W. Hedrick. Gametic disequilibrium measures: Proceed with caution. Genetics, 117(2):331–341, 1987. ISSN 0016-6731. URL https://www.genetics.org/content/117/2/331.

W. Hill and A. Robertson. Linkage disequilibrium in finite populations. Theoretical and applied genetics, 38(6):226–231, 1968. doi: 10.1007/BF01245622.

W. G. Hill. Estimation of linkage disequilibrium in randomly mating populations. Heredity, 33(2): 229, 1974. doi: 10.1038/hdy.1974.89.

W. G. Hill. Estimation of effective population size from data on linkage disequilibrium. Genetical Research, 38(3):209–216, 1981. doi: 10.1017/S0016672300020553.

K. Huang, D. W. Dunn, K. Ritland, and B. Li. polygene: Population genetics analyses for autopolyploids based on allelic phenotypes. Methods in Ecology and Evolution, 11(3):448–456, 2020. doi:10.1111/2041-210X.13338.

T.-Y. J. Hui and A. Burt. Estimating linkage disequilibrium from genotypes under Hardy-Weinberg equilibrium. BMC genetics, 21(1):1–11, 2020. doi: 10.1186/s12863-020-0818-9.

W. James and C. Stein. Estimation with quadratic loss. In Proceedings of the Fourth Berkeley Symposium on Mathematical Statistics and Probability, Volume 1: Contributions to the Theory of Statistics, pages 361–379,

Berkeley, Calif., 1961. University of California Press. URL https://projecteuclid.org/euclid.bsmsp/1200512173.

L. B. Jorde. Linkage disequilibrium as a gene-mapping tool. American journal of human genetics, 56(1):11, 1995.

B. Julier. A program to test linkage disequilibrium between loci in autotetraploid species. Molecular ecology resources, 9(3):746–748, 2009. doi: 10.1111/j.1755-0998.2009.02530.x.

R. Lewontin. The interaction of selection and linkage. i. general considerations; heterotic models. Genetics, 49(1):49, 1964. URL https://www.genetics.org/content/49/1/49.

R. C. Lewontin. On measures of gametic disequilibrium. Genetics, 120(3):849–852, 1988. ISSN 0016-6731. URL https://www.genetics.org/content/120/3/849.

R. C. Lewontin and K.-i. Kojima. The evolutionary dynamics of complex polymorphisms. Evolution, 14(4):458–472, 1960. doi: 10.1111/j.1558-5646.1960.tb03113.x.

H. Li. A statistical framework for SNP calling, mutation discovery, association mapping and population genetical parameter estimation from sequencing data. Bioinformatics, 27(21):2987, 2011. doi: 10.1093/bioinformatics/btr509.

Y. Li, C. J. Willer, J. Ding, P. Scheet, and G. R. Abecasis. MaCH: using sequence and genotype data to estimate haplotypes and unobserved genotypes. Genetic epidemiology, 34(8):816–834, 2010. doi: 10.1002/gepi.20533.

F. Lu, A. E. Lipka, J. Glaubitz, R. Elshire, J. H. Cherney, M. D. Casler, E. S. Buckler, and D. E. Costich. Switchgrass genomic diversity, ploidy, and evolution: novel insights from a network-based SNP discovery protocol. PLOS Genetics, 9(1):1–14, 01 2013. doi: 10.1371/journal.pgen.1003215.

I. Mackay and W. Powell. Methods for linkage disequilibrium mapping in crops. Trends in Plant Science, 12(2):57–63, 2007. ISSN 1360-1385. doi: 10.1016/j.tplants.2006.12.001.

T. Maruki and M. Lynch. Genome-wide estimation of linkage disequilibrium from population-level high-throughput sequencing data. Genetics, 197(4):1303–1313, 2014. ISSN 0016-6731. doi:10.1534/genetics.114.165514.

T. Maruki and M. Lynch. Genotype calling from population-genomic sequencing data. G3: Genes, Genomes, Genetics, 7(5):1393–1404, 2017. doi: 10.1534/g3.117.039008.

F. I. Matias, K. G. Xavier Meireles, S. T. Nagamatsu, S. C. Lima Barrios, C. Borges do Valle, M. F. Carazzolle, R. Fritsche-Neto, and J. B. Endelman. Expected genotype quality and diploidized marker data from genotyping-by-sequencing of Urochloa spp. tetraploids. The Plant Genome, 12 (3):190002, 2019. doi: 10.3835/plantgenome2019.01.0002.

C. A. McAllister and A. J. Miller. Single nucleotide polymorphism discovery via genotyping by sequencing to assess population genetic structure and recurrent polyploidization in Andropogon gerardii. American Journal of Botany, 103(7):1314–1325, 2016. doi: 10.3732/ajb.1600146.

C. A. McAllister and A. J. Miller. Data from: Single nucleotide polymorphism discovery via genotyping by sequencing to assess population genetic structure and recurrent polyploidization in Andropogon gerardii, 2017. URL https://doi.org/10.5061/dryad.05qs7. Dataset.

A. McKenna, M. Hanna, E. Banks, A. Sivachenko, K. Cibulskis, A. Kernytsky, K. Garimella, D. Altshuler, S. Gabriel, M. Daly, and M. DePristo. The genome analysis toolkit: a MapReduce framework for analyzing next-generation DNA sequencing data. Genome research, 20(9):1297–1303, 2010. doi: 10.1101/gr.107524.110.

M. Mollinari and A. A. F. Garcia. Linkage analysis and haplotype phasing in experimental autopolyploid populations with high ploidy level using hidden markov models. G3: Genes, Genomes, Genetics, 9(10):3297–3314, 2019. doi: 10.1534/g3.119.400378.

M. Mollinari and O. Serang. Quantitative SNP genotyping of polyploids with MassARRAY and other platforms. In J. Batley, editor, Plant Genotyping: Methods and Protocols, pages 215–241. Springer New York, New York, NY, 2015. ISBN 978-1-4939-1966-6. doi: 10.1007/978-1-4939-1966-6_17.

J. Nocedal and S. Wright. Numerical optimization. Springer Science & Business Media, 2006. ISBN 0-387-30303-0.

P. Oeth, G. del Mistro, G. Marnellos, T. Shi, and D. van den Boom. Qualitative and quantitative genotyping using single base primer extension coupled with matrix-assisted laser desorption/ionization time-of-flight mass spectrometry (MassARRAY®). In A. Komar, editor, Single Nucleotide Polymorphisms, pages 307–343. Humana Press, 2009. ISBN 978-1-60327-411-1. doi: 10.1007/978-1-60327-411-1_20.

Y. Okada. eLD: entropy-based linkage disequilibrium index between multiallelic sites. Human genome variation, 5(1):1–3, 2018. doi: 10.1038/s41439-018-0030-x.

L.-M. Raboin, J. Pauquet, M. Butterfield, A. D’Hont, and J.-C. Glaszmann. Analysis of genome-wide linkage disequilibrium in the highly polyploid sugarcane. Theoretical and Applied Genetics, 116(5):701–714, 2008. doi: 10.1007/s00122-007-0703-1.

G. P. Ramstein, J. Evans, S. M. Kaeppler, R. B. Mitchell, K. P. Vogel, C. R. Buell, and M. D. Casler. Accuracy of genomic prediction in switchgrass (Panicum virgatum l.) improved by accounting for linkage disequilibrium. G3: Genes, Genomes, Genetics, 6(4):1049–1062, 2016. doi: 10.1534/g3.115.024950.

A. R. Rogers and C. Huff. Linkage disequilibrium between loci with unknown phase. Genetics, 182 (3):839–844, 2009. ISSN 0016-6731. doi: 10.1534/genetics.108.093153.

C. Roux and J. R. Pannell. Inferring the mode of origin of polyploid species from next-generation sequence data. Molecular Ecology, 24(5):1047–1059, 2015. doi: 10.1111/mec.13078.

P. Scheet and M. Stephens. A fast and flexible statistical model for large-scale population genotype data: applications to inferring missing genotypes and haplotypic phase. The American Journal of Human Genetics, 78(4):629–644, 2006. doi: 10.1086/502802.

C. A. Schmitz Carley, J. J. Coombs, D. S. Douches, P. C. Bethke, J. P. Palta, R. G. Novy, and J. B. Endelman. Automated tetraploid genotype calling by hierarchical clustering. Theoretical and Applied Genetics, 130(4):717–726, 2017. ISSN 1432-2242. doi: 10.1007/s00122-016-2845-5.

J. Schäfer and K. Strimmer. A shrinkage approach to large-scale covariance matrix estimation and implications for functional genomics. Statistical Applications in Genetics and Molecular Biology, 4(1), 2005. doi: https://doi.org/10.2202/1544-6115.1175.

O. Serang, M. Mollinari, and A. A. F. Garcia. Efficient exact maximum a posteriori computation for Bayesian SNP genotyping in polyploids. PLOS ONE, 7(2):1–13, 02 2012. doi: 10.1371/jour-nal.pone.0030906.

S. K. Sharma, K. MacKenzie, K. McLean, F. Dale, S. Daniels, and G. J. Bryan. Linkage disequilibrium and evaluation of genome-wide association mapping models in tetraploid potato. G3: Genes, Genomes, Genetics, 8(10):3185–3202, 2018. doi: 10.1534/g3.118.200377.

J. Shen, Z. Li, J. Chen, Z. Song, Z. Zhou, and Y. Shi. SHEsisPlus, a toolset for genetic studies on polyploid species. Scientific reports, 6:24095, 2016. doi: 10.1038/srep24095.

I. Simko, K. G. Haynes, and R. W. Jones. Assessment of linkage disequilibrium in potato genome with single nucleotide polymorphism markers. Genetics, 173(4):2237–2245, 2006. ISSN 0016-6731. doi: 10.1534/genetics.106.060905.

M. Slatkin. Linkage disequilibrium-understanding the evolutionary past and mapping the medical future. Nature Reviews Genetics, 9(6):477, 2008. doi: 10.1038/nrg2361.

D. E. Soltis, C. J. Visger, and P. S. Soltis. The polyploidy revolution then…and now: Stebbins revisited. American Journal of Botany, 101(7):1057–1078, 2014. doi: 10.3732/ajb.1400178.

D. E. Soltis, C. J. Visger, D. B. Marchant, and P. S. Soltis. Polyploidy: Pitfalls and paths to a paradigm. American Journal of Botany, 103(7):1146–1166, 2016. doi: 10.3732/ajb.1500501.

G. L. Stebbins. Types of polyploids: their classification and significance. In M. Demerec, editor, Advances in Genetics, volume 1, pages 403–429. Academic Press, 1947. doi: 10.1016/S0065-2660(08)60490-3.

M. Stephens. False discovery rates: a new deal. Biostatistics, 18(2):275–294, 10 2016. ISSN 1465-4644. doi: 10.1093/biostatistics/kxw041.

S.-Y. Su, J. White, D. J. Balding, and L. J. Coin. Inference of haplotypic phase and missing genotypes in polyploid organisms and variable copy number genomic regions. BMC bioinformatics, 9 (1):1–9, 2008. doi: 10.1186/1471-2105-9-513.

X. Sun, R. Fernando, and J. Dekkers. Contributions of linkage disequilibrium and co-segregation information to the accuracy of genomic prediction. Genetics Selection Evolution, 48(1):77, 2016. doi: 10.1186/s12711-016-0255-4.

J. A. Sved and W. G. Hill. One hundred years of linkage disequilibrium. Genetics, 209(3):629–636, 2018. ISSN 0016-6731. doi: 10.1534/genetics.118.300642.

K. Swarts, H. Li, J. A. R. Navarro, D. An, M. C. Romay, S. Hearne, C. Acharya, J. Glaubitz, S. E. Mitchell, R. J. Elshire, E. S. Buckler, and P. J. Bradbury. Novel methods to optimize genotypic imputation for low-coverage, next-generation sequence data in crop plants. The Plant Genome, 7(3):1–12, 2014. doi: 10.3835/plantgenome2014.05.0023.

G. Thomson and R. M. Single. Conditional asymmetric linkage disequilibrium (ALD): Extending the biallelic r^2^ measure. Genetics, 198(1):321–331, 2014. ISSN 0016-6731. doi: 10.1534/genet-ics.114.165266.

J. A. Udall and J. F. Wendel. Polyploidy and crop improvement. Crop Science, 46(Supplement 1): S–3, 2006. doi: 10.2135/cropsci2006.07.0489tpg.

J. G. A. M. L. Uitdewilligen, A.-M. A. Wolters, B. B. D’hoop, T. J. A. Borm, R. G. F. Visser, and H. J. van Eck. Data from: A next-generation sequencing method for genotyping-by-sequencing of highly heterozygous autotetraploid potato, 2013a. Dataset, doi:10.1371/journal.pone.0062355.s004, doi:10.1371/journal.pone.0062355.s007, doi:10.1371/journal.pone.0062355.s009, doi:10.1371/journal.pone.0062355.s010.

J. G. A. M. L. Uitdewilligen, A.-M. A. Wolters, B. B. D’hoop, T. J. A. Borm, R. G. F. Visser, and H. J. van Eck. A next-generation sequencing method for genotyping-by-sequencing of highly heterozygous autotetraploid potato. PLOS ONE, 8(5):1–14, 05 2013b. doi: 10.1371/jour-nal.pone.0062355.

M. Van Wyngaarden, P. V. R. Snelgrove, C. DiBacco, L. C. Hamilton, N. Rodríguez-Ezpeleta, N. W. Jeffery, R. R. E. Stanley, and I. R. Bradbury. Identifying patterns of dispersal, connectivity and selection in the sea scallop, Placopecten magellanicus, using RADseq-derived SNPs. Evolutionary Applications, 10(1):102–117, 2017. doi: 10.1111/eva.12432.

R. E. Voorrips and C. A. Maliepaard. The simulation of meiosis in diploid and tetraploid organisms using various genetic models. BMC Bioinformatics, 13(1):248, Sep 2012. ISSN 1471-2105. doi:10.1186/1471-2105-13-248.

R. E. Voorrips, G. Gort, and B. Vosman. Genotype calling in tetraploid species from bi-allelic marker data using mixture models. BMC Bioinformatics, 12(1):172, 2011. ISSN 1471-2105. doi:10.1186/1471-2105-12-172.

P. G. Vos, M. J. Paulo, R. E. Voorrips, R. G. Visser, H. J. van Eck, and F. A. van Eeuwijk. Evaluation of LD decay and various LD-decay estimators in simulated and SNP-array data of tetraploid potato. Theoretical and Applied Genetics, 130(1):123–135, 2017. doi: 10.1007/s00122-016-2798-8.

R. S. Waples. A bias correction for estimates of effective population size based on linkage disequilibrium at unlinked gene loci. Conservation Genetics, 7(2):167, 2006. doi: 10.1007/s10592-005-9100-y.

B. Weir. Genetic data analysis II. Sinauer Associates, Inc., 1996. ISBN 0-87893-902-4.

B. Weir. Linkage disequilibrium and association mapping. Annual Review of Genomics and Human Genetics, 9(1):129–142, 2008. doi: 10.1146/annurev.genom.9.081307.164347. PMID: 18505378.

B. Weir and C. C. Cockerham. Estimation of linkage disequilibrium in randomly mating populations. Heredity, 42(1):105–111, 1979. doi: 10.1038/hdy.1979.10.

B. S. Weir. Inferences about linkage disequilibrium. Biometrics, pages 235–254, 1979. doi:10.2307/2529947.

X. Wen and M. Stephens. Using linear predictors to impute allele frequencies from summary or pooled genotype data. The annals of applied statistics, 4(3):1158–1182, 2010. ISSN 1932-6157. doi: 10.1214/10-aoas338.

Y. C. J. Wientjes, R. F. Veerkamp, and M. P. L. Calus. The effect of linkage disequilibrium and family relationships on the reliability of genomic prediction. Genetics, 193(2):621–631, 2013. ISSN 0016-6731. doi: 10.1534/genetics.112.146290.

K. Wu, W. Burnquist, M. Sorrells, T. Tew, P. Moore, and S. Tanksley. The detection and estimation of linkage in polyploids using single-dose restriction fragments. Theoretical and Applied Genetics, 83(3):294–300, 1992. doi: 10.1007/BF00224274.

M. Xiong and S.-W. Guo. Fine-scale genetic mapping based on linkage disequilibrium: Theory and applications. The American Journal of Human Genetics, 60(6):1513–1531, 1997. ISSN 0002-9297. doi: 10.1086/515475.

J. Yang, T. Ferreira, A. P. Morris, S. E. Medland, G. I. of ANthropometric Traits (GIANT) Consortium, D. G. Replication, M. analysis (DIAGRAM) Consortium, P. A. Madden, A. C. Heath, N. G. Martin, G. W. Montgomery, M. N. Weedon, R. J. Loos, T. M. Frayling, M. I. McCarthy, J. N. Hirschhorn, M. E. Goddard, and P. M. Visscher. Conditional and joint multiple-SNP analysis of GWAS summary statistics identifies additional variants influencing complex traits. Nature genetics, 44(4):369, 2012. doi: 10.1038/ng.2213.

D. V. Zaykin. Bounds and normalization of the composite linkage disequilibrium coefficient. Genetic Epidemiology, 27(3):252–257, 2004. doi: 10.1002/gepi.20015.

C. Zheng, R. E. Voorrips, J. Jansen, C. A. Hackett, J. Ho, and M. C. Bink. Probabilistic multilocus haplotype reconstruction in outcrossing tetraploids. Genetics, 203(1):119–131, 2016. doi:10.1534/genetics.115.185579.

X. Zhu and M. Stephens. Bayesian large-scale multiple regression with summary statistics from genome-wide association studies. Ann. Appl. Stat., 11(3):1561–1592, 09 2017. doi: 10.1214/17-AOAS1046.

X. Zhu, F. Xu, S. Zhao, W. Bo, L. Jiang, X. Pang, and R. Wu. Inferring the evolutionary history of outcrossing populations through computing a multiallelic linkage–linkage disequilibrium map. Methods in Ecology and Evolution, 6(11):1259–1269, 2015. doi: 10.1111/2041-210X.12428.

K. Zych, G. Gort, C. A. Maliepaard, R. C. Jansen, and R. E. Voorrips. FitTetra 2.0–improved genotype calling for tetraploids with multiple population and parental data support. BMC bioinformatics, 20(1):148, 2019. doi: 10.1186/s12859-019-2703-y.

