## Supplementary File S1 for "Pairwise Linkage Disequilibrium Estimation for Polyploids"

David Gerard

Department of Mathematics and Statistics, American University, Washington, DC, 20016, USA

**Abstract**

This document contains additional theoretical considerations, derivations, simulations, and figures to supplement the manuscript "Pairwise Linkage Disequilibrium Estimation for Polyploids".

#### S1 Derivation of Equation (8)

Under Hardy-Weinberg equilibrium, $(X_{iAB}, X_{iAb}, X_{iaB}, X_{iab})$ follows a multinomial distribution (7) with size parameter $K$ and probability parameters $\boldsymbol{p} = (p_{AB}, p_{Ab}, p_{aB}, p_{AB})$. We will denote the multinomial probability mass function by $\text{Multinom}(X_{iAB}, X_{iAb}, X_{iaB}, X_{iab}|K, \boldsymbol{p})$. Letting $G_{iA} = X_{iAB} + X_{iAb}$ and $G_{iB} = X_{iAB} + X_{iaB}$, the change of variables results in

$$Pr(G_{iA}, G_{iB}|\boldsymbol{p}) = \sum_{\substack{X_{iAB}, X_{iAb}, X_{iaB}, X_{iab} \text{ s.t.} \\ G_{iA}=X_{iAB}+X_{iAb}, \\ G_{iB}=X_{iAB}+X_{iaB}, \text{ and} \\ X_{iAB}+X_{iAb}+X_{iaB}+X_{iab}=K}} \text{Multinom}(X_{iAB}, X_{iAb}, X_{iaB}, X_{iab}|K, \boldsymbol{p}). \tag{S1}$$

Noting that

$$\begin{aligned} X_{iAb} &= G_{iA} - X_{iAB} && \text{(S2)} \\ X_{iaB} &= G_{iB} - X_{iAB} && \text{(S3)} \\ X_{iab} &= K - X_{iAb} - X_{iaB} - X_{iAB} && \text{(S4)} \\ &= K - G_{iA} - G_{iB} + X_{iAB}, && \text{(S5)} \end{aligned}$$

and then relabeling $z = X_{iAB}$, (S1) becomes

$$Pr(G_{iA}, G_{iB}|\boldsymbol{p}) = \sum_{z} \text{Multinom}(z, G_{iA} - z, G_{iB} - z, K - G_{iA} - G_{iB} + z|K, \boldsymbol{p}). \tag{S6}$$

It remains to find the limits of the summation in (S6). Since each $X$ lies between 0 and $K$ we have

$$\begin{aligned} && 0 \leq z \leq K && \text{(S7)} \\ 0 \leq G_{iA} - z \leq K \Rightarrow && G_{iA} - K \leq z \leq G_{iA} && \text{(S8)} \\ 0 \leq G_{iB} - z \leq K \Rightarrow && G_{iB} - K \leq z \leq G_{iB} && \text{(S9)} \\ 0 \leq K - G_{iA} - G_{iB} + z \leq K \Rightarrow && G_{iA} + G_{iB} - K \leq z \leq G_{iA} + G_{iB}. && \text{(S10)} \end{aligned}$$

Taking the intersection of bounds (S7)-(S10), we obtain

$$\max(0, G_{iA} + G_{iB} - K) \leq z \leq \min(G_{iA}, G_{iB}). \tag{S11}$$

Placing the bounds of (S11) in (S6) and substituting in the multinomial probability mass function yields (8).

#### S2 Optimization algorithms to estimate haplotype frequencies in autopolyploids under HWE

The following derivation generalizes from diploids to polyploids the EM algorithm described in Li [2011] and later used in Fox et al. [2019]. However, unlike the algorithm in Li [2011] that uses the haplotypes as the latent variable, we use the *number* of each haplotype as the latent variable. This simplifies the EM algorithm derivation for polyploids and significantly reduces the number of summands each iteration from $4^K$ to $\binom{K+3}{K}$. For example, for an octoploid species like strawberry ($K = 8$), the number of summands reduces from 65536 each iteration to 165 each iteration.

For individual $i$, let $A_{i1}$ be the number of "00" haplotypes, $A_{i2}$ be the number of "01" haplotypes, $A_{i3}$ be the number of "10" haplotypes, and $A_{i4}$ be the number of "11" haplotypes. Let $\boldsymbol{A}_i = (A_{i1}, A_{i2}, A_{i3}, A_{i4})$. Then $\boldsymbol{A} \sim \text{Multinom}(K, \boldsymbol{p})$, where $\boldsymbol{p} = (p_1, p_2, p_3, p_4)$ are the haplotype frequencies. Let $\boldsymbol{y}_i = (y_{i1}, y_{i2})$ be the data at loci 1 and 2 for individual $i$. We assume the user provides $p(y_{i1}|g_1)$ and $p(y_{i2}|g_2)$, the genotype likelihoods given genotypes $g_1$ and $g_2$ for individual $i$ at loci 1 and 2. Let $\boldsymbol{y} = (\boldsymbol{y}_1, \ldots, \boldsymbol{y}_n)$ and $\boldsymbol{A} = (\boldsymbol{A}_1, \ldots, \boldsymbol{A}_n)$. Then the complete log-likelihood is:

$$p(\boldsymbol{y}, \boldsymbol{A}|\boldsymbol{p}) = \sum_{i=1}^{n} \sum_{\substack{\boldsymbol{a} \text{ s.t.} \\ a_1+a_2+a_3+a_4=K}} I(\boldsymbol{A}_i = \boldsymbol{a}) \log\left[p(y_{i1}|a_3 + a_4)p(y_{i2}|a_2 + a_4)Pr(\boldsymbol{a}|\boldsymbol{p})\right]. \tag{S12}$$

The E-step involves calculating the following posterior probabilities:

$$w_{i\boldsymbol{a}} := Pr(\boldsymbol{a}|\boldsymbol{y}_i, \boldsymbol{p}^{(old)}) \tag{S13}$$

$$= \frac{p(y_{i1}|a_3 + a_4)p(y_{i2}|a_2 + a_4)Pr(\boldsymbol{a}|\boldsymbol{p}^{(old)})}{\sum_{\substack{\boldsymbol{a} \text{ s.t.} \\ a_1+a_2+a_3+a_4=K}} p(y_{i1}|a_3 + a_4)p(y_{i2}|a_2 + a_4)Pr(\boldsymbol{a}|\boldsymbol{p}^{(old)})} \tag{S14}$$

$$= \frac{p(y_{i1}|a_3 + a_4)p(y_{i2}|a_2 + a_4)\,\text{Multinom}(\boldsymbol{a}|K, \boldsymbol{p}^{(old)})}{\sum_{\substack{\boldsymbol{a} \text{ s.t.} \\ a_1+a_2+a_3+a_4=K}} p(y_{i1}|a_3 + a_4)p(y_{i2}|a_2 + a_4)\,\text{Multinom}(\boldsymbol{a}|K, \boldsymbol{p}^{(old)})} \tag{S15}$$

(S16)

The M-step thus involves maximizing the following objective function:

$$\sum_{i=1}^{n} \sum_{\substack{\boldsymbol{a} \text{ s.t.} \\ a_1+a_2+a_3+a_4=K}} w_{i\boldsymbol{a}} \log\left[p(y_{i1}|a_3 + a_4)p(y_{i2}|a_2 + a_4)Pr(\boldsymbol{a}|\boldsymbol{p})\right] \tag{S17}$$

$$= \sum_{i=1}^{n} \sum_{\substack{\boldsymbol{a} \text{ s.t.} \\ a_1+a_2+a_3+a_4=K}} w_{i\boldsymbol{a}} \log\left[Pr(\boldsymbol{a}|\boldsymbol{p})\right] + C, \tag{S18}$$

where $C$ is some constant with respect to $\boldsymbol{p}$. Using Lagrange multipliers, we find that the update is

$$\eta_\ell := \sum_{i=1}^{n} \sum_{\substack{\boldsymbol{a} \text{ s.t.} \\ a_\ell > 0 \text{ and} \\ a_1+a_2+a_3+a_4=K}} a_\ell w_{i\boldsymbol{a}} \tag{S19}$$

$$p_\ell^{(new)} = \frac{\eta_\ell}{\sum_\ell \eta_\ell} \tag{S20}$$

If one uses a Dirichlet($\boldsymbol{\alpha}$) prior on the haplotype proportions, then (S19) is modified to

$$\eta_\ell := \sum_{i=1}^{n} \sum_{\substack{\boldsymbol{a} \text{ s.t.} \\ a_\ell > 0 \text{ and} \\ a_1+a_2+a_3+a_4=K}} a_\ell w_{i\boldsymbol{a}} + (\alpha_\ell - 1). \tag{S21}$$

We also implemented a gradient ascent procedure to maximize (10). Optimization was performed over the unit 3-simplex using the unconstrained transformed parameter space used in Betancourt [2012] before back-transforming to the original parameter space. When LD is close to 1, this can cause MLEs on the boundary of the parameter space and, thus, nonsensical standard error estimates (described in Section S8). We thus take the approach of [Agresti and Coull, 1998] and add a small penalty on the haplotype frequencies. This penalty is equivalent to placing a Dirichlet(2,2,2,2) prior on the haplotype frequencies and corresponds to the "add two" rule. We call the resulting penalized MLEs $\hat{D}_{gl}$, $\hat{D}'_{gl}$, and $\hat{r}_{gl}$ for "genotype likelihoods".

### S3 Moments of genotypes

In this section, we derive the moments for the genotypes at two loci under the assumption of HWE. The calculations are simple, but demonstrate that composite measures of LD are equal to haplotypic measures of LD for populations in HWE. Let

$$(X_1, X_2, X_3, X_4) \sim \text{Multinom}(K, p_1, p_2, p_3, p_4), \tag{S22}$$

where $X_1$ are the counts of haplotype 00, $X_2$ are the counts of haplotype 10, $X_3$ are the counts of haplotype 01, and $X_4$ are the counts of haplotype 11. Then we have the following moments of multinomial counts:

$$E[X_i] = Kp_i \tag{S23}$$

$$\text{var}(X_i) = Kp_i(1 - p_i) \tag{S24}$$

$$\text{cov}(X_i, X_j) = -Kp_ip_j \text{ when } i \neq j. \tag{S25}$$

Let

$$G_1 = X_2 + X_4 \tag{S26}$$

$$G_2 = X_3 + X_4. \tag{S27}$$

Then

$$p_A = (p_2 + p_4) \tag{S28}$$
$$p_B = (p_3 + p_4) \tag{S29}$$
$$E[G_1] = K(p_2 + p_4) = Kp_A \tag{S30}$$
$$E[G_2] = K(p_3 + p_4) = Kp_B \tag{S31}$$
$$\begin{aligned}
\operatorname{var}(G_1) &= \operatorname{var}(X_2) + \operatorname{var}(X_4) + 2\operatorname{cov}(X_2, X_4) && \text{(S32)}\\
&= Kp_2(1-p_2) + Kp_4(1-p_4) - 2Kp_2p_4 && \text{(S33)}\\
&= K(p_2+p_4)(1-(p_2+p_4)) && \text{(S34)}\\
&= Kp_A(1-p_A) && \text{(S35)}\\
\operatorname{var}(G_2) &= K(p_3+p_4)(1-(p_3+p_4)) && \text{(S36)}\\
&= Kp_B(1-p_B) && \text{(S37)}
\end{aligned}$$

We will now derive the covariance between $G_1$ and $G_2$:

$$\begin{aligned}
K\Delta &= \operatorname{cov}(G_1, G_2) && \text{(S38)}\\
&= E[G_1G_2] - E[G_1]E[G_2] && \text{(S39)}\\
&= E[X_2X_3] + E[X_2X_4] + E[X_3X_4] + E[X_4^2] - E[G_1]E[G_2] && \text{(S40)}\\
&= K(K-1)(p_2p_3 + p_2p_4 + p_3p_4) + Kp_4(1-p_4) + K^2p_4^2 - K^2(p_2+p_4)(p_3+p_4) && \text{(S41)}\\
&= K[p_4 - (p_2+p_4)(p_3+p_4)] && \text{(S42)}\\
&= KD && \text{(S43)}
\end{aligned}$$

The correlation between $G_1$ and $G_2$ is

$$\begin{aligned}
\rho &= \operatorname{cor}(G_1, G_2) && \text{(S44)}\\
&= \frac{\operatorname{cov}(G_1, G_2)}{\sqrt{\operatorname{var}(G_1)\operatorname{var}(G_2)}} && \text{(S45)}\\
&= \frac{KD}{\sqrt{K^2p_A(1-p_A)p_B(1-p_B)}} && \text{(S46)}\\
&= \frac{D}{\sqrt{p_A(1-p_A)p_B(1-p_B)}} && \text{(S47)}\\
&= r. && \text{(S48)}
\end{aligned}$$

Under HWE, $\Delta_e$ defined in (20) is equal to

$$\begin{aligned}
\Delta_e &= \begin{cases} \min\{p_Ap_B, (1-p_A)(1-p_B)\} & \text{if } \Delta < 0,\\ \min\{p_A(1-p_B), (1-p_A)p_B\} & \text{if } \Delta > 0 \end{cases} && \text{(S49)}\\
&= D_{max}. && \text{(S50)}
\end{aligned}$$

Thus,

$$\Delta'_a := \frac{\frac{1}{K}\operatorname{cov}(G_1, G_2)}{\Delta_e} \tag{S51}$$

$$= D/D_{max} \tag{S52}$$

$$= D'. \tag{S53}$$

#### S4 Burrows' Δ and genotype covariance

Let $G_A$ and $G_B$ be the genotypes of a diploid at loci 1 and 2. Then the covariance between $G_A$ and $G_B$ is

$$\frac{1}{2}\operatorname{cov}(G_A, G_B) = \frac{1}{2}(E[G_A G_B] - E[G_A]E[G_B]) \tag{S54}$$

$$= \frac{1}{2}\sum_{g_A=0}^{2}\sum_{g_B=0}^{2} g_A g_B q_{g_A g_B} - 2p_A p_B \tag{S55}$$

$$= 2q_{22} + q_{21} + q_{12} + \frac{1}{2}q_{11} - 2p_A p_B \tag{S56}$$

$$= (12). \tag{S57}$$

#### S5 Closed-form bounds for Δ conditional on allele frequencies

**Theorem S1.** *Suppose $X$ and $Y$ are random variables such that $0 \leq X, Y \leq K$ almost surely. Furthermore, suppose $E[X] = Kp_A$ and $E[Y] = Kp_B$. Then*

$$E[XY] \leq K^2 \min(p_A, p_B). \tag{S58}$$

*Proof.* The following proof was provided by Dr. Y. Samuel Wang, University of Chicago, via a personal correspondence.

$$E[XY] = E[|XY|] \tag{S59}$$

$$\leq \min_{\substack{p,q\geq 1\\ 1/p+1/q=1}} E[|X|^p]^{1/p} E[|Y|^q]^{1/q} \tag{S60}$$

$$\leq \min_{\substack{p,q\geq 1\\ 1/p+1/q=1}} \max_{\substack{\tilde{X},\tilde{Y}\\ E[\tilde{X}]=Kp_A, E[\tilde{Y}]=Kp_B\\ 0\leq\tilde{X},\tilde{Y}\leq K}} E[|\tilde{X}|^p]^{1/p} E[|\tilde{Y}|^q]^{1/q} \tag{S61}$$

$$= \min_{\substack{p,q\geq 1\\ 1/p+1/q=1}} [K^p p_A]^{1/p} [K^q p_B]^{1/q} \tag{S62}$$

$$= \min_{\substack{p,q\geq 1\\ 1/p+1/q=1}} K^2 p_A^{1/p} p_B^{1/q} \tag{S63}$$

$$\leq K^2 \min(p_A, p_B). \tag{S64}$$

Equation (S60) follows by Hölder's inequality. Equation (S62) holds because for any $p \geq 1$, the maximum over $\tilde{X}$ is achieved when $Pr(\tilde{X} = K) = p_A$ and $Pr(\tilde{X} = 0) = 1 - p_A$. Similarly, the maximum over $\tilde{Y}$ is achieved when $Pr(\tilde{Y} = K) = p_B$ and $Pr(\tilde{Y} = 0) = 1 - p_B$. Equation (S64) results by letting $p = \infty$ and $q = 1$ when $p_B \leq p_A$, and letting $p = 1$ and $q = \infty$ when $p_B \geq p_A$. □

**Theorem S2.** *Let $X$ and $Y$ be two random variables, each with support on $\{0, 1, \ldots, K\}$. Furthermore, suppose that $E[X] = Kp_A$ and $E[Y] = Kp_B$. Then*

$$-K^2 \min\{p_A p_B, (1 - p_A)(1 - p_B)\} \leq \text{cov}(X, Y) \leq K^2 \min\{p_A(1 - p_B), (1 - p_A)p_B\}, \tag{S65}$$

*and these bounds are tight.*

*Proof.*

$$\begin{aligned}
\text{cov}(X, Y) &= E[XY] - E[X]E[Y] && \text{(S66)}\\
&\leq K^2 \min(p_A, p_B) - E[X]E[Y] \text{ (Theorem S1)} && \text{(S67)}\\
&= K^2 \min(p_A, p_B) - K^2 p_A p_B && \text{(S68)}\\
&= K^2 \min\{p_A(1 - p_B), (1 - p_A)p_B\}. && \text{(S69)}
\end{aligned}$$

Bound (S69) is tight since it is achieved when $Pr(X = K) = p_A$, $Pr(X = 0) = 1 - p_A$, $Pr(Y = K) = p_B$, and $Pr(Y = 0) = 1 - p_B$.

To prove the lower bound, first set $U := K - Y$. Then $E[U] = K(1 - p_B)$ and we have

$$\text{cov}(X, U) \leq K^2 \min\{p_A p_B, (1 - p_A)(1 - p_B)\}. \tag{S70}$$

But $\text{cov}(X, U) = -\text{cov}(X, Y)$, and thus

$$\text{cov}(X, Y) \geq -K^2 \min\{p_A p_B, (1 - p_A)(1 - p_B)\}. \tag{S71}$$

□

#### S6 Alternative algorithm for obtaining bounds on $\Delta$ conditional on marginal genotype distributions

In Section 2.3, though we were able to find closed-form bounds on $\Delta$ when conditioning on the genotype expectations, we resorted to the methods of linear programming to numerically find the bounds on $\Delta$ when conditioning on the marginal distributions. Under more general conditions, Whitt [1976] characterized the maximum and minimum correlation between two random variables given fixed marginals, corresponding to scenario (15)–(16). Specifically, given random variables $X$ and $Y$ with inverse cumulative distribution functions $F^{-1}(\cdot)$ and $G^{-1}(\cdot)$, Whitt [1976] found that

$$\text{cor}(F^{-1}(U), G^{-1}(1 - U)) \leq \text{cor}(X, Y) \leq \text{cor}(F^{-1}(U), G^{-1}(U)), \tag{S72}$$

where $U$ is a Uniform$(0, 1)$ random variable. Using (S72), Leonov and Qaqish [2020] derived a purpose-built algorithm to find the bounds on the correlation given two random variables that follow general categorical distributions. This algorithm could be used to solve (15)–(16) and might be computationally faster. However, solving the linear program (15)–(16) is not the computational bottleneck we face when estimating LD. The time to solve (15)–(16) for an octoploid species is on the order of a millisecond, whereas the optimization procedures discussed in Section 2.4 take about half a second (Figures S11 and S15). We thus leave improved optimization of (15)–(16) as future work.

#### S7 A decomposition of $\Delta$

Cockerham and Weir [1977] derived their composite LD measure by summing haplotypic and non-haplotypic LD. From a different point of view, this can be seen as a decomposition of $\Delta$. We will now derive a generalized decomposition of (13) for polyploids. Let $z_{kA}$ be the indicator variable that equals 1 if an individual has the A allele on locus 1 of chromosome $k$, and 0 otherwise. Similarly, let $z_{kB}$ be the indicator variable that equals 1 if an individual has the B allele on locus 2 of chromosome $k$, and 0 otherwise. Then the genotype for an individual equals

$$G_A = \sum_{k=1}^{K} z_{kA} \text{ and } G_B = \sum_{k=1}^{K} z_{kB}. \tag{S73}$$

Thus,

$$\begin{aligned}
\Delta &= \frac{1}{K}\operatorname{cov}(G_A, G_B) && \text{(S74)}\\
&= \frac{1}{K}\operatorname{cov}\Big(\sum_{k_A=1}^{K} z_{k_A A}, \sum_{k_B=1}^{K} z_{k_B B}\Big) && \text{(S75)}\\
&= \frac{1}{K}\sum_{k_A=1}^{K}\sum_{k_B=1}^{K}\operatorname{cov}(z_{k_A A}, z_{k_B B}). && \text{(S76)}
\end{aligned}$$

Equation (S76) is as simple a decomposition as can be attained for $\Delta$ without further assumptions. However, in the special case of auto- and allopolyploidy, we can further reduce (S76). In the case of autopolyploidy, we have the following two identities:

$$\begin{aligned}
D_{AB} &:= \operatorname{cov}(z_{iA}, z_{iB}) = \operatorname{cov}(z_{i'A}, z_{i'B}) \text{ for all } i \text{ and } i', \text{ and} && \text{(S77)}\\
D_{A/B} &:= \operatorname{cov}(z_{iA}, z_{jB}) = \operatorname{cov}(z_{i'A}, z_{j'B}) \text{ for all } i \neq j \text{ and } i' \neq j', && \text{(S78)}
\end{aligned}$$

where $D_{AB}$ is haplotypic LD and $D_{A/B}$ is non-haplotypic LD. Thus, for autopolyploids, we obtain

$$\Delta = D_{AB} + (K-1)D_{A/B}, \tag{S79}$$

which more clearly demonstrates the generalization of the decomposition in Cockerham and Weir [1977] from diploids to autopolyploids. In the case of HWE, $D_{A/B} = 0$ and we obtain $\Delta = D_{AB}$.

In the case of allopolyploidy with an even ploidy level, there are $K/2$ homologous pairs. There are three types of LD that may occur. First, there is haplotypic LD in homologous pair $i$, denoted $D_{ABi}$. Second, there is non-haplotypic LD in homologous pair $i$, denoted $D_{A/Bi}$. Third, there is non-haplotypic LD between a chromosome in homologous pair $i$ and a chromosome in homologous pair $j$, denoted $D_{A/Bij}$. Thus, in the case of allopolyploids, we obtain

$$\Delta = \frac{2}{K}\left[\sum_{i=1}^{K/2} D_{ABi} + \sum_{i=1}^{K/2} D_{A/Bi} + 4\sum_{i=1}^{K/2-1}\sum_{j=i+1}^{K/2} D_{A/Bij}\right]. \tag{S80}$$

In the case of HWE, we have $D_{A/Bi} = 0$ for all $i$ and $D_{A/Bij} = 0$ for all $(i, j)$. This indicates that for allopolyploids, in HWE, the composite measure of LD (S76) is the average of haplotypic LDs for all homologous pairs:

$$\Delta = \frac{1}{K/2}\sum_{i=1}^{K/2} D_{ABi}. \tag{S81}$$

### S8 Maximum likelihood standard error calculations

**Haplotypic LD standard errors** We now provide a description for how to obtain standard errors for $\hat{D}_{gl}$, $\hat{D}'_{gl}$, and $\hat{r}_{gl}$ (Section 2.2) using standard maximum likelihood theory. First, we numerically approximated the Hessian matrix of $\mathcal{L}(p_{ab}, p_{Ab}, p_{aB})$ evaluated at the maximum likelihood estimators $(\hat{p}_{ab}, \hat{p}_{Ab}, \hat{p}_{aB})$, $\boldsymbol{H} := (\frac{\partial^2\mathcal{L}}{\partial p_i \partial p_j})|_{p_{ab}=\hat{p}_{ab}, p_{Ab}=\hat{p}_{Ab}, p_{aB}=\hat{p}_{aB}}$. This likelihood can be either that using genotypes (9) or that using genotype likelihoods (10), both of which are functions of $(p_{ab}, p_{Ab}, p_{aB})$ since the haplotype frequencies must sum to one. Standard maximum likelihood theory guarantees that for large $n$, the limiting covariance matrix of $(\hat{p}_{ab}, \hat{p}_{Ab}, \hat{p}_{aB})$ is well approximated by $-\boldsymbol{H}^{-1}$ [Lehmann and Casella, 1998]. The MLEs of the various LD measures ($\hat{D}$, $\hat{D}'$, and $\hat{r}$) are all functions of $(\hat{p}_{ab}, \hat{p}_{Ab}, \hat{p}_{aB})$, and so we can use the $\delta$-method to obtain the limiting variance of these LD estimators. For example, let $\boldsymbol{g} = (\frac{\partial r}{\partial p_{ab}}, \frac{\partial r}{\partial p_{Ab}}, \frac{\partial r}{\partial p_{aB}})^\intercal$ be the gradient of $r$ with respect to $(p_{ab}, p_{Ab}, p_{aB})$ evaluated at $(\hat{p}_{ab}, \hat{p}_{Ab}, \hat{p}_{aB})$. Then the limiting variance of $\hat{r}$ is $-\boldsymbol{g}^\intercal\boldsymbol{H}^{-1}\boldsymbol{g}$. The gradient calculations are all standard and so have been omitted. We have implemented this procedure for standard error calculation in our software.

**Composite LD standard errors using the general categorical genotype distribution** To calculate standard errors of $\hat{\rho}_{gc}$, $\hat{\Delta}_{gc}$ and $\hat{\Delta}_{gc}$ (Section 2.4), it would be possible to take the same approach as in Section 2.2 and derive asymptotic variances using the Fisher information and appealing to the $\delta$-method (Section S9). However, issues arise with ploidies greater than two. As there are $(K+1)^2$ possible genotype conditions ($(K+1)^2 - 1$ free parameters), except for very large $n$ it will not be uncommon for the MLEs to occur on the boundary of the parameter space, thereby violating the regulatory conditions for the asymptotic standard errors of the MLE [Lehmann and Casella, 1998]. In such cases, we appeal to bootstrap standard errors [Efron, 1979] for the composite measures of LD.

**Composite LD standard errors using the proportional normal genotype distribution** Let $\boldsymbol{\Sigma} = \boldsymbol{L}\boldsymbol{L}^\intercal$ be the Cholesky decomposition of $\boldsymbol{\Sigma}$, where $\boldsymbol{L}$ is a lower-triangular matrix. To obtain the standard errors of $\hat{\rho}_{pn}$, $\hat{\Delta}_{pn}$ and $\hat{\Delta}'_{pn}$ (Section 2.4) we first obtain the Hessian of the log-likelihood (25) evaluated at the maximum likelihood estimators $(\hat{\boldsymbol{\mu}}, \text{vec}(\hat{\boldsymbol{L}}))$, where $\text{vec}(\boldsymbol{L})$ is the vectorization of the lower-triangle of $\boldsymbol{L}$. Call this Hessian $\boldsymbol{H}$. The MLEs of $\rho$, $\Delta$, and $\Delta'_a$ are all functions of the MLEs $(\hat{\boldsymbol{\mu}}, \text{vec}(\hat{\boldsymbol{L}}))$. To see this, set $q_{ij} = Pr(i, j|\boldsymbol{\mu}, \boldsymbol{\Sigma})$ and substitute the

$q_{ij}$'s in (S84), (S86), and (S95). Thus, they each admit a gradient for the functions mapping from $(\hat{\boldsymbol{\mu}}, \text{vec}(\hat{\boldsymbol{L}}))$ to $\hat{\rho}$, $\hat{\Delta}$, and $\hat{\Delta}'_a$. Call each gradient $\boldsymbol{g}$. Then the asymptotic variance of an estimator of composite LD is $-\boldsymbol{g}^{\intercal}\boldsymbol{H}\boldsymbol{g}^{\intercal}$. We have implemented all gradient calculations numerically. We additionally placed weakly informative priors over $\boldsymbol{\mu}$ and $\boldsymbol{\Sigma}$ as we noticed some scenarios resulted in divergent optimization behavior:

$$\boldsymbol{\mu} \sim N(\boldsymbol{0}, (K/2, K/2)^{\intercal}, \text{diag}((2K)^2, (2K)^2)),\ \boldsymbol{\Sigma} \sim Wishart_2(\boldsymbol{I}_2, 2). \tag{S82}$$

The induced distribution over $\boldsymbol{L}$ can be found by Bartlett's decomposition [Bartlett, 1934].

#### S9 MLE standard errors calculations when using the general categorical genotype distribution

The following are the derivatives relating to the log of (22) necessary to derive asymptotic standard errors of the MLEs when using the general categorical genotype distribution to estimate composite measures of LD. The Hessian of the log of (22) can be calculated in closed form:

$$\frac{dL}{dq_{ij}dq_{km}} = -\sum_{\ell=0}^{n} \frac{Pr(D_{\ell 1}|i)Pr(D_{\ell 2}|j)Pr(D_{\ell 1}|k)Pr(D_{\ell 2}|m)}{\left(\sum_{i=0}^{K}\sum_{j=0}^{K} Pr(D_{\ell 1}|i)Pr(D_{\ell 2}|j)q_{ij}\right)^2} \tag{S83}$$

For $\Delta$, we have

$$\Delta = \frac{1}{K}\sum_{i=0}^{K}\sum_{j=0}^{K} ijq_{ij} - \frac{1}{K}\left(\sum_{i=0}^{K} i \sum_{j=0}^{K} q_{ij}\right)\left(\sum_{j=0}^{K} j \sum_{i=0}^{K} q_{ij}\right) \tag{S84}$$

$$\frac{d\Delta}{dq_{\ell m}} = \frac{\ell m}{K} - \frac{\ell}{K}\left(\sum_{j=0}^{K} j \sum_{i=0}^{K} q_{ij}\right) - \frac{m}{K}\left(\sum_{i=0}^{K} i \sum_{j=0}^{K} q_{ij}\right), \tag{S85}$$

For $\rho^2$, we have

$$\rho^2 = \frac{K^2\Delta^2}{\operatorname{var}(G_A)\operatorname{var}(G_B)} \tag{S86}$$

$$\operatorname{var}(G_A) = \sum_{i=0}^{K} i^2 \sum_{j=0}^{K} q_{ij} - \left(\sum_{i=0}^{K} i \sum_{j=0}^{K} q_{ij}\right)^2 \tag{S87}$$

$$\operatorname{var}(G_B) = \sum_{j=0}^{K} j^2 \sum_{i=0}^{K} q_{ij} - \left(\sum_{j=0}^{K} j \sum_{i=0}^{K} q_{ij}\right)^2 \tag{S88}$$

$$\frac{d\operatorname{var}(G_A)}{dq_{\ell m}} = \ell^2 - 2\ell\left(\sum_{i=0}^{K} i \sum_{j=0}^{K} q_{ij}\right) \tag{S89}$$

$$\frac{d\operatorname{var}(G_B)}{dq_{\ell m}} = m^2 - 2m\left(\sum_{j=0}^{K} j \sum_{i=0}^{K} q_{ij}\right) \tag{S90}$$

$$\frac{d\rho^2}{dq_{\ell m}} = \frac{2K^2\Delta\frac{d\Delta}{dq_{\ell m}}}{\operatorname{var}(G_A)\operatorname{var}(G_B)} - \frac{K^2\Delta^2\frac{d\operatorname{var}(G_A)}{dq_{\ell m}}}{\operatorname{var}(G_A)^2\operatorname{var}(G_B)} - \frac{K^2\Delta^2\frac{d\operatorname{var}(G_B)}{dq_{\ell m}}}{\operatorname{var}(G_A)\operatorname{var}(G_B)^1} \tag{S91}$$

For $\Delta'$, we have

$$E[G_A] = \sum_{i=0}^{K} i \sum_{j=0}^{K} q_{ij} \tag{S92}$$

$$E[G_B] = \sum_{j=0}^{K} j \sum_{i=0}^{K} q_{ij} \tag{S93}$$

$$\Delta_e := \begin{cases} \min\{E[G_A]E[G_B], (K - E[G_A])(K - E[G_B])\}/K^2 & \text{if } \Delta < 0, \\ \min\{E[G_A](K - E[G_B]), (K - E[G_A])E[G_B]\}/K^2 & \text{if } \Delta > 0. \end{cases} \tag{S94}$$

$$\Delta' = \Delta/\Delta_e \tag{S95}$$

$$\frac{d\Delta'}{dq_{\ell m}} = \frac{\frac{d\Delta}{dq_{\ell m}}}{\Delta_e} - \frac{\Delta\frac{d\Delta_e}{dq_{\ell m}}}{\Delta_e^2} \tag{S96}$$

$$\frac{d\Delta_e}{dq_{\ell m}} = \begin{cases} \frac{\ell E[G_B] + mE[G_A]}{K^2} & \text{if } \Delta < 0 \\ & \text{and } E[G_A]E[G_B] < (K - E[G_A])(K - E[G_B]) \\ \frac{-\ell(K - E[G_B]) - m(K - E[G_A])}{K^2} & \text{if } \Delta < 0 \\ & \text{and } E[G_A]E[G_B] > (K - E[G_A])(K - E[G_B]) \\ \frac{\ell(K - E[G_B]) - mE[G_A]}{K^2} & \text{if } \Delta > 0 \\ & \text{and } E[G_A](K - E[G_B]) < (K - E[G_A])E[G_B] \\ \frac{-\ell E[G_B] + m(K - E[G_A])}{K^2} & \text{if } \Delta > 0 \\ & \text{and } E[G_A](K - E[G_B]) > (K - E[G_A])E[G_B] \end{cases} \tag{S97}$$

We will now provide an example of how to obtain standard errors for the MLEs. Let $\boldsymbol{q}$ =

$(q_{00}, q_{01}, \ldots, q_{ij}, \ldots, q_{KK})$, let $\hat{\boldsymbol{q}}$ be the MLEs of $\boldsymbol{q}$, and let $\boldsymbol{H}$ be the Hessian of the log-likelihood with elements (S83). Then standard maximum likelihood theory states that $\boldsymbol{H}^{-1/2}(\hat{\boldsymbol{q}} - \boldsymbol{q}) \rightarrow N(0, \boldsymbol{I})$. Since the covariance and correlation of genotypes are functions of the $q_{ij}$'s, we can use the $\delta$-method to obtain the limiting variances of the covariance and correlation of the genotypes. For example, if we set $\boldsymbol{g}$ to contain the elements of (S85), then the asymptotic variance we use for $\hat{\Delta}_{gc}$ is $-\boldsymbol{g}^{\intercal}\boldsymbol{H}^{-1}\boldsymbol{g}$.

The asymptotic standard errors are not valid at points for which the gradient does not exist, which for $\Delta'$ occur when $\Delta > 0$ and $E[G_A] = E[G_B]$, when $\Delta < 0$ and $E[G_A] + E[G_B] = K$, or when $\Delta = 0$. These situations occur with Lebesgue measure 0, and so should not invalidate the standard errors.

#### S10 Standard errors of moment-based estimators

The results in this section can be derived directly from well-known results in the literature [Example 6.6.4 Lehmann and Casella, 1998, e.g.]. These results hold only for multivariate normal random variables, which is not applicable when estimating LD. However, we found that for most estimators the approximations are decent (Figure S9). Improved asymptotic standard errors could be implemented by using the techniques described in Chapter 8 of Ferguson [2002].

Let $\hat{\rho}$ be the sample correlation between genotypes, let $\hat{z} = \operatorname{atanh}(\hat{\rho})$, let $\hat{\Delta}$ be the sample covariance between genotypes divided by $K$, let $\hat{\Delta}'$ be the sample estimator of $\Delta'$, let $\sigma_1^2$ be the variance of genotypes at locus 1, let $\sigma_2^2$ be the variance of genotypes at locus 2, let $\mu_1$ be the mean genotype at locus 1, and let $\mu_2$ be the mean genotype at locus 2. Then

$$\sqrt{n}(\hat{\rho}_{mom} - \rho) \rightarrow N\left(0, (1-\rho^2)^2\right) \tag{S98}$$

$$\sqrt{n}(\hat{\rho}^2_{mom} - \rho^2) \rightarrow N\left(0, 4\rho^2(1-\rho^2)^2\right) \tag{S99}$$

$$\sqrt{n}(\hat{z}_{mom} - z) \rightarrow N(0, 1) \tag{S100}$$

$$\sqrt{n}(\hat{\Delta}_{mom} - \Delta) \rightarrow N\left(0, \sigma_1^2\sigma_2^2/K^2 + \Delta^2\right) \tag{S101}$$

Equation (S99) follows from the $\delta$-method using (S98). Equation (S101) follows because $K\hat{\Delta}_{mom}$ is the sample covariance of dosages. Equation (S100) is well known [Fisher, 1921, Hotelling, 1953].

For the composite measure $\Delta'$ (19), since

$$\hat{\Delta}_e \rightarrow \Delta_e := \begin{cases} \min(\mu_1\mu_2, (K-\mu_1)(K-\mu_2))/K^2 & \text{if } \Delta < 0, \\ \min((K-\mu_1)\mu_2, \mu_1(K-\mu_2))/K^2 & \text{if } \Delta > 0, \end{cases} \tag{S102}$$

we have by Slutsky's theorem that

$$\sqrt{n}(\hat{\Delta}'_{mom} - \Delta') \rightarrow N\left(0, \frac{\sigma_1^2\sigma_2^2/K^2 + \Delta^2}{\Delta_e^2}\right). \tag{S103}$$

#### S11 Pairwise LD simulations when HWE is violated

In this section, we evaluate the performance of the various LD estimators when HWE is violated. We do this by simulating genotypes directly from the proportional bivariate normal distribution. In this case, since HWE is violated, the composite LD estimators (Section 2.4) are still appropriate measures of association, but the estimators of haplotypic LD (Section 2.2) are estimators under a misspecified model.

Each replication, given a ploidy $K$, we generated genotypes for 100 individuals from a proportional bivariate normal distribution with mean $\boldsymbol{\mu} \in \mathbb{R}^2$ and covariance matrix $\boldsymbol{\Sigma} = (\sigma_{ij}) \in \mathbb{R}^{2\times 2}$. Given these genotypes, we simulated read-counts using `updog`'s `rflexdog()` function at a specified read-depth, a 0.01 sequencing error rate, no allele bias, and an overdispersion value of 0.01. `Updog` was then used to generate genotype likelihoods, posterior mode genotypes, and posterior mean genotypes. These outputs were fed into `ldsep` to provide the estimators listed in Table 1. The parameters that varied within the simulation were: the ploidy $K \in \{2, 4, 6, 8\}$, the mean parameter $\boldsymbol{\mu} = (p_1K, p_2K)$ where $(p_1, p_2) \in \{(0.5, 0.5), (0.5, 0.75), (0.9, 0.9)\}$, the scale parameter $\sigma_{11} = \sigma_{22} \in \{K^2/4, K^2\}$, the association parameter $\sigma_{12} \in \{0, 0.5\sqrt{\sigma_{11}\sigma_{22}}, 0.9\sqrt{\sigma_{11}\sigma_{22}}\}$, and the read-depth $\in \{1, 5, 10, 50, 100\}$. Each unique combination of parameter values was replicated 200 times.

The conclusions for estimating $\rho^2$ when $\mu_1 = \mu_2 = K/2$ and $\sigma_{11} = \sigma_{22} = K^2/4$ are presented in Figures S12, S13, and S14. The results for other scenarios are similar and are available on Figshare (`https://doi.org/10.6084/m9.figshare.12765803.v1`). The general conclusions are:

- Composite measures perform the best under high levels of LD. Most methods are unbiased under low levels of LD, but only composite measures are unbiased under high levels of LD.
- The composite measure using genotype likelihoods and the proportional normal genotype distribution class ($\hat{\Delta}_{pn}, \hat{\Delta}'_{pn}, \hat{\rho}_{pn}$) generally had the best performance overall, while using the general categorical class of genotype distributions ($\hat{\Delta}_{gc}, \hat{\Delta}'_{gc}, \hat{\rho}_{gc}$) resulted in higher standard errors.

Computation times for each method are presented in Figure S15, the results being similar to those described in Section 3.1.

### S12 Pairwise LD simulations under preferential pairing and double reduction

In Section S11, we ran simulations under violations of HWE. However, the resulting genotype distributions were not interpretable to any specific biological process. So in this section, we evaluate our LD estimators using more interpretable violations from HWE, namely, preferential pairing and double reduction.

We used PedigreeSim [Voorrips and Maliepaard, 2012] to generate a population of individuals. We created a founder population of one thousand tetraploid individuals. Each individual contained two chromosomes of entirely reference (A) and two chromosomes of entirely alternative (a) alleles at each SNP. This results in an overall allele frequency at each SNP in the founder population of 0.5, which means that both $D'$ and $r^2$ in this founder population both start at 1 between all pairs of SNPs. Preferential pairing/allopolyploidy will result in underdispersed genotype distributions, but only in cases where the subgenomes have different allele frequencies (Figure S16). Therefore, to induce allele frequencies of 0.9 and 0.1 on each subgenome, respectively, we set 80% of individuals to have one subgenome that contains both all-A chromosomes and one subgenome that contains both all-a chromosomes. The remaining 20% of individuals have both one all-A and one all-a chromosome in each subgenome. Letting $n$ be the number of individuals in the dataset, this results in the desired allele frequencies of each subgenome because

$$\frac{2 \times 0.8n + 0.2n}{2n} = 0.9, \tag{S104}$$

where $2 \times 0.8n$ is the number of A's contributed by individuals with all-A subgenomes, and $0.2n$ is the number of A's contributed by the other individuals. Pedigrees were generated by random mating, keeping the population size constant, for a total of ten generations. While in diploids, HWE is attained after one generation, in autopolyploids, HWE is only approached asymptotically [Haldane, 1930]. However, after ten generations of random mating, an autopolyploid species is already almost at HWE (Table S1). This allows us to study the effects of double reduction and preferential pairing, as any further deviations from HWE should only be a result of these forces. After ten generations of random mating, it appears that the effects of double reduction are rather modest, and the effects of preferential pairing are only practically significant in the absence of quadrivalent formation (Table S2). Thus, for these simulations, we expect major deviations from autopolyploid HWE only when the individuals are pure (or nearly pure) allopolyploids.

To study the effects of preferential pairing, we varied the probability of homologous pairing to be 1/3 (autopolyploids), 2/3 (intermediate levels of preferential pairing), and 1 (allopolyploids). To study the effects of double reduction, we varied the proportion of quadrivalents to be 0, 1/3, and 2/3, where higher levels of quadrivalent pairing should result in increased double reduction. The upper bound of 2/3 was chosen because this is the maximum frequency of quadrivalents under the "natural pairing" model where the ends of chromosomes pair randomly [Morrison and Rajhathy, 1960]. The centromere was placed at the midpoint of a 100 cM chromosome. We calculated pairwise LD between SNPs located at 50 cM, 51 cM, 60 cM, 70 cM, 80 cM, 90 cM, 99 cM, and 100 cM, where SNPs closer together should result in larger levels of LD. SNPs closer to the centromere (50 cM) should experience less double reduction.

We took the genotypes of the one thousand individuals in the tenth generation and calculated the $\rho^2$ between each pair of SNPs, considering this the "true" $\rho^2$ (Figure S19). We then randomly sampled 100 individuals, took their genotypes, and generated their read-counts using the `rflexdog()` function from the `updog` R package, using a sequencing error rate of 0.01, no allele bias, and an overdispersion value of 0.01. We then used `updog` to generate genotype likelihoods, posterior mean genotypes, and posterior mode genotypes. These were fed into `ldsep` to obtain $\hat{r}_g^2$, $\hat{r}_{gl}^2$, $\hat{\rho}_{mom}^2$, $\hat{\rho}_{gc}^2$, and $\hat{\rho}_{pn}^2$.

To visualize what LD looks like after 10 generations of random mating at the chosen SNPs, we generated heatmaps of $\rho^2$ between SNPs placed at 50 cM, 51 cM, 60 cM, 70 cM, 80 cM, 90 cM, 99 cM, and 100 cM, under different levels of quadrivalent formation and preferential pairing (Figure S17). These were calculated from 10000 individuals. A pairs plot comparing these LD values is presented in Figure S18. In general, preferential pairing and double reduction have very little effect on the final "true" values of LD.

For each unique combination of the probability of homologous pairing (1/3, 2/3, and 1) and the proportion of quadrivalents (0, 1/3, and 2/3), we ran 100 replications, each time obtaining a "true" $\rho^2$ and the corresponding estimates. Boxplots for the difference between the true $\rho^2$ and the estimated $\rho^2$ for each method are presented in Figures S20–S25. We see generally that the effects of double reduction and moderate levels of preferential pairing are rather modest. This is to be expected as these conditions only result in minor deviations from HWE (Table S2). The biggest effects occur under under complete preferential pairing (allopolyploids) with no quadrivalent formation. Under this scenario, all methods have large levels of negative bias (indicating attenuation toward 0) when the LD is large (Figures S20 and S23). This bias is reduced in the methods that use the genotype likelihoods and not genotype point estimates. The effects of assuming autopolyploid HWE when we have allopolyploids is very severe when the LD is small (Figures S21, S22, S24, and S25). For allopolyploids, assuming HWE results in extreme overestimation of LD, indicating SNPs are much more associated than in reality. In these cases, the composite estimators outperform the haplotypic estimators.

#### S13 LD estimates using data from *Andropogon gerardii*

In this section, we evaluate our LD estimators using the genotyping-by-sequencing data from McAllister and Miller [2016], downloaded from Dryad as a variant call format file [McAllister and Miller, 2017]. These data come from natural populations of *Andropogon gerardii*, where reads were mapped onto the *Sorghum bicolor* genome. *A. gerardii* contains two common cytotypes: hexaploid ($2n = 6x = 60$) and enneaploid ($2n = 9x = 90$). All results in this section use only hexaploid individuals. Unlike the data from Uitdewilligen et al. [2013], the individuals from McAllister and Miller [2016] were sequenced at relatively low depth (on the order of $10\times$ versus $60\times$).

As in Section 3.3, we selected two arbitrary regions of the *Sorghum bicolor* genome, located on two different chromosomes, and extracted all biallelic SNPs from these two regions. SNPs were discarded if they contained an alternative allele frequency less than 0.1 or greater than 0.9. SNPs were also discarded if their average read-depth was less than 3. We then estimated genotypes using `updog` [Gerard et al., 2018] using a proportional normal prior class [Gerard and Ferrão, 2019]. The resulting genotype likelihoods were used to estimate pairwise LD between all SNPs.

Heat-maps of all pairwise LD estimates are provided in Figures S33–S37. It at first appears that there is a relatively rapid decay of pairwise LD. However, this is likely because the genomic regions span 100 kb, which is much larger than the regions explored in Section 3.3. The second dominant result is that estimating LD by maximum likelihood using posterior mode genotypes performed very poorly. The other methods performed comparably. Though, there was greater noise in the methods that estimate composite LD using genotype likelihoods. When the LD estimates are shrunk using the hierarchical shrinkage procedure of Stephens [2016] and Dey and Stephens [2018], the results appear to be very close (Figures S38–S41), indicating the signal-to-noise ratio was very similar for the different estimators. This shrinkage was performed on the Fisher-$z$ transformation [Fisher, 1921] of the estimated Pearson correlation, whose distribution in simulations appears to be very well approximated by the normal distribution (Figure S10). The major difference between the shrunken LD heatmaps is that genotype likelihood methods result in a fewer number of non-zero LD estimates, but each are of higher magnitude.

#### S14 Genotype likelihoods up to a proportional constant

Some genotyping programs do not explicitly return genotype likelihoods, but rather quantities that are proportional to the genotype likelihoods, where this constant of proportionality does not depend on the genotypes or the parameters being estimated. For example, McKenna et al. [2010] returns posterior probabilities for each genotype, but since they use a uniform prior over the genotypes, these posteriors are proportional to the genotype likelihoods. We will now briefly explain why it is enough to obtain the genotype likelihoods up to a proportional constant and use these in

(10), (22), and (25) to obtain maximum likelihood estimates. Let $p(y_{iA}|g_A)$ and $p(y_{iB}|g_B)$ be the genotype likelihoods for individual $i$ at loci 1 and 2, respectively, for genotypes $g_A \in \{0, 1, \ldots, K\}$ and $g_B \in \{0, 1, \ldots, K\}$. Suppose the user provides not the genotype likelihoods, but $f(y_{iA}|g_A) := p(y_{iA}|g_A)/c_{iA}$ and $f(y_{iB}|g_B) := p(y_{iB}|g_B)/c_{iB}$ for unknown constants $c_{iA}$ and $c_{iB}$ which do not depend on $g_A$ and $g_B$. If we maximize the following over $p_A$, $p_B$, and $D$,

$$\sum_{i=1}^{n} \log\left(\sum_{g_B=0}^{K}\sum_{g_A=0}^{K} f(y_{iA}|g_A)f(y_{iB}|g_B)Pr(g_A, g_B|D, p_A, p_B)\right) \tag{S105}$$

$$= \sum_{i=1}^{n} \log\left(\sum_{g_B=0}^{K}\sum_{g_A=0}^{K} (p(y_{iA}|g_A)/c_{iA})(p(y_{iB}|g_B)/c_{iB})Pr(g_A, g_B|D, p_A, p_B)\right) \tag{S106}$$

$$= \sum_{i=1}^{n} \log\left(\frac{1}{c_{iA}c_{iB}}\sum_{g_B=0}^{K}\sum_{g_A=0}^{K} p(y_{iA}|g_A)p(y_{iB}|g_B)Pr(g_A, g_B|D, p_A, p_B)\right) \tag{S107}$$

$$= \sum_{i=1}^{n} \log\left(\sum_{g_B=0}^{K}\sum_{g_A=0}^{K} p(y_{iA}|g_A)p(y_{iB}|g_B)Pr(g_A, g_B|D, p_A, p_B)\right) - \sum_{i=1}^{n}\log(c_{iA}c_{iB}), \tag{S108}$$

then we are equivalently maximizing equation (10), since $\sum_{i=1}^{n}\log(c_{iA}c_{iB})$ does not depend on $p_A$, $p_B$, and $D$. Identical arguments hold for (22) and (25). In the case of McKenna et al. [2010], we have

$$c_{iA} = \sum_{g_A=0}^{K} p(y_{iA}|g_A), \text{ and} \tag{S109}$$

$$c_{iB} = \sum_{g_B=0}^{K} p(y_{iB}|g_B). \tag{S110}$$

which does not depend on the genotypes ($g_A$ and $g_B$) because we are summing over them.

#### S15 Supplementary figures and tables

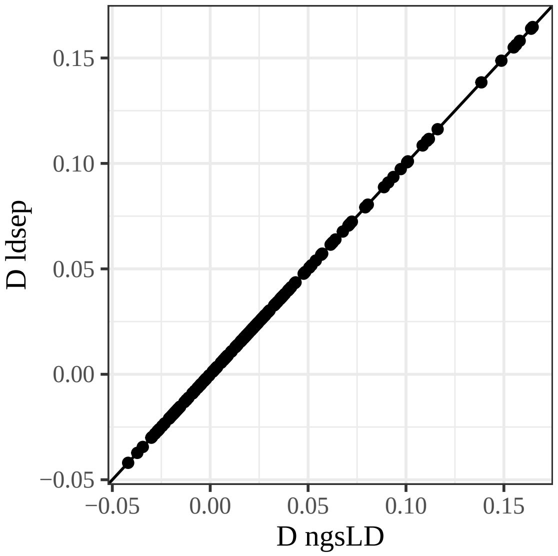


Figure S1: Maximum likelihood estimates of $D$ between 20 simulated loci of 100 diploid individuals in HWE as calculated by `ngsLD` ($x$-axis) and `ldsep` ($y$-axis). The line is the $y = x$ line. The estimates are identical.

| Dosage | 0 | 1 | 2 | 3 | 4 |
|---:|---:|---:|---:|---:|---:|
| Generation 1 | 0 | 0 | 1 | 0 | 0 |
| Generation 2 | .0278 | .2222 | .5000 | .2222 | .0278 |
| Generation 3 | .0494 | .2469 | .4074 | .2469 | .0494 |
| Generation 4 | .0580 | .2497 | .3848 | .2497 | .0580 |
| Generation 5 | .0610 | .2500 | .3781 | .2500 | .0610 |
| Generation 6 | .0620 | .2500 | .3760 | .2500 | .0620 |
| Generation 7 | .0623 | .2500 | .3753 | .2500 | .0623 |
| Generation 8 | .0624 | .2500 | .3751 | .2500 | .0624 |
| Generation 9 | .0625 | .2500 | .3750 | .2500 | .0625 |
| Generation 10 | .0625 | .2500 | .3750 | .2500 | .0625 |
| HWE | .0625 | .2500 | .3750 | .2500 | .0625 |

Table S1: Genotype frequencies for ten generations of random mating of an autopolyploid species, starting at a founder population where all individuals have a dosage of 2. The bottom row is the genotype distribution under Hardy Weinberg equilibrium. After 10 generations, the genotype distribution of the population is close to that of a population in HWE, where each genotype frequency is equal to that under HWE up to four decimal places.

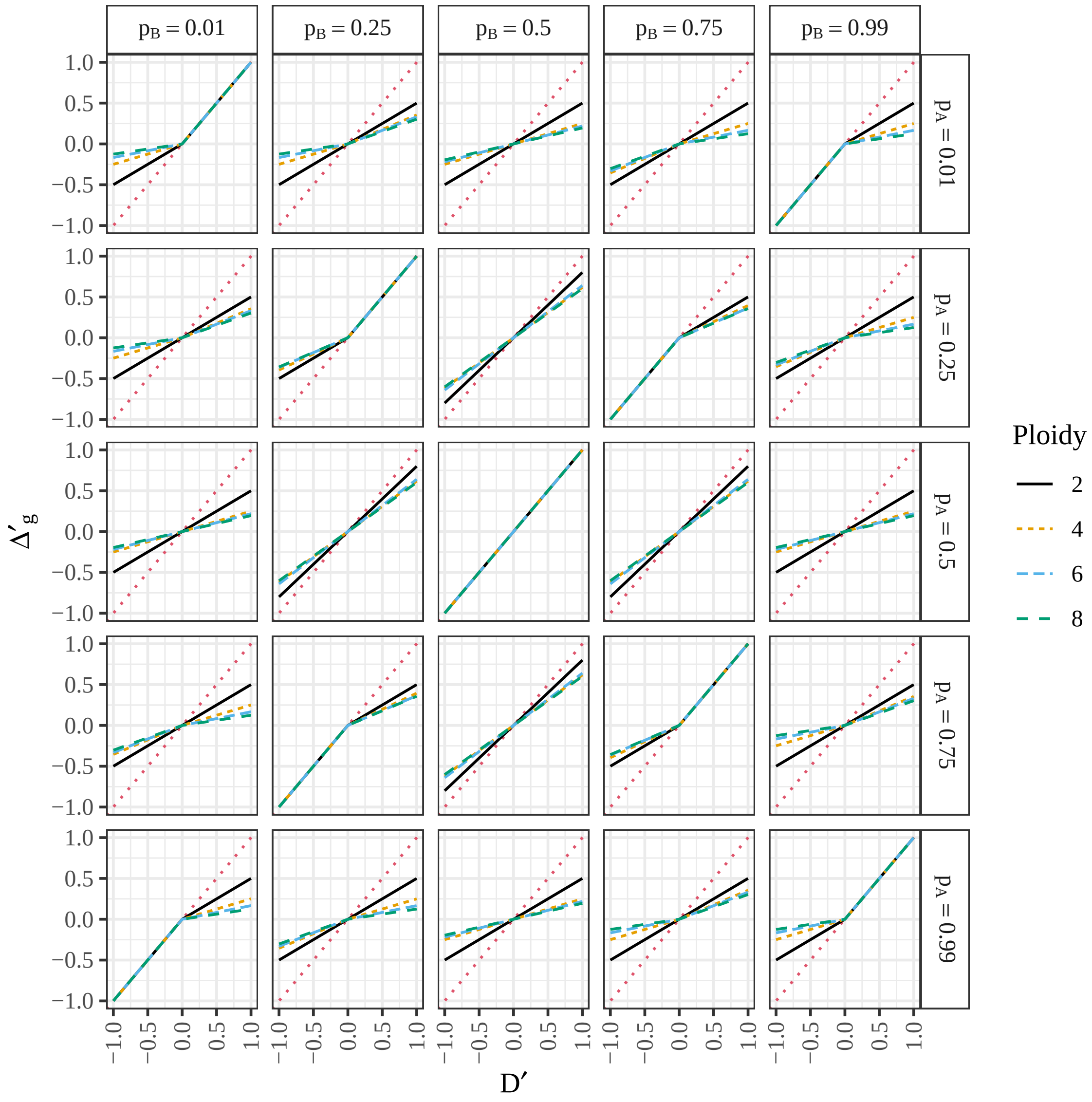


Figure S2: Relationship between $D'$ (4) ($x$-axis) and $\Delta'_g$ ($y$-axis) (17) for populations with different ploidy (color) under HWE. Row-facets index the allele frequency at the first locus and column-facets index the allele frequency at the second locus. The red dotted line is the $y = x$ line. The functional relationship appears to be piecewise linear with a change of slope at 0.

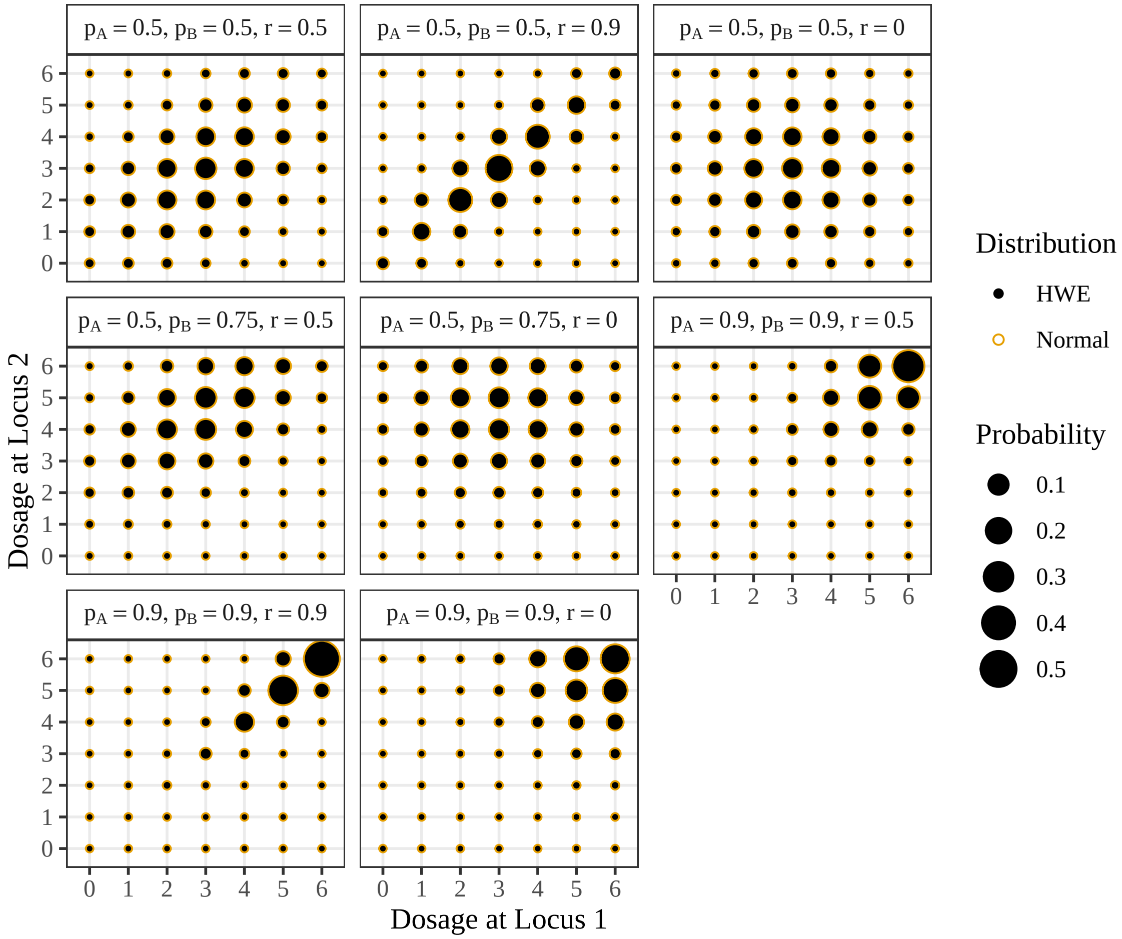


Figure S3: Joint probability distribution of two dosages. Probabilities are denoted by size. The target probability distribution is denoted by solid black circles, the closest (in Kullback-Leibler divergence) probability distribution to the target distribution among the class of proportional bivariate normal distributions is denoted by hollow orange circles. Each facet represents one of the settings of $p_A$, $p_B$, and $r$ used in the simulation study in Section 3.1.

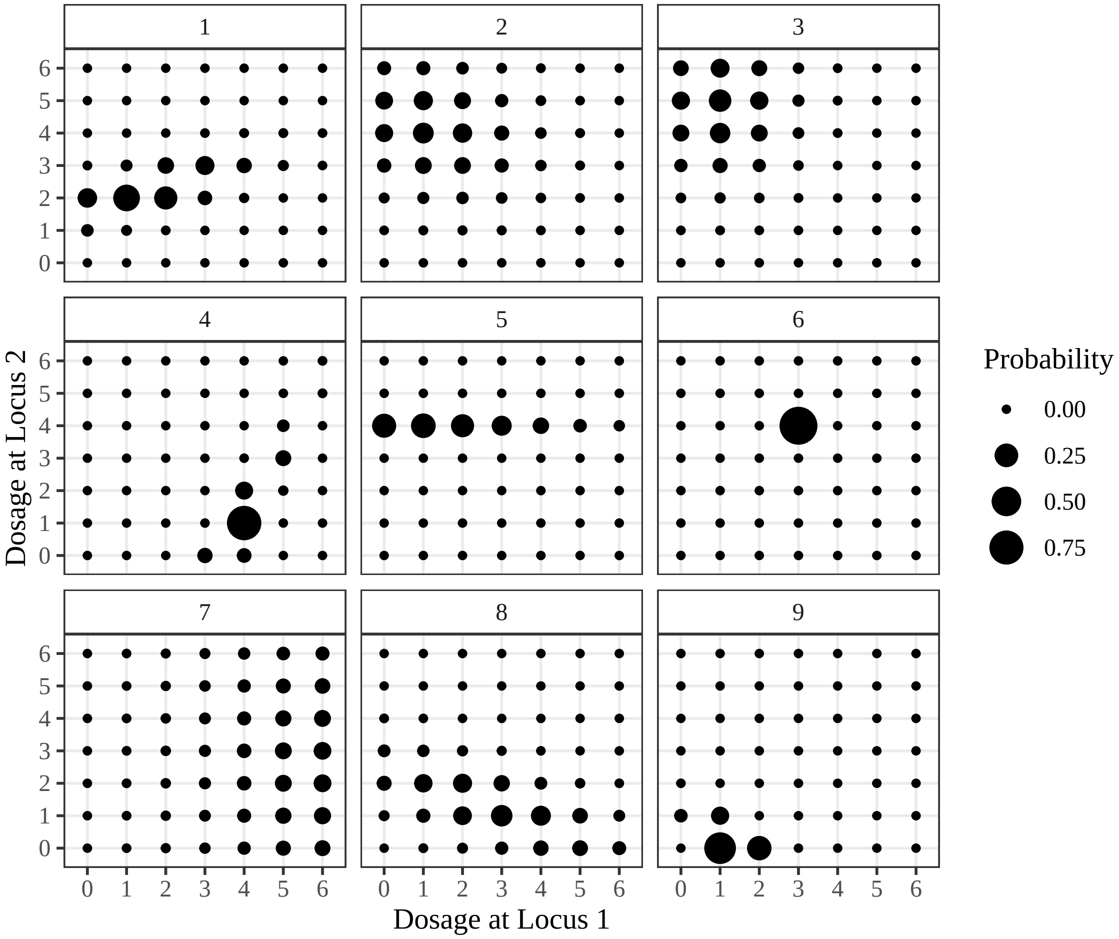


Figure S4: Joint probability distribution of two dosages. Probabilities are denoted by size. These probability distributions were chosen randomly among the class of proportional bivariate normal distributions. The proportional bivariate normal distribution can take on a variety of shapes beyond those when assuming HWE.

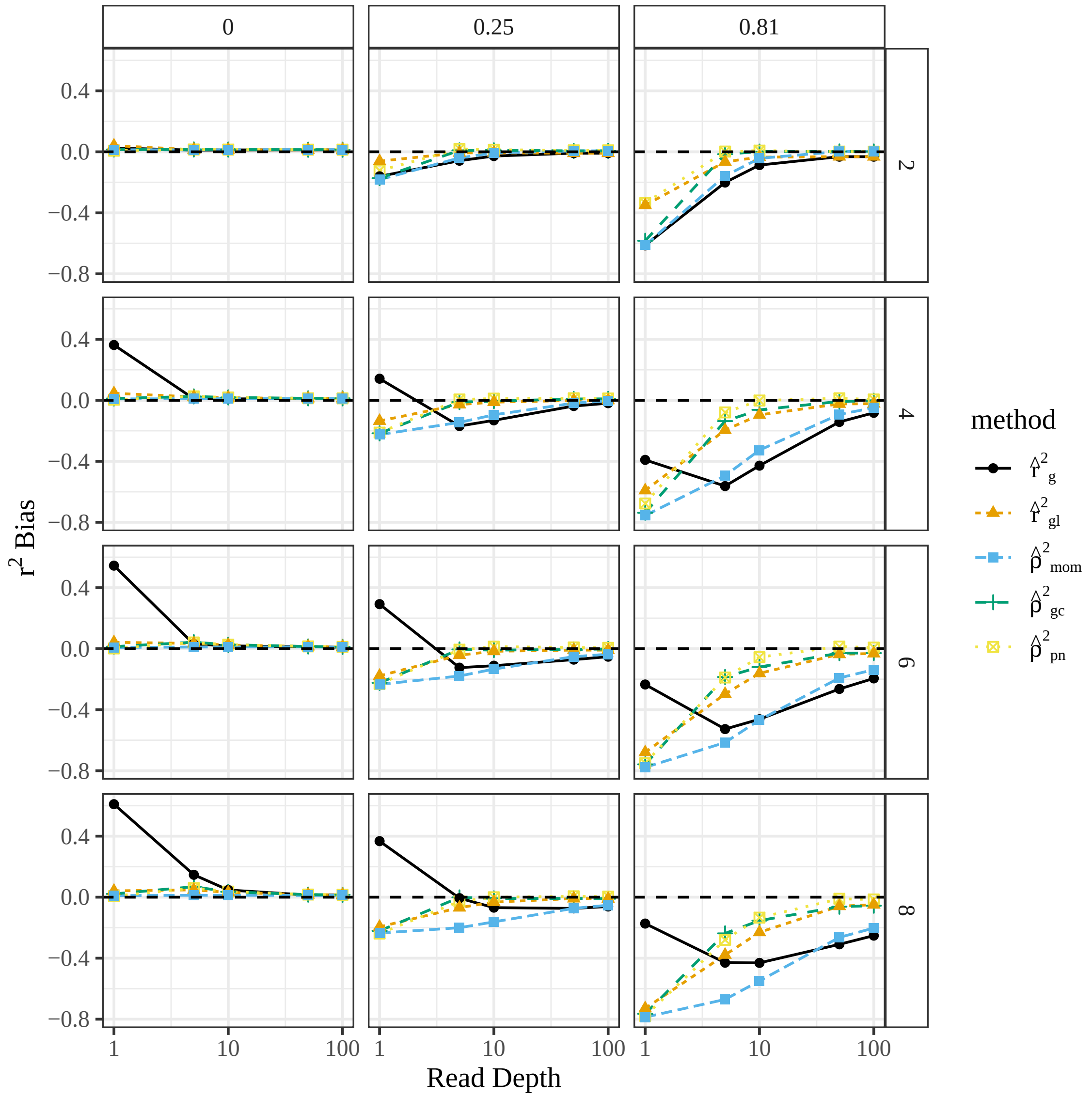


Figure S5: Bias of estimates of $r^2$ ($y$-axis) stratified by read-depth ($x$-axis), estimation method (color), ploidy (row-facets) and true $r^2$ (column-facets). The moment-based estimator has a strong bias toward zero. The MLE using genotype likelihoods is the least-biased. Simulations were performed with $p_A = 0.5$ and $p_B = 0.5$.

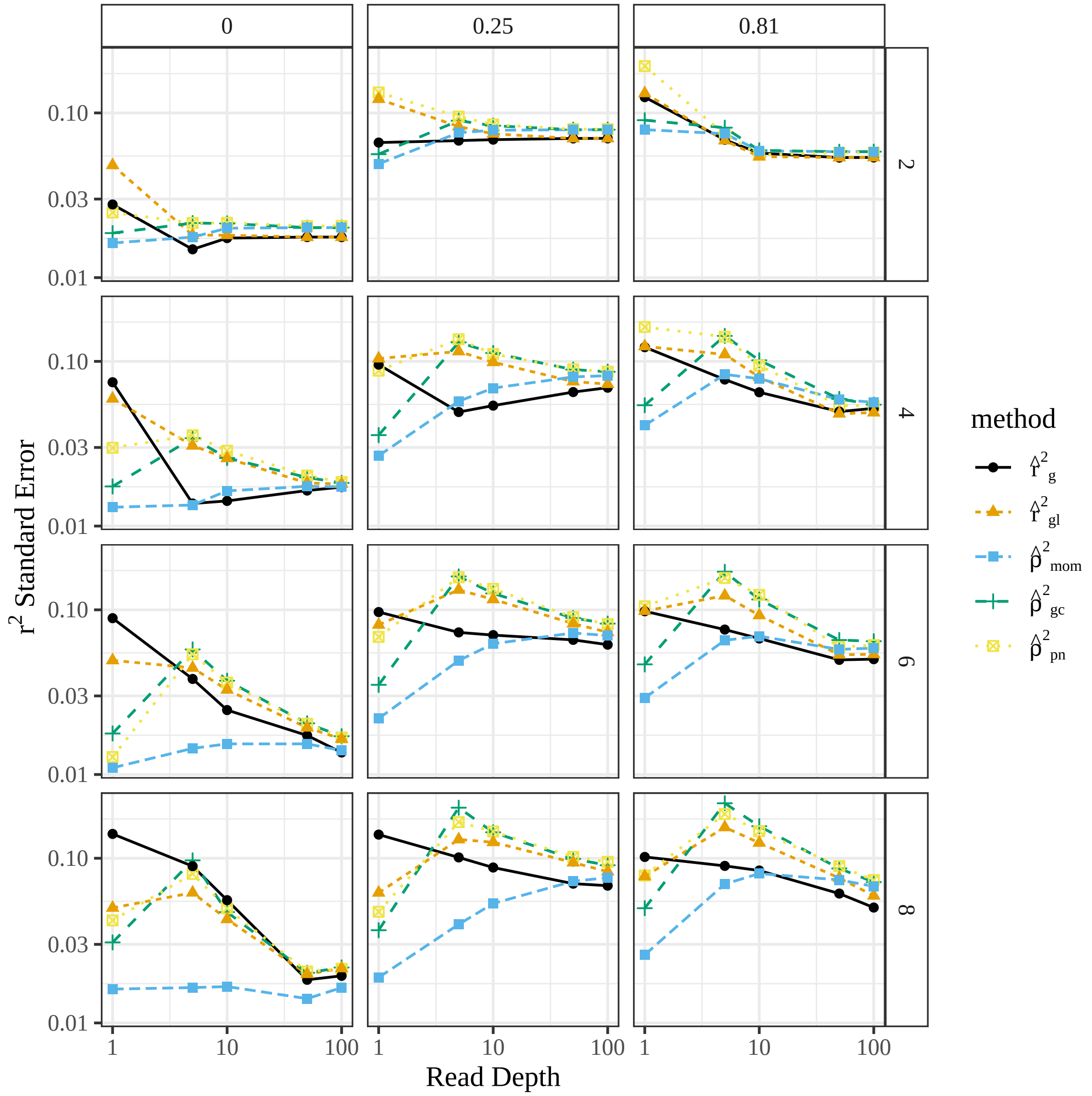


Figure S6: Standard error of $r^2$ estimators ($y$-axis) stratified by read-depth ($x$-axis), estimation method (color), ploidy (row-facets) and true $r^2$ (column-facets). Methods that use genotype likelihoods have higher standard errors. Simulations were performed with $p_A = 0.5$ and $p_B = 0.5$.

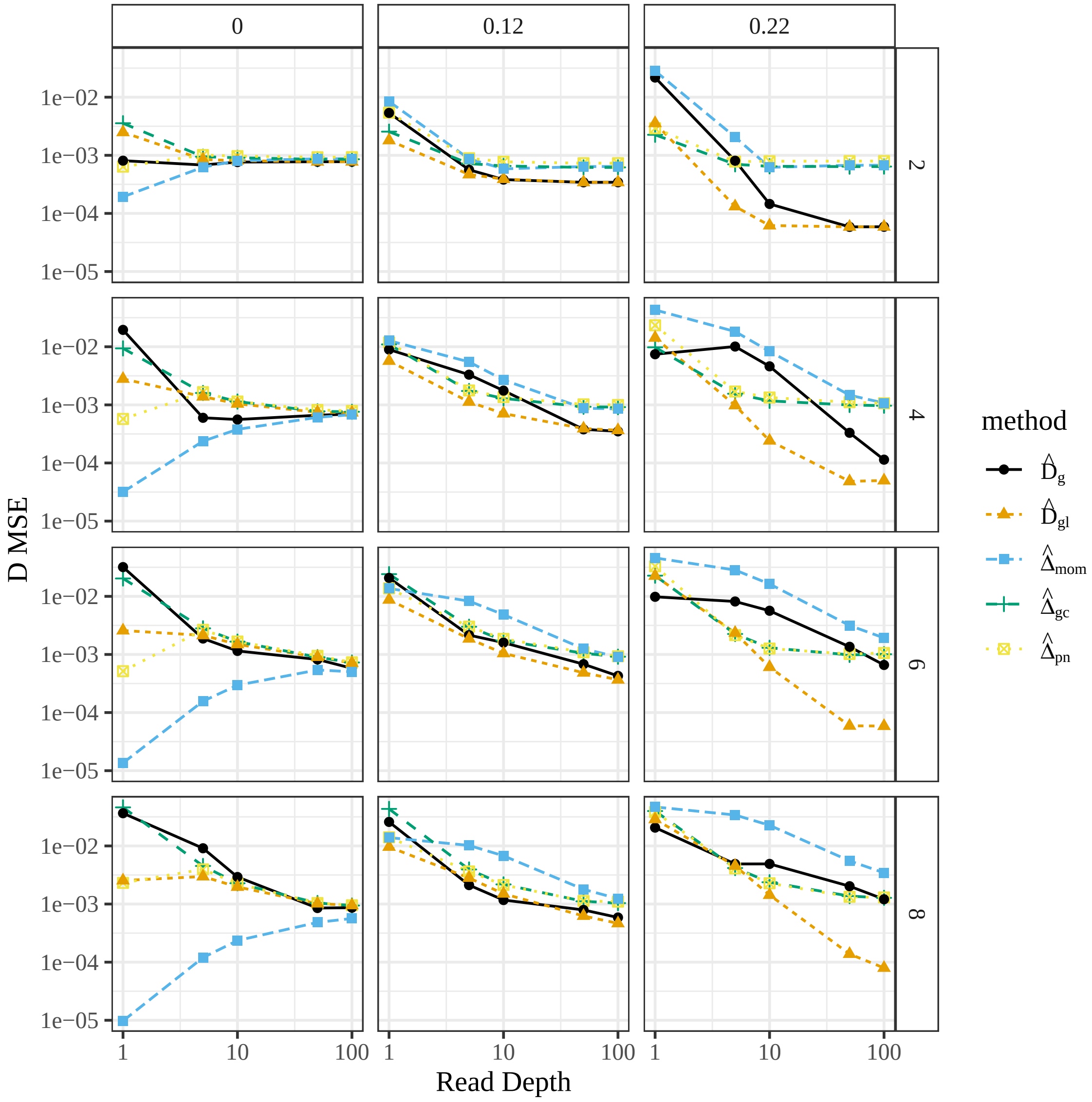


Figure S7: Mean-squared error of $D$ estimators ($y$-axis) stratified by read-depth ($x$-axis), estimation method (color), ploidy (row-facets), and true $D$ (column-facets) for the simulations from Section 3.1. Simulations were performed with $p_A = 0.5$ and $p_B = 0.5$.

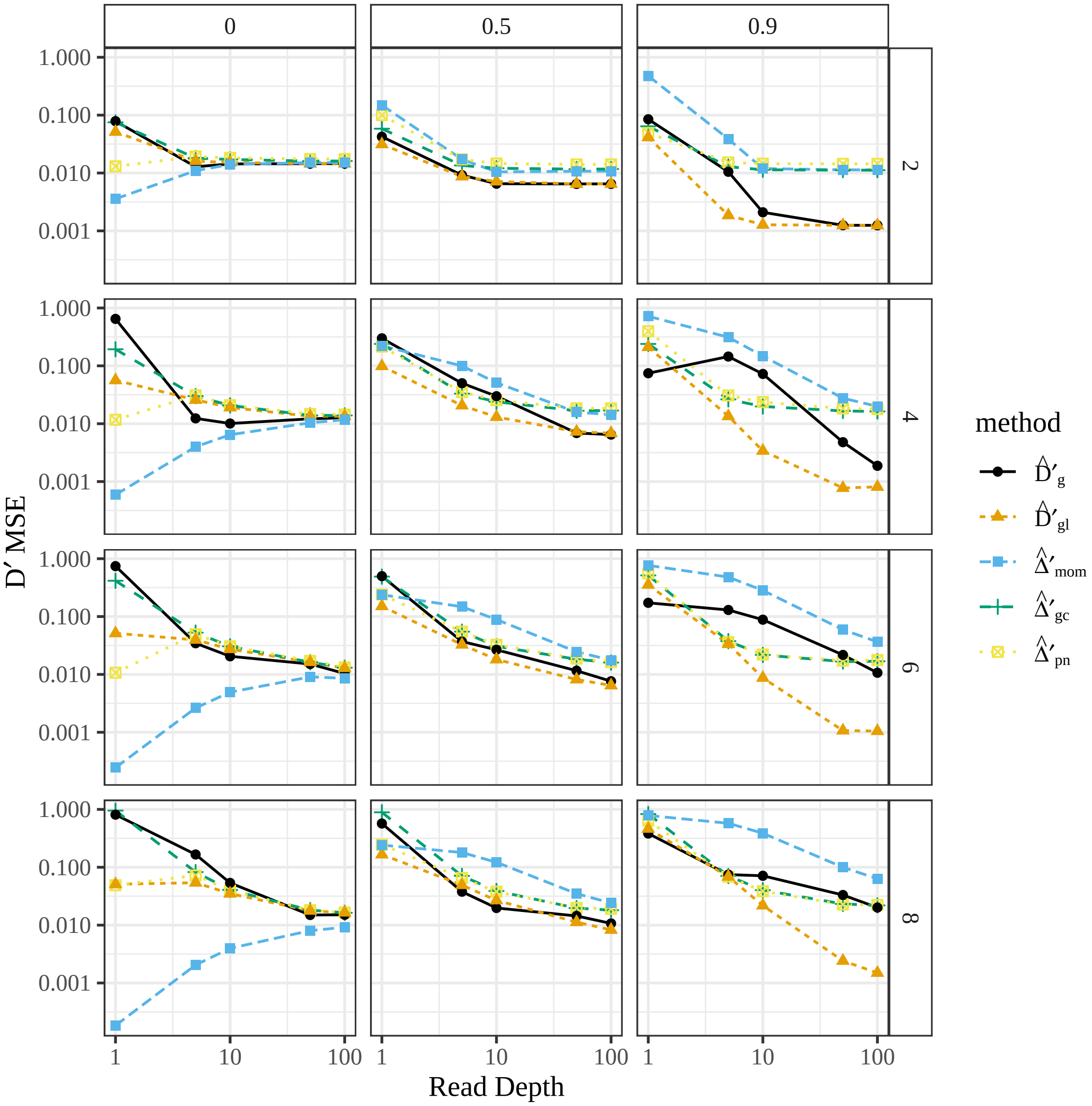


Figure S8: Mean-squared error of $D'$ estimators ($y$-axis) stratified by read-depth ($x$-axis), estimation method (color), ploidy (row-facets), and true $D'$ (column-facets) for the simulations from Section 3.1. Simulations were performed with $p_A = 0.5$ and $p_B = 0.5$.

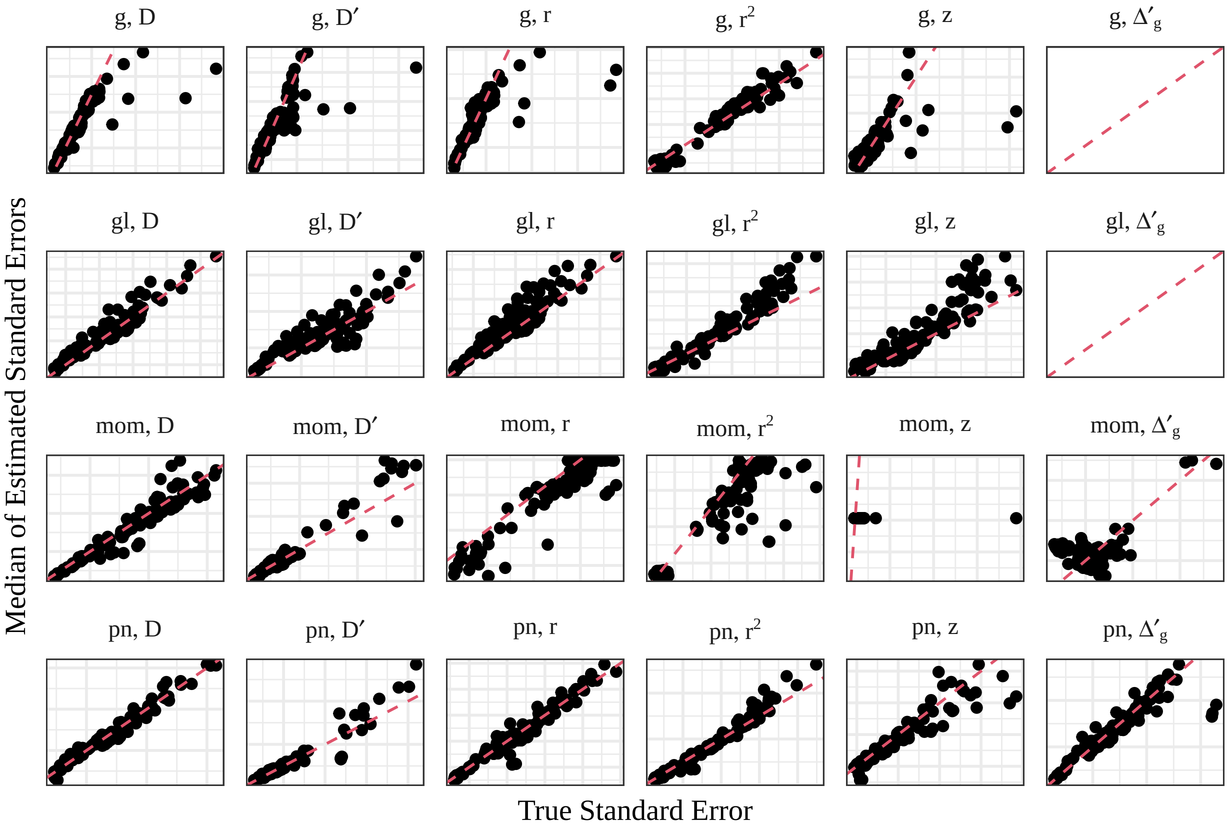


Figure S9: Standard errors of LD estimators ($x$-axis) versus the median of the estimated standard errors ($y$-axis) from the simulations in Section 3.1. Scales are freely varying between facets to allow for visualization. Each point is a different simulation setting. The scenarios where the read-depth was 1 were excluded due to poor behavior. The line is the $y = x$ line and any points above that line indicate that the estimated standard errors are typically larger than the true standard errors. Column-facets index the LD measure being estimated: $D$ (1), $D'$ (4), $r$ (6), $r^2$, $z = \text{atanh}(r)$, and $\Delta'_g$ (17). Row-facets index the estimators: "g" for $(\hat{D}_g, \hat{D}'_g, \hat{r}_g)$, "gl" for $(\hat{D}_{gl}, \hat{D}'_{gl}, \hat{r}_{gl})$, "mom" for $(\hat{\Delta}_{mom}, \hat{\Delta}'_{mom}, \hat{\rho}_{mom})$, and "pn" for $(\hat{\Delta}_{pn}, \hat{\Delta}'_{pn}, \hat{\rho}_{pn})$. Two empty facets are present because haplotypic methods cannot estimate $\Delta'_g$, a purely composite measure.

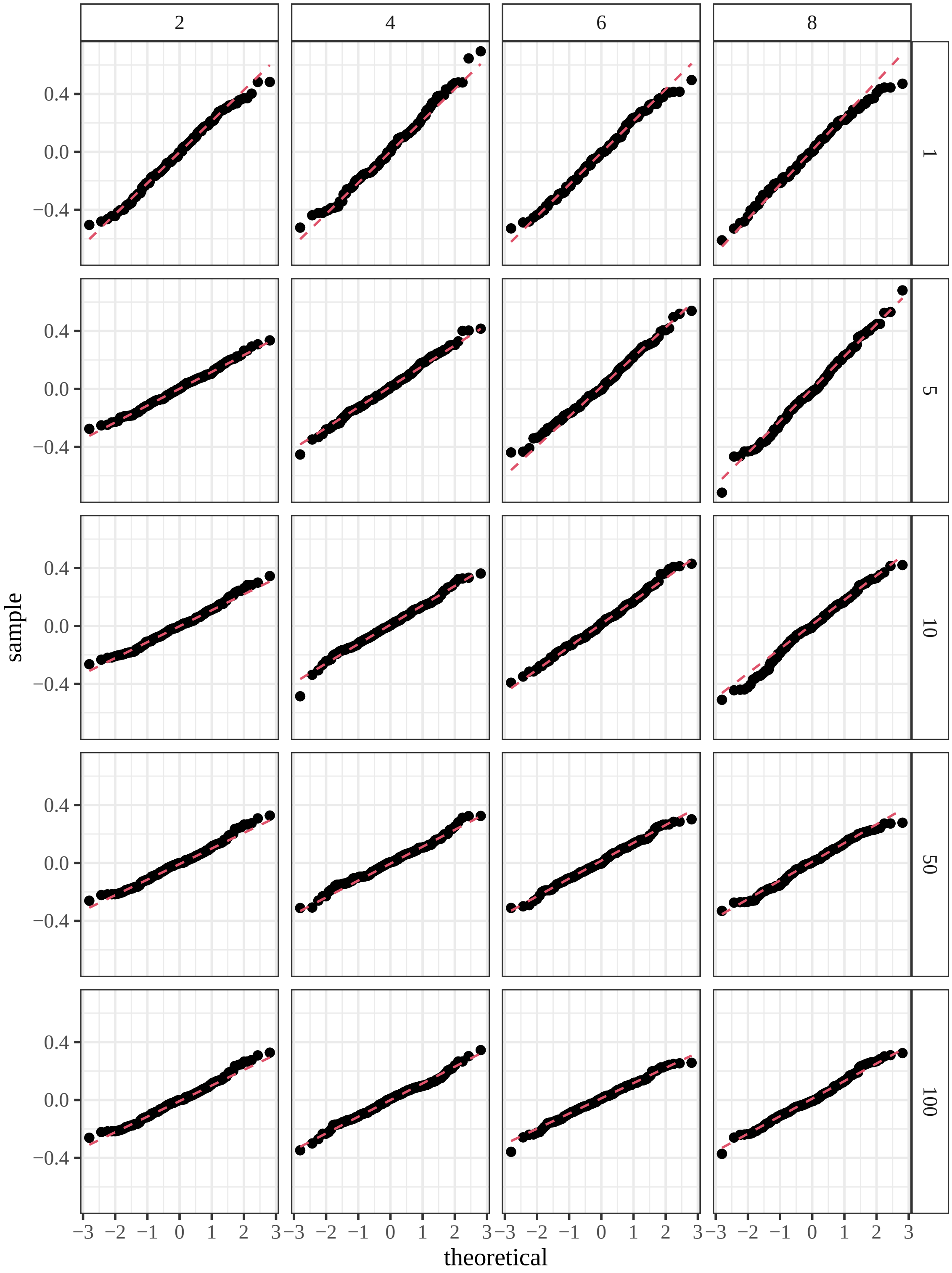


Figure S10: QQ-plots of of the Fisher-$z$ transformation of $\hat{r}_{gl}$ when $p_A = 0.5$ and $p_B = 0.5$ from the simulations in Section 3.1. The estimates appear to approximately follow a normal distribution.

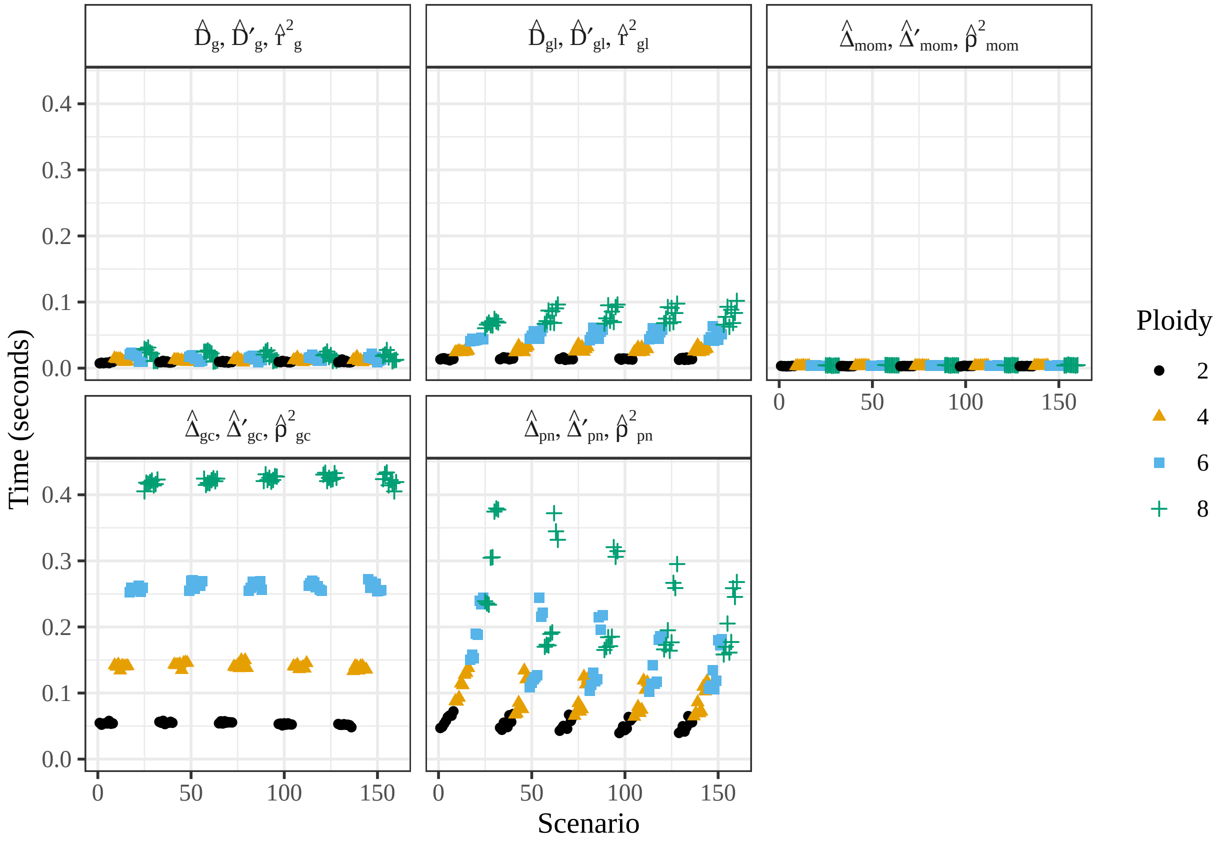


Figure S11: Mean computation time in seconds ($y$-axis) for each method (facets) stratified by the simulation settings ($x$-axis) for the simulations in Section 3.1. Methods using genotype likelihoods are generally slower, but all methods take less than half a second on average.

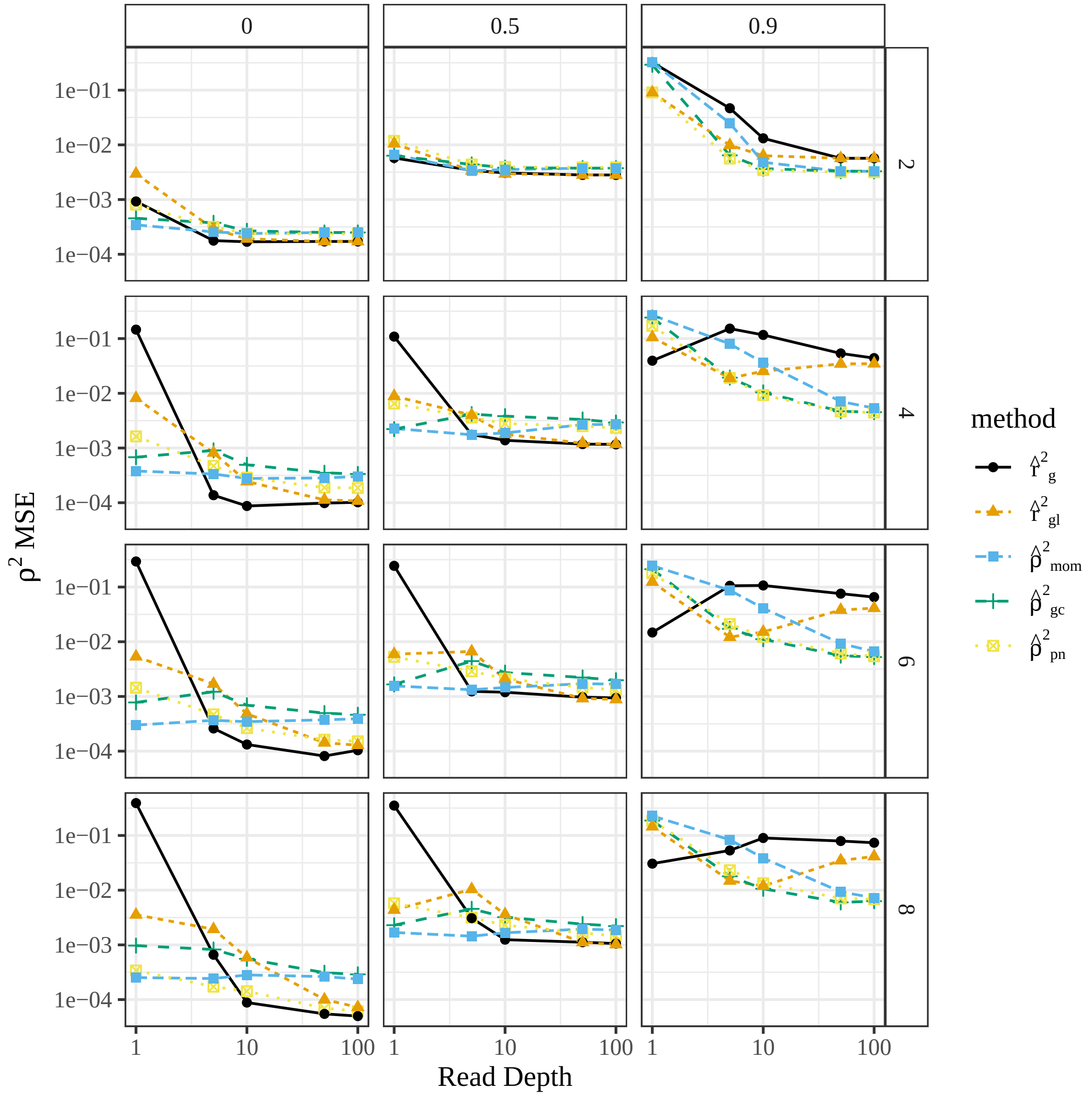


Figure S12: Average mean-squared error ($y$-axis) stratified by read-depth ($x$-axis), ploidy (row-facets), and association parameter of the proportional bivariate normal distribution (column-facets) for the simulations from Section S11.

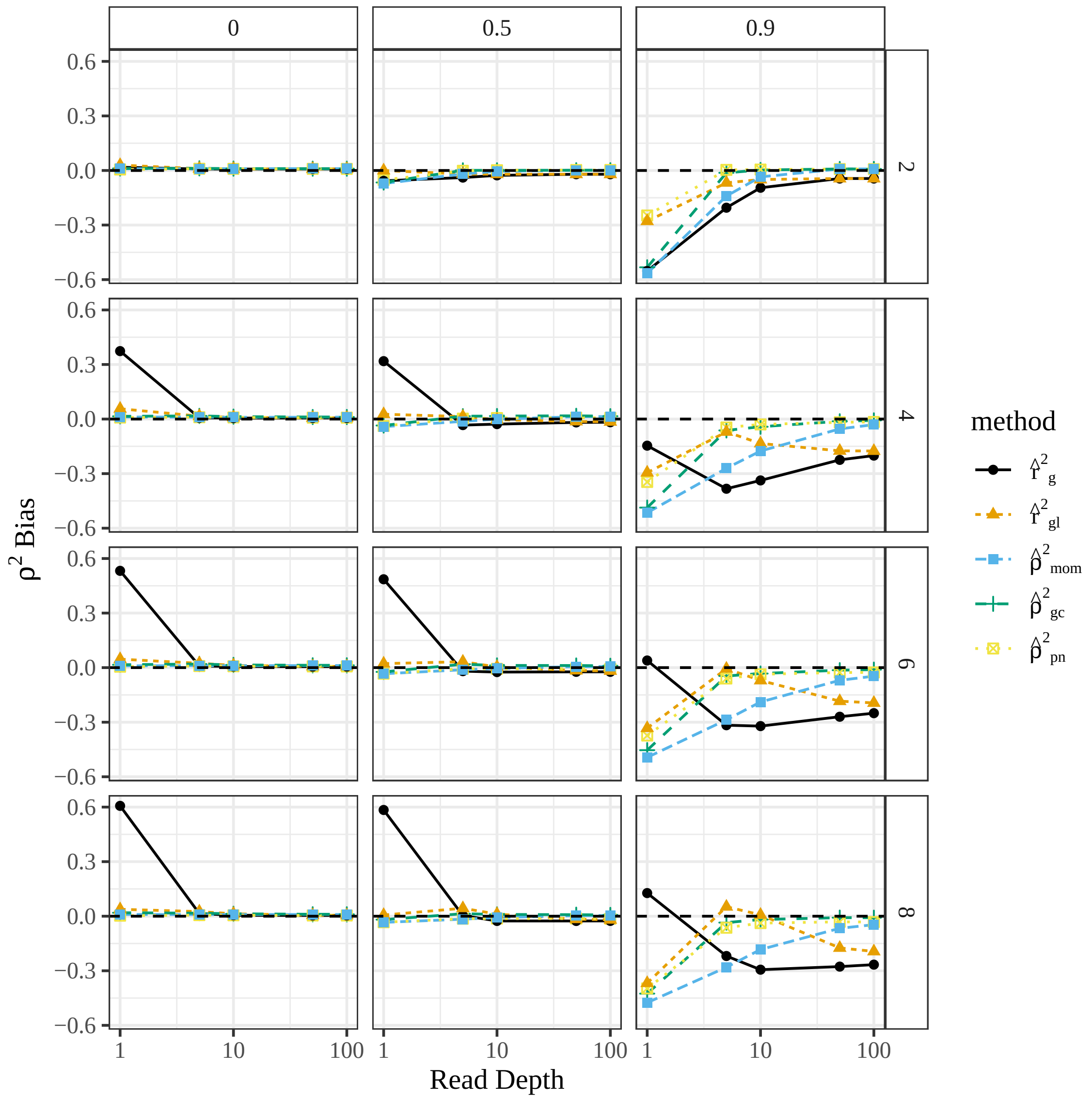


Figure S13: Average bias ($y$-axis) stratified by read-depth ($x$-axis), ploidy (row-facets) and association parameter of the proportional bivariate normal distribution (column-facets).

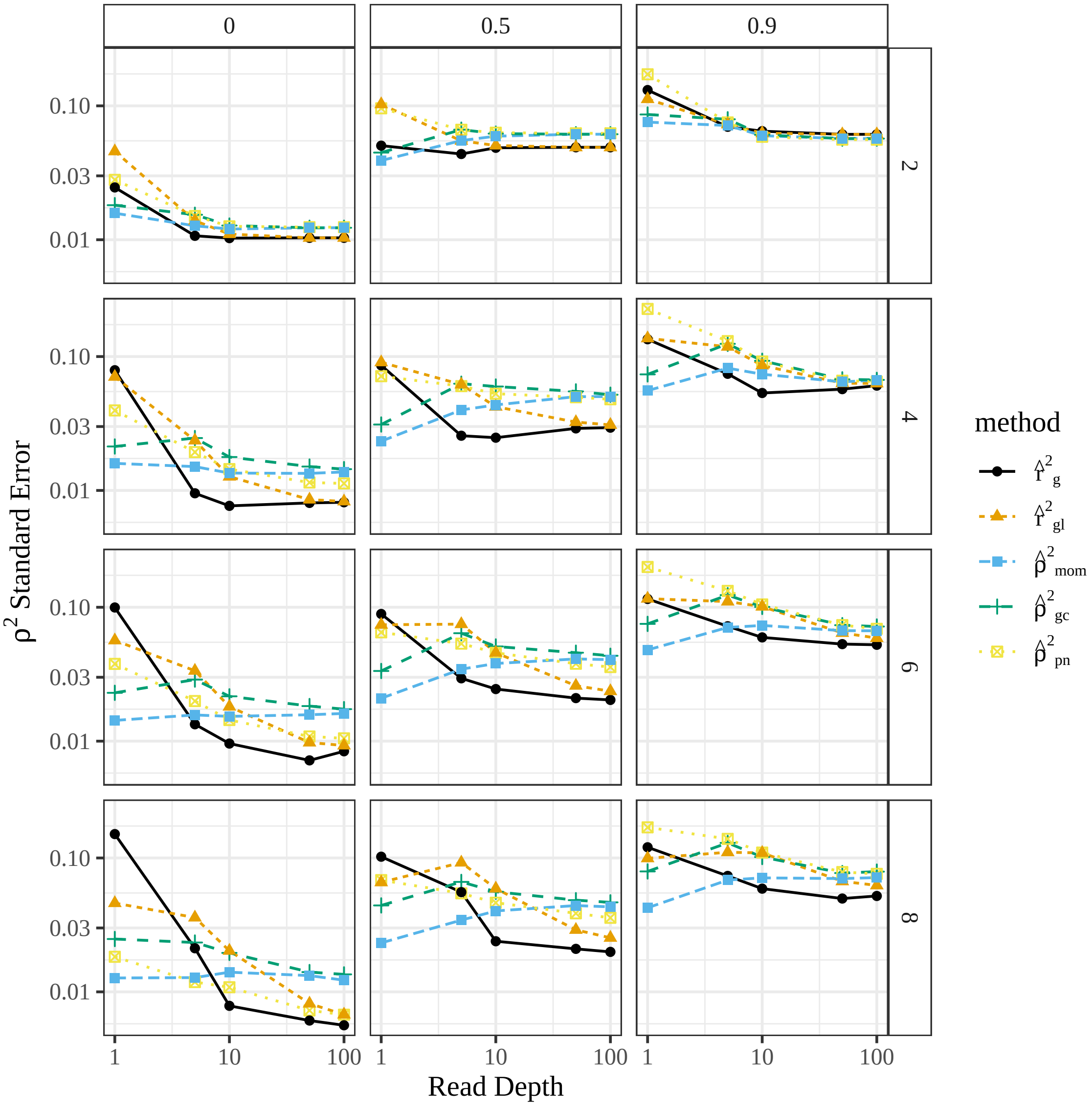


Figure S14: Average standard error ($y$-axis) stratified by read-depth ($x$-axis), ploidy (row-facets) and association parameter of the proportional bivariate normal distribution (column-facets).

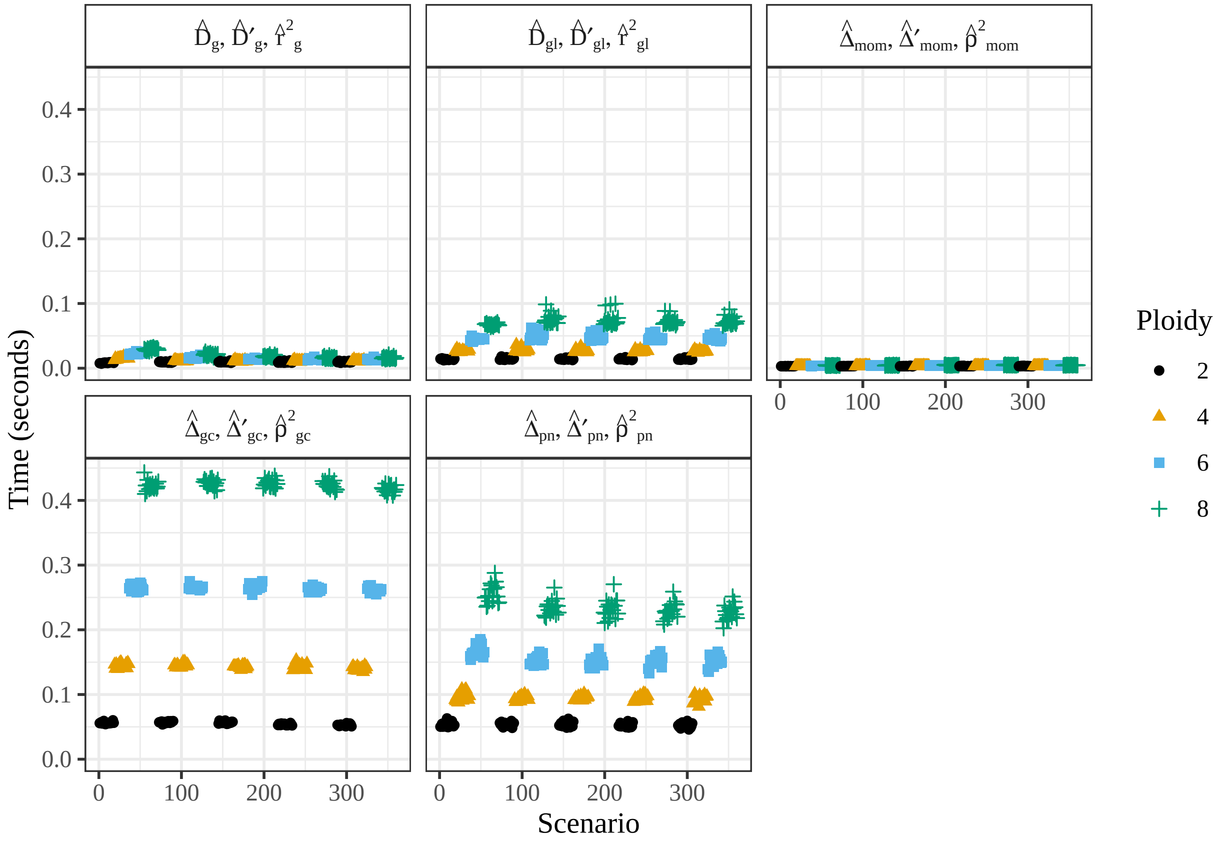


Figure S15: Mean computation time in seconds ($y$-axis) for each method (facets) stratified by the simulation settings ($x$-axis) for the simulations in Section S11. Methods using genotype likelihoods are generally slower, but all methods take less than half a second on average.

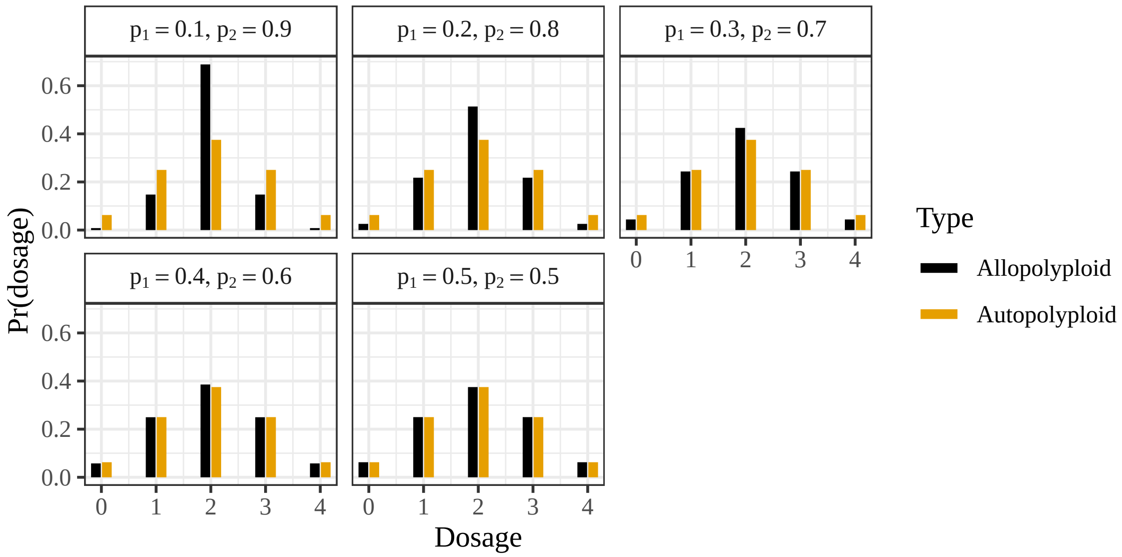


Figure S16: Comparing genotype distributions of autopolyploids under HWE with an allele frequency of 0.5 with allopolyploids under HWE with allele frequencies of $p_1$ and $p_2$ on their subgenomes. The distributions are equivalent at $p_1 = p_2 = 0.5$, but other levels of $p_1$ and $p_2$ result in underdispersion for the allopolyploid genotype distributions.

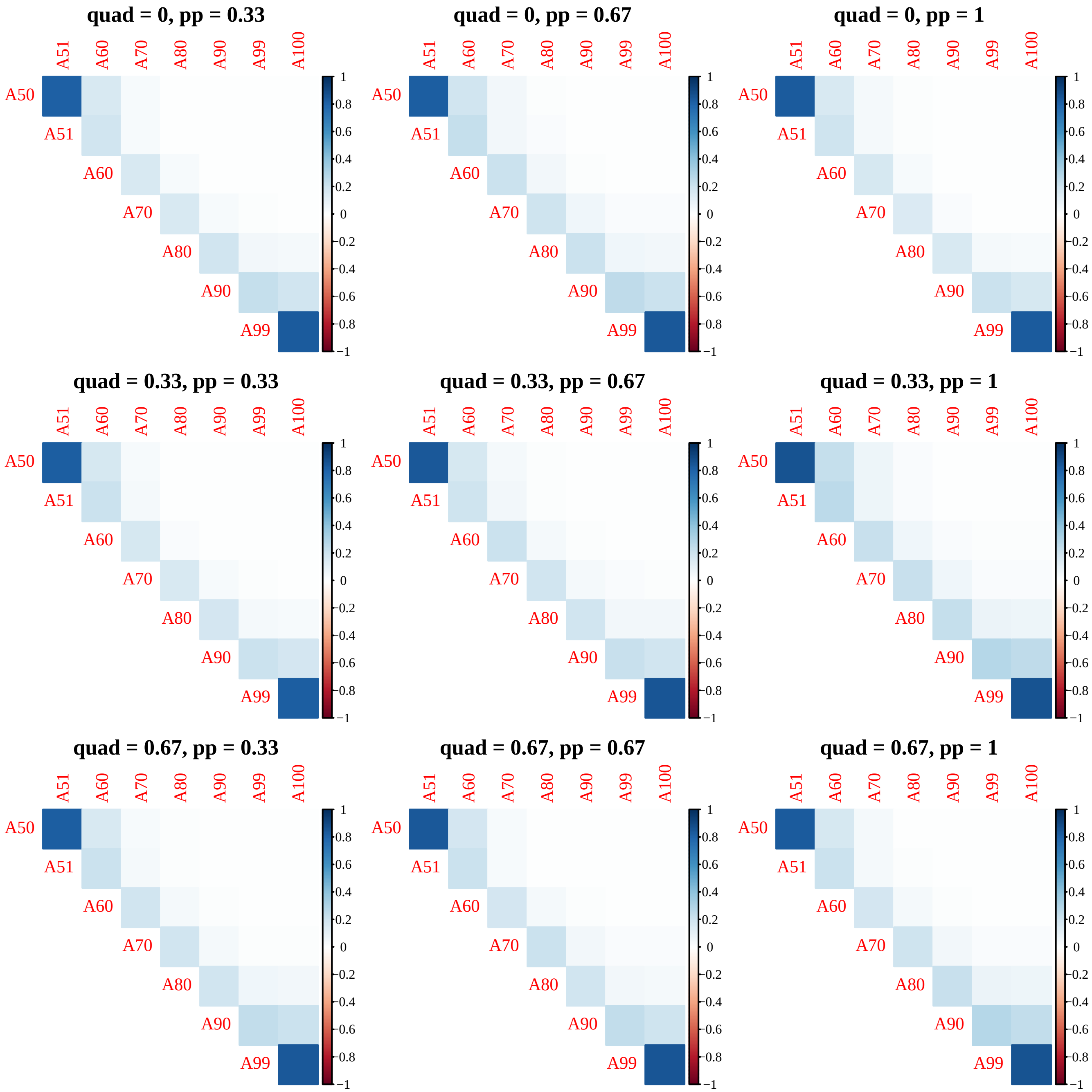


Figure S17: Heatmap of pairwise $\rho^2$ between SNPs at 50 cM, 51 cM, 60 cM, 70 cM, 80 cM, 90 cM, 99 cM, and 100 cM, where the centromere is located at 50 cM, using 10000 individuals simulated from 10 generations of random mating using PedigreeSim under different levels of quadrivalent formation (row facets) and different levels of homologous pairing (column facets).

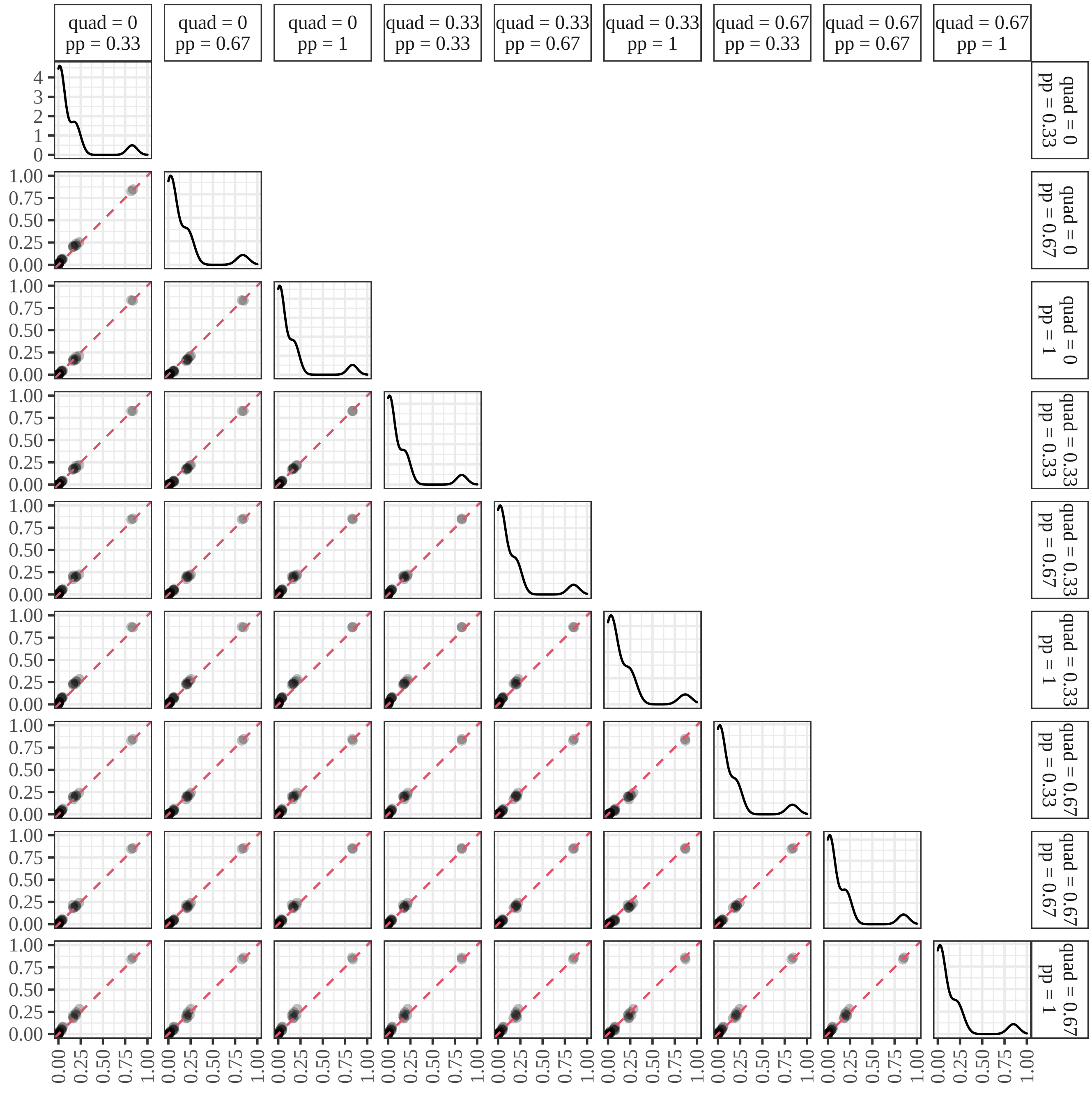


Figure S18: Pairs plot of pairwise $\rho^2$ between SNPs at 50 cM, 51 cM, 60 cM, 70 cM, 80 cM, 90 cM, 99 cM, and 100 cM, where the centromere is located at 50 cM, using 10000 individuals simulated from 10 generations of random mating using PedigreeSim under different levels of quadrivalent formation ("quad") and different levels of homologous pairing ("pp").

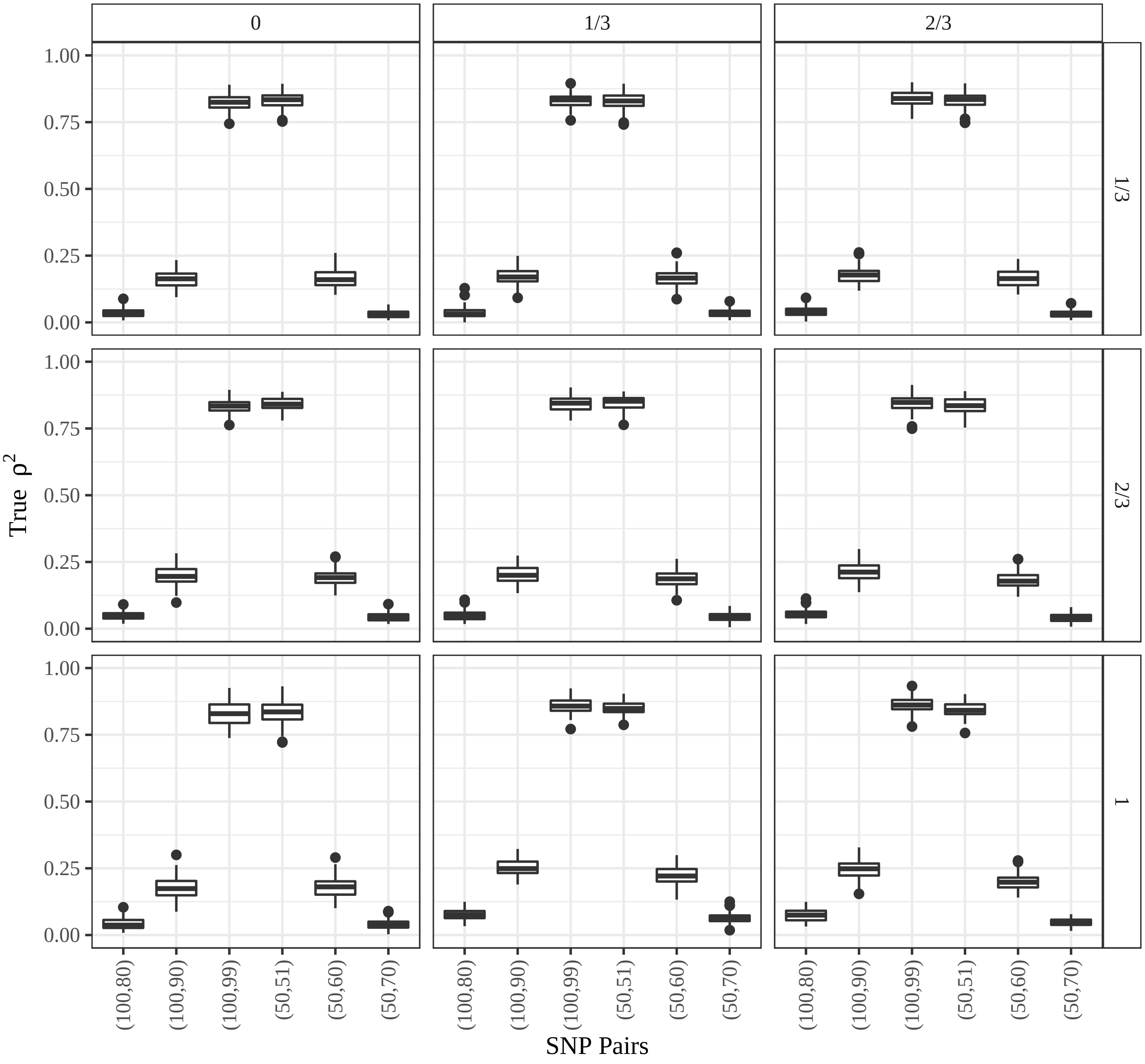


Figure S19: Boxplots of the true $\rho^2$ ($y$-axis) for pairs of SNPs ($x$-axis) under different levels of homologous pairing (row-facets) and quadrivalent formation (column facets). SNP pairs are labeled based on their location on the chromosome in centimorgans. Data were generated from 10 generations of random mating of a tetraploid species, where the founder population began at an allele frequency of 0.5, subgenome allele frequencies of 0.1 and 0.9, and an initial $\rho^2$ of 1. Populations were kept at a constant size of 1000 individuals. The "true" $\rho^2$ values were calcuated using all 1000 individuals from the final generation.

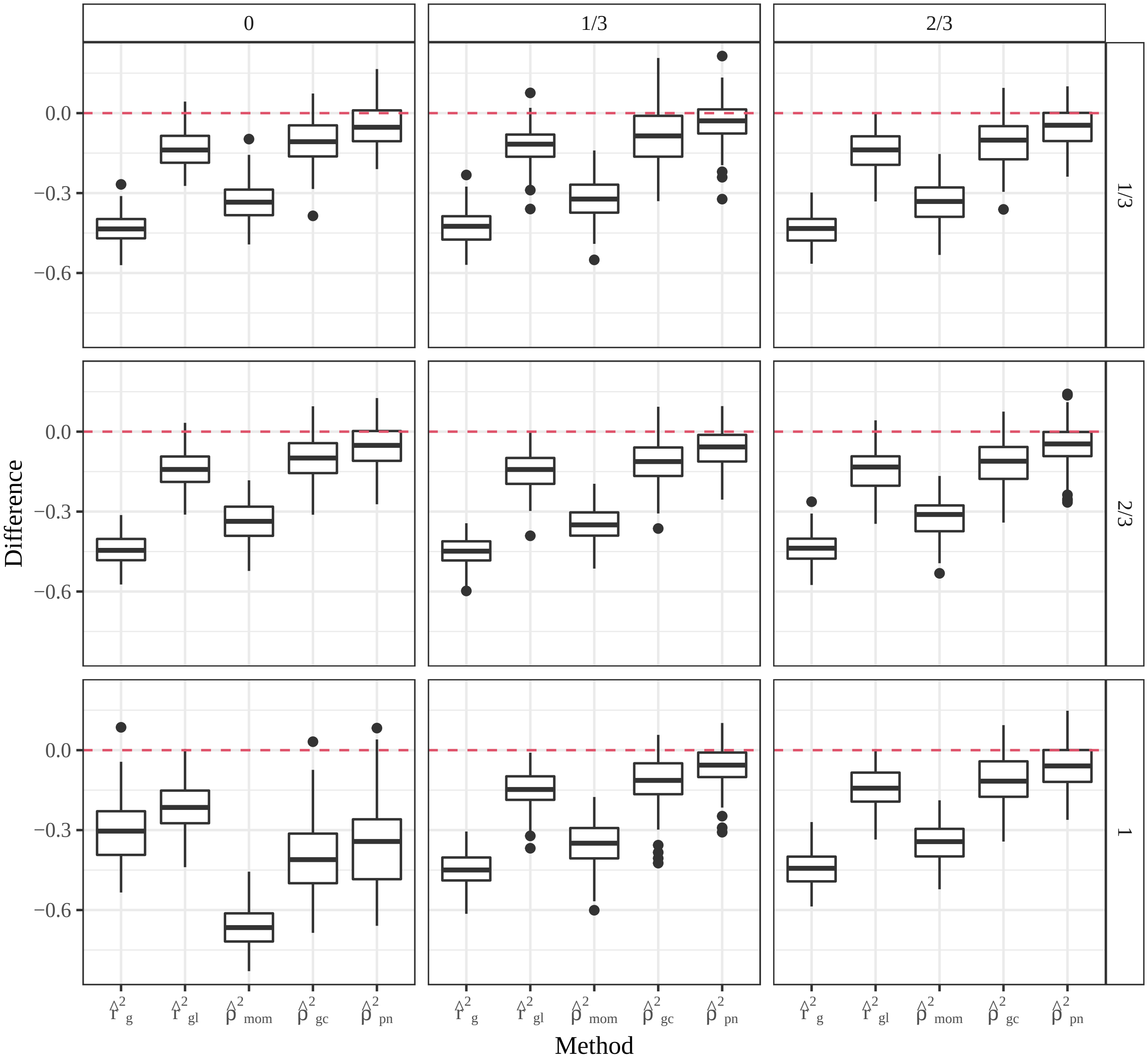


Figure S20: Boxplots of the difference between estimated $\rho^2$ and true $\rho^2$ ($y$-axis) for each method ($x$-axis) between two SNPs were located at 50 cM and 51 cM, and so should exibit large levels of LD (Figures S17 and S19). Values near zero (the dashed line) indicate unbiased behavior, while values over the dashed line indicate overestimation of $\rho^2$. Data were generated from 10 generations of random mating of a tetraploid species, where the founder population began at an allele frequency of 0.5, subgenome allele frequencies of 0.1 and 0.9, and an initial $\rho^2$ of 1. Populations were kept at a constant size of 1000 individuals. Different levels of homologous pairing (row-facets) and quadrivalent formation (column facets) were explored. "True" $\rho^2$ was calculated using all 1000 individuals from the final generation, and the estimated $\rho^2$ was calculated using a random sample of 100 individuals. The centromere is located at 50 cM.

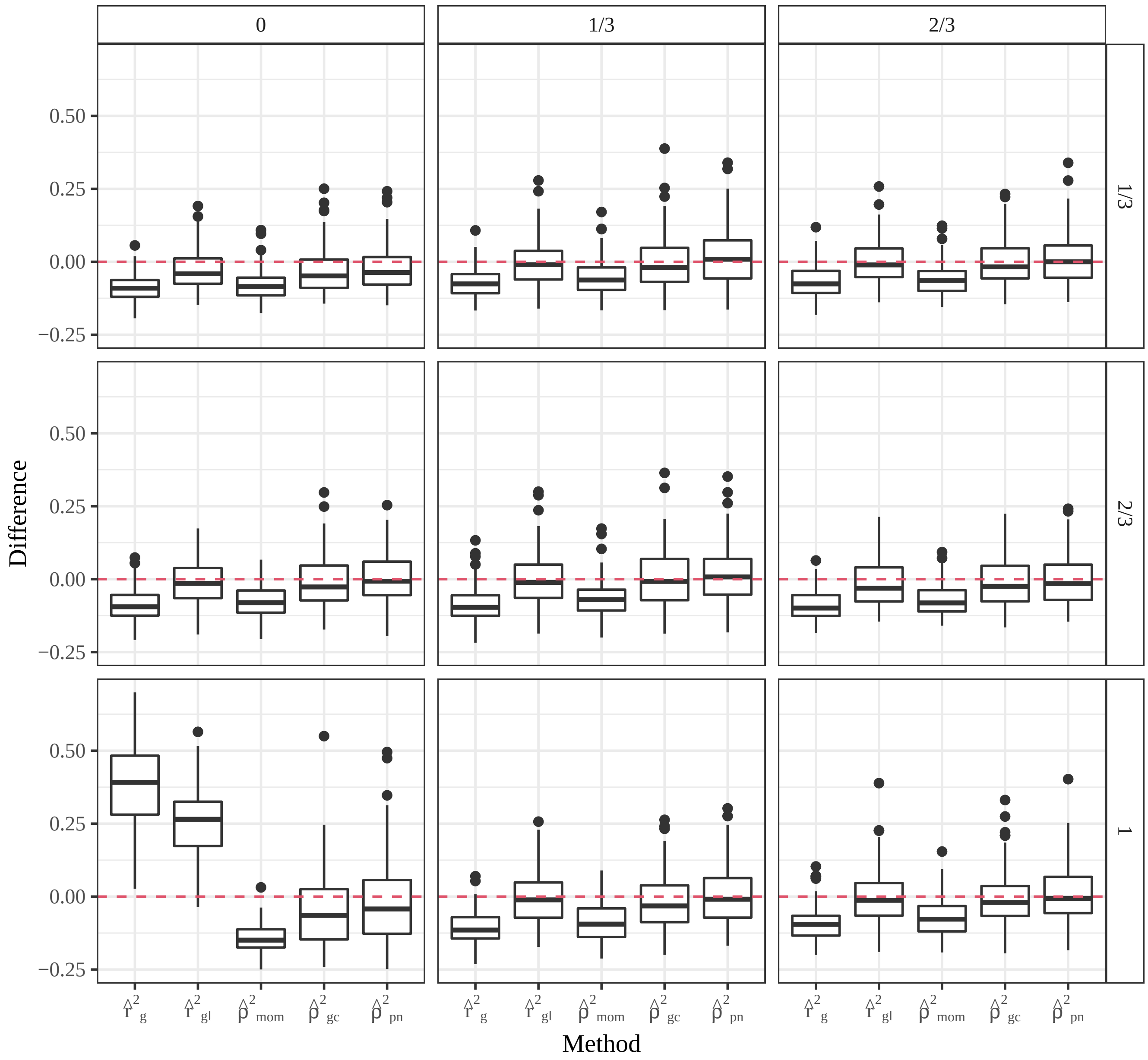


Figure S21: Boxplots of the difference between estimated $\rho^2$ and true $\rho^2$ ($y$-axis) for each method ($x$-axis) between two SNPs were located at 50 cM and 60 cM, and so should exibit moderate levels of LD (Figures S17 and S19). Values near zero (the dashed line) indicate unbiased behavior, while values over the dashed line indicate overestimation of $\rho^2$. Data were generated from 10 generations of random mating of a tetraploid species, where the founder population began at an allele frequency of 0.5, subgenome allele frequencies of 0.1 and 0.9, and an initial $\rho^2$ of 1. Populations were kept at a constant size of 1000 individuals. Different levels of homologous pairing (row-facets) and quadrivalent formation (column facets) were explored. "True" $\rho^2$ was calculated using all 1000 individuals from the final generation, and the estimated $\rho^2$ was calculated using a random sample of 100 individuals. The centromere is located at 50 cM.

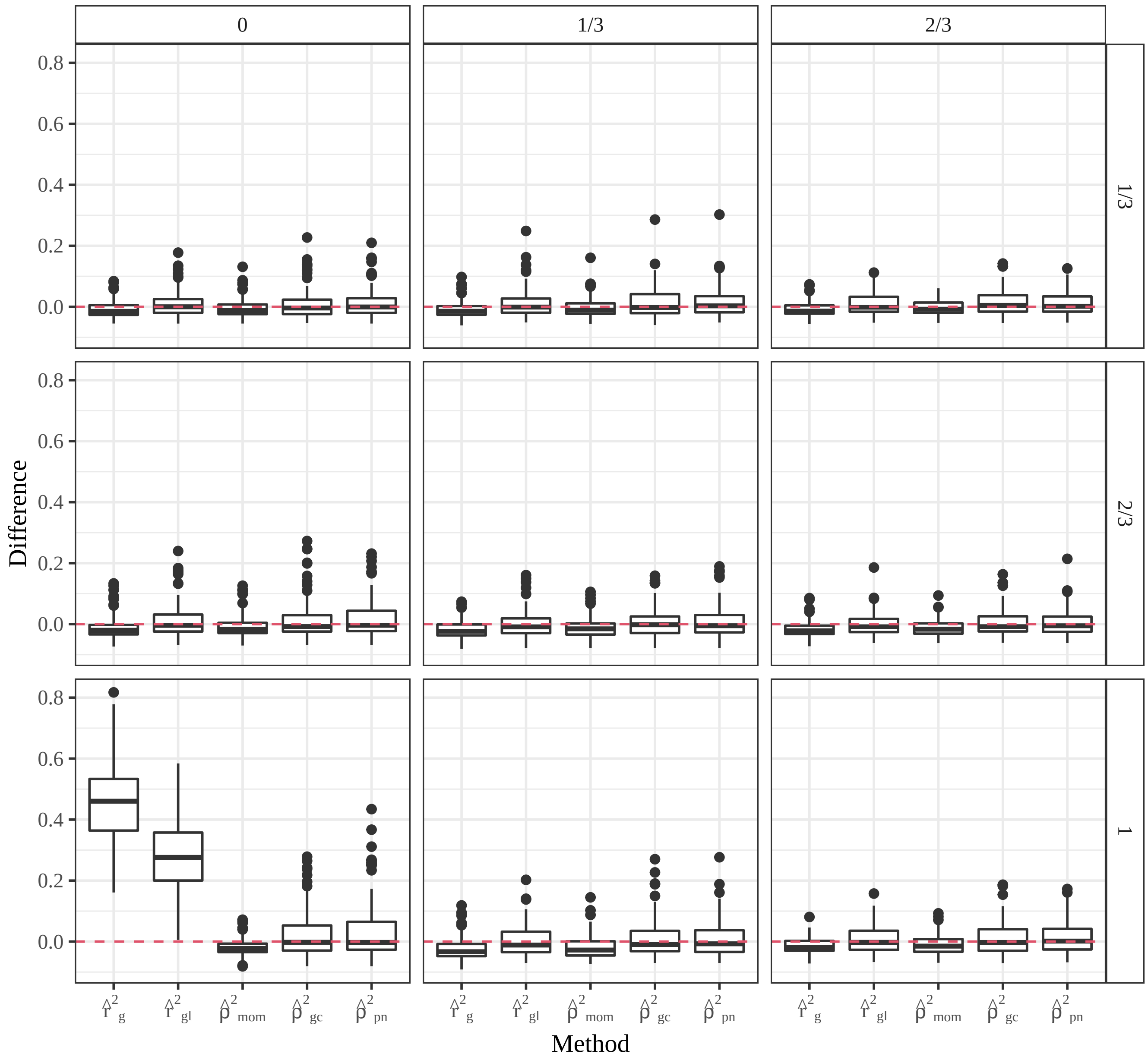


Figure S22: Boxplots of the difference between estimated $\rho^2$ and true $\rho^2$ ($y$-axis) for each method ($x$-axis) between two SNPs were located at 50 cM and 70 cM, and so should exibit LD close to 0 (Figures S17 and S19). Values near zero (the dashed line) indicate unbiased behavior, while values over the dashed line indicate overestimation of $\rho^2$. Data were generated from 10 generations of random mating of a tetraploid species, where the founder population began at an allele frequency of 0.5, subgenome allele frequencies of 0.1 and 0.9, and an initial $\rho^2$ of 1. Populations were kept at a constant size of 1000 individuals from the final generation. Different levels of homologous pairing (row-facets) and quadrivalent formation (column facets) were explored. "True" $\rho^2$ was calculated using all 1000 individuals, and the estimated $\rho^2$ was calculated using a random sample of 100 individuals. The centromere is located at 50 cM.

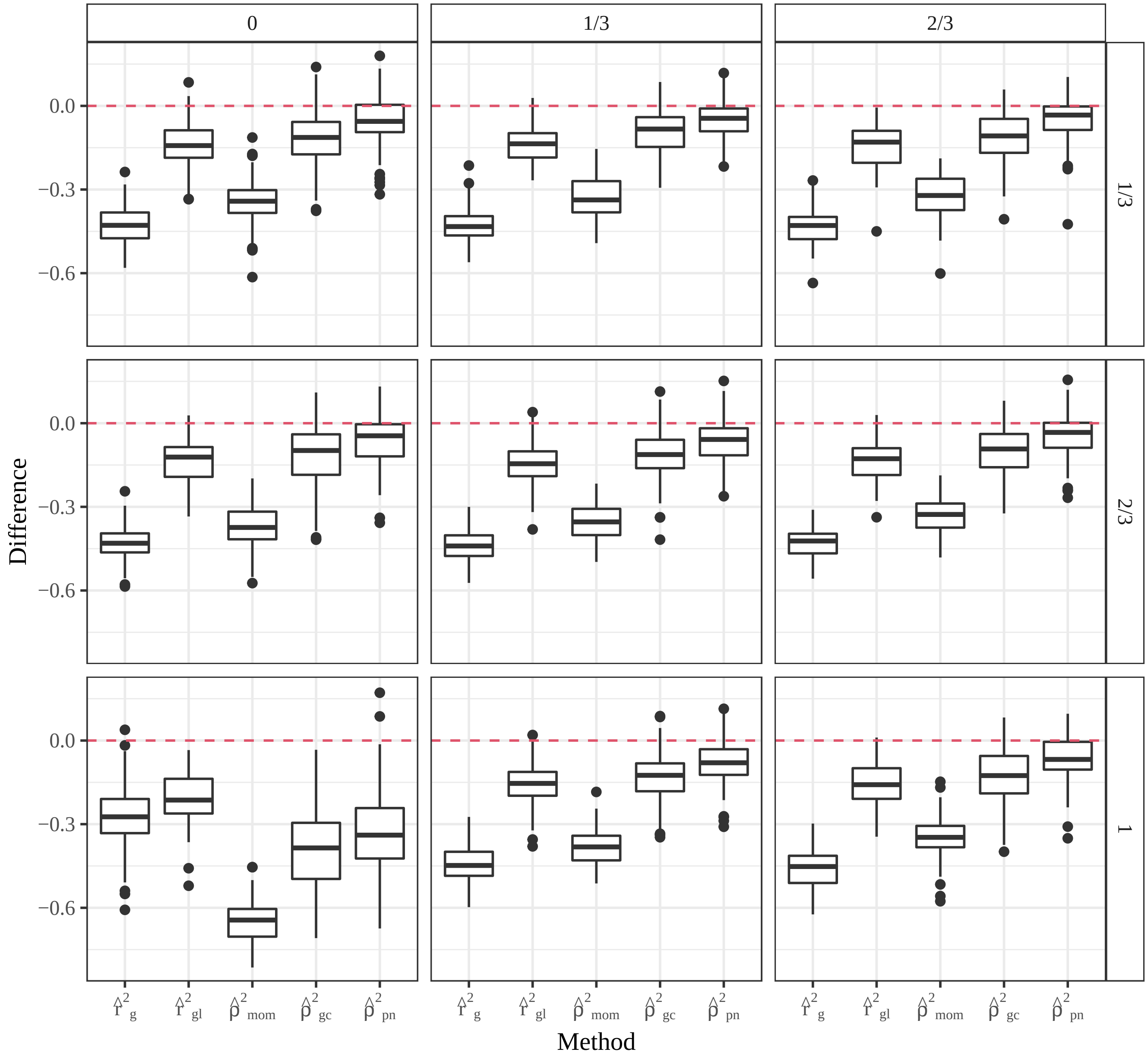


Figure S23: Boxplots of the difference between estimated $\rho^2$ and true $\rho^2$ ($y$-axis) for each method ($x$-axis) between two SNPs were located at 100 cM and 99 cM, and so should exibit large levels of LD (Figures S17 and S19). Values near zero (the dashed line) indicate unbiased behavior, while values over the dashed line indicate overestimation of $\rho^2$. Data were generated from 10 generations of random mating of a tetraploid species, where the founder population began at an allele frequency of 0.5, subgenome allele frequencies of 0.1 and 0.9, and an initial $\rho^2$ of 1. Populations were kept at a constant size of 1000 individuals from the final generation. Different levels of homologous pairing (row-facets) and quadrivalent formation (column facets) were explored. “True” $\rho^2$ was calculated using all 1000 individuals, and the estimated $\rho^2$ was calculated using a random sample of 100 individuals. The centromere is located at 50 cM.

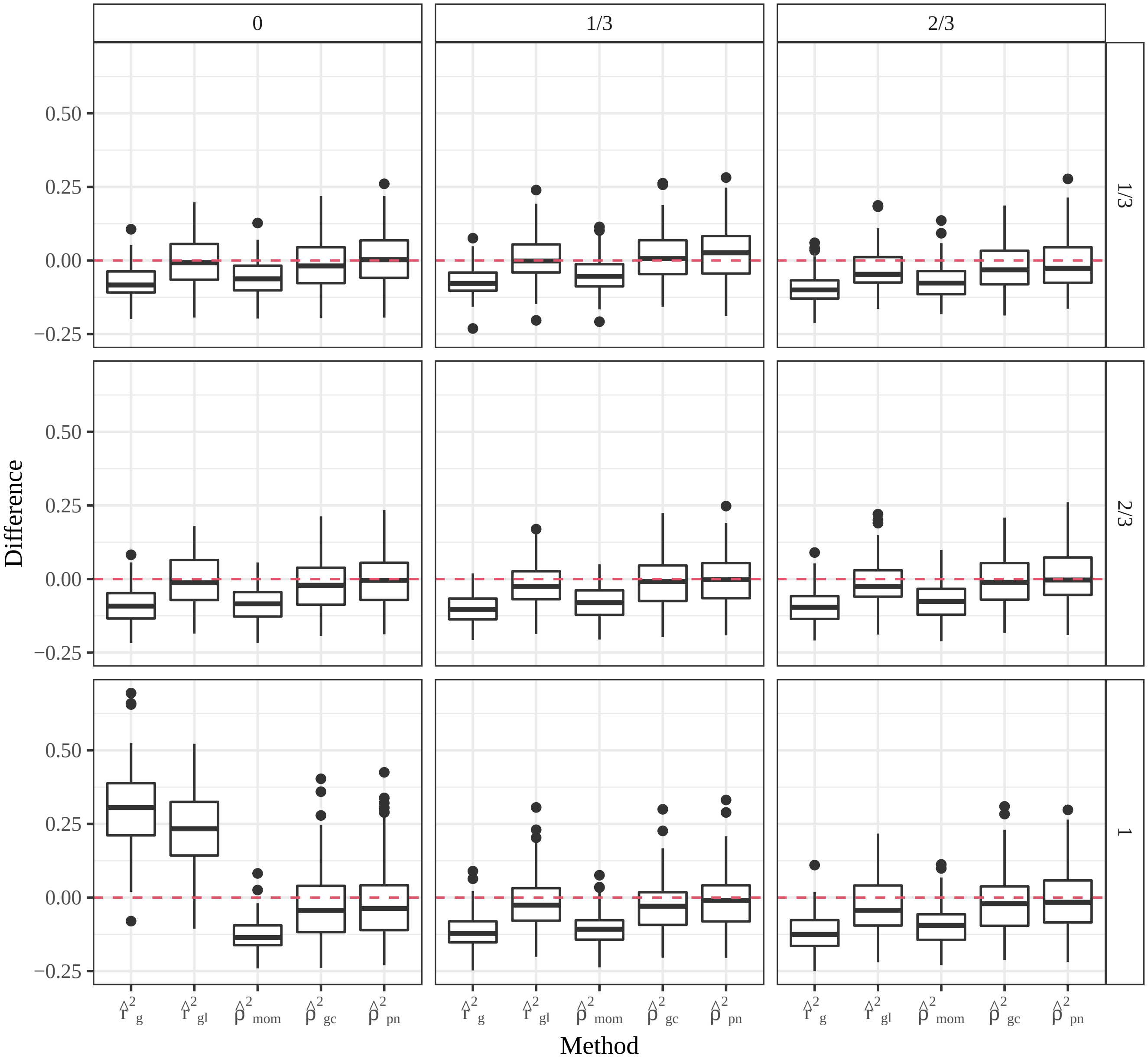


Figure S24: Boxplots of the difference between estimated $\rho^2$ and true $\rho^2$ ($y$-axis) for each method ($x$-axis) between two SNPs were located at 100 cM and 90 cM, and so should exibit moderate levels of LD (Figures S17 and S19). Values near zero (the dashed line) indicate unbiased behavior, while values over the dashed line indicate overestimation of $\rho^2$. Data were generated from 10 generations of random mating of a tetraploid species, where the founder population began at an allele frequency of 0.5, subgenome allele frequencies of 0.1 and 0.9, and an initial $\rho^2$ of 1. Populations were kept at a constant size of 1000 individuals from the final generation. Different levels of homologous pairing (row-facets) and quadrivalent formation (column facets) were explored. "True" $\rho^2$ was calculated using all 1000 individuals, and the estimated $\rho^2$ was calculated using a random sample of 100 individuals. The centromere is located at 50 cM.

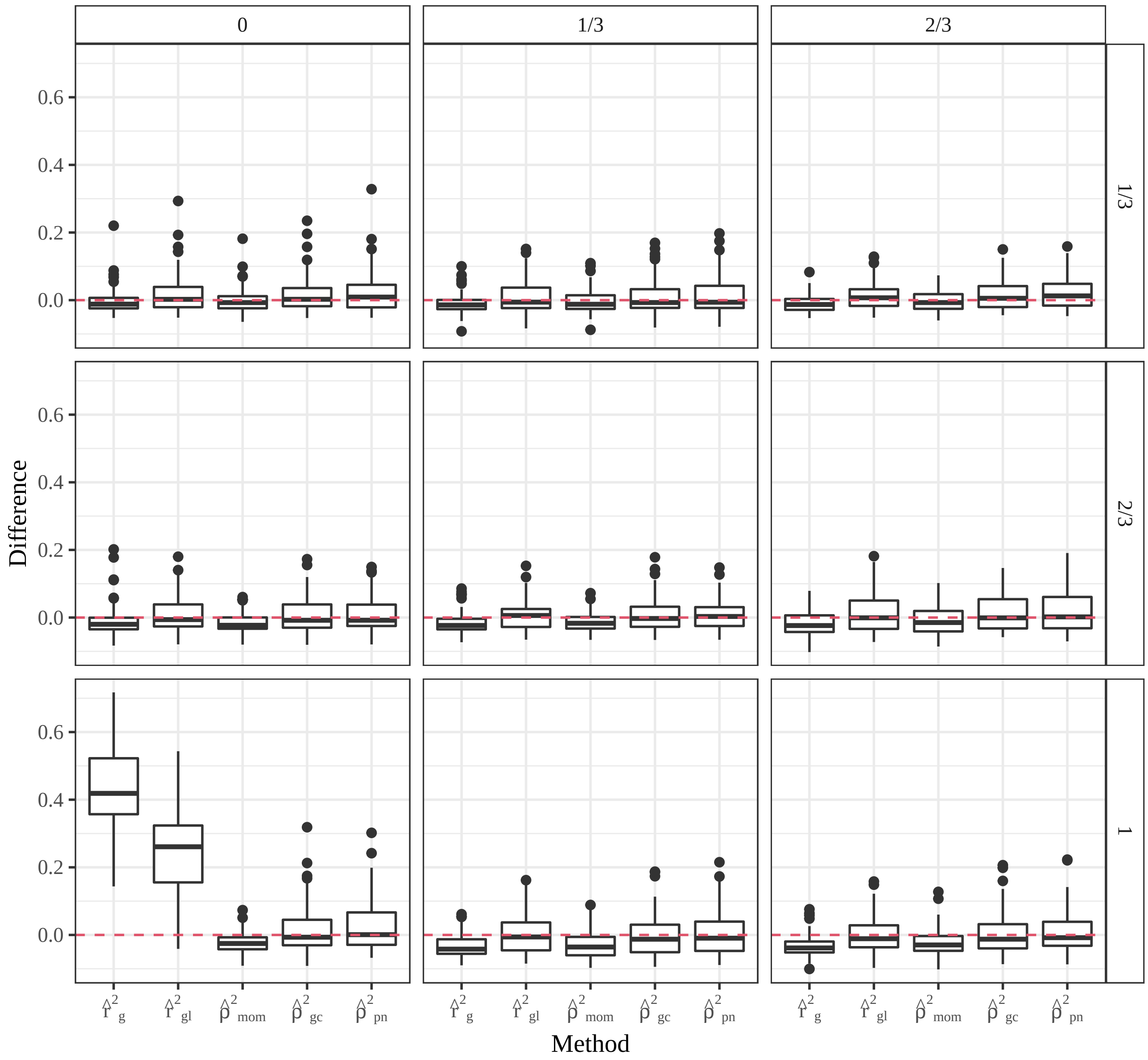


Figure S25: Boxplots of the difference between estimated $\rho^2$ and true $\rho^2$ ($y$-axis) for each method ($x$-axis) between two SNPs were located at 100 cM and 80 cM, and so should exibit LD close to 0 (Figures S17 and S19). Values near zero (the dashed line) indicate unbiased behavior, while values over the dashed line indicate overestimation of $\rho^2$. Data were generated from 10 generations of random mating of a tetraploid species, where the founder population began at an allele frequency of 0.5, subgenome allele frequencies of 0.1 and 0.9, and an initial $\rho^2$ of 1. Populations were kept at a constant size of 1000 individuals. Different levels of homologous pairing (row-facets) and quadrivalent formation (column facets) were explored. "True" $\rho^2$ was calculated using all 1000 individuals from the final generation, and the estimated $\rho^2$ was calculated using a random sample of 100 individuals. The centromere is located at 50 cM.

| Preferential Pairing | Quadrivalent Formation | Location (cM) | 0 | 1 | 2 | 3 | 4 |
|---|---|---|---|---|---|---|---|
| 1/3 | 0 | 50 | .06 | .26 | .38 | .24 | .06 |
| 2/3 | 0 | 50 | .06 | .25 | .38 | .25 | .06 |
| 1 | 0 | 50 | .01 | .15 | .69 | .14 | .01 |
| 1/3 | 1/3 | 50 | .06 | .26 | .38 | .24 | .06 |
| 2/3 | 1/3 | 50 | .06 | .25 | .37 | .25 | .06 |
| 1 | 1/3 | 50 | .07 | .24 | .38 | .25 | .07 |
| 1/3 | 2/3 | 50 | .06 | .25 | .38 | .25 | .06 |
| 2/3 | 2/3 | 50 | .06 | .25 | .38 | .25 | .06 |
| 1 | 2/3 | 50 | .06 | .25 | .38 | .26 | .06 |
| 1/3 | 0 | 100 | .06 | .24 | .39 | .25 | .06 |
| 2/3 | 0 | 100 | .05 | .24 | .41 | .25 | .06 |
| 1 | 0 | 100 | .01 | .15 | .68 | .14 | .01 |
| 1/3 | 1/3 | 100 | .06 | .24 | .38 | .25 | .06 |
| 2/3 | 1/3 | 100 | .06 | .25 | .38 | .25 | .06 |
| 1 | 1/3 | 100 | .05 | .23 | .40 | .26 | .06 |
| 1/3 | 2/3 | 100 | .07 | .24 | .36 | .25 | .08 |
| 2/3 | 2/3 | 100 | .07 | .25 | .37 | .24 | .07 |
| 1 | 2/3 | 100 | .07 | .24 | .37 | .26 | .07 |
| HWE Autopolyploid | | | .06 | .25 | .38 | .25 | .06 |
| HWE Allopolyploid | | | .01 | .15 | .69 | .15 | .01 |

Table S2: Distribution of genotypes (columns 0, 1, 2, 3, and 4) of SNPs at 50 cM and 100 cM using 10000 individuals simulated from 10 generations of random mating under various levels of homologous pairing and quadrivalent formation. The botton two rows contain the genotype distributions at equilibrium for autopolyploids and allopolyploids in the absence of double reduction. The effects of double reduction appear to be minor, and the effects of preferential pairing seem to only result in major deviations from autopolyploid HWE in absence of quadrivalent formation. In which case, the genotype distributions very closely follow the expected distribution in allopolyploid HWE.

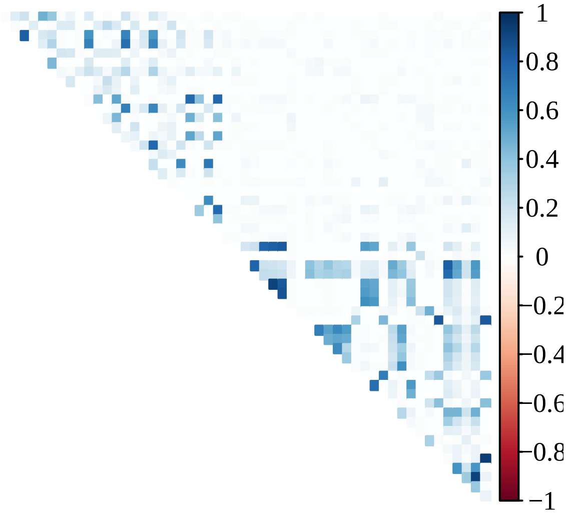


Figure S26: Heatmap of $\hat{r}_g^2$ from Section 3.3 using posterior mode genotypes. These *Solanum tuberosum* data come from Uitdewilligen et al. [2013].

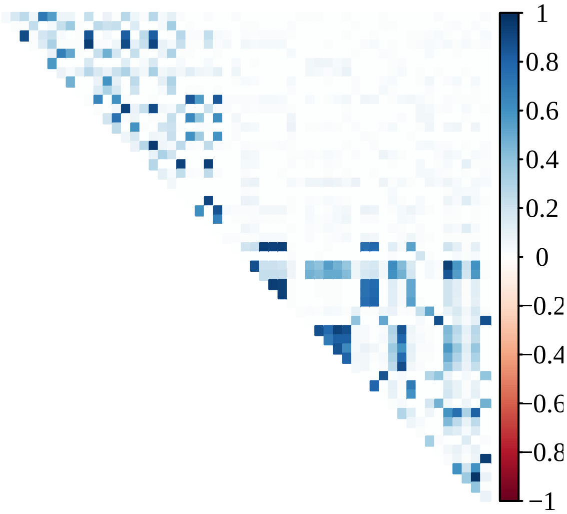


Figure S27: Heatmap of $\hat{r}_{gl}^2$ from Section 3.3. These *Solanum tuberosum* data come from Uitdewilligen et al. [2013].

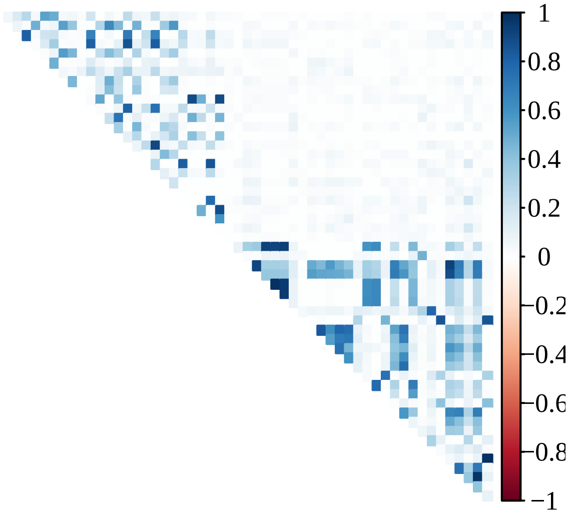


Figure S28: Heatmap of $\hat{\rho}^2_{mom}$ from Section 3.3 using posterior mean genotypes. These *Solanum tuberosum* data come from Uitdewilligen et al. [2013].

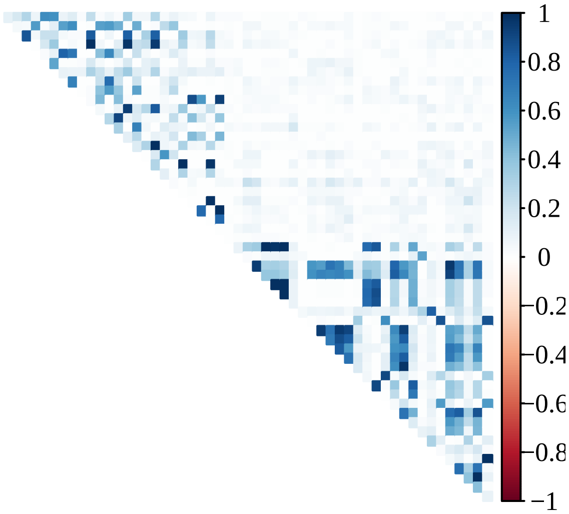


Figure S29: Heatmap of $\hat{\rho}^2_{gc}$ from Section 3.3. These *Solanum tuberosum* data come from Uitdewilligen et al. [2013].

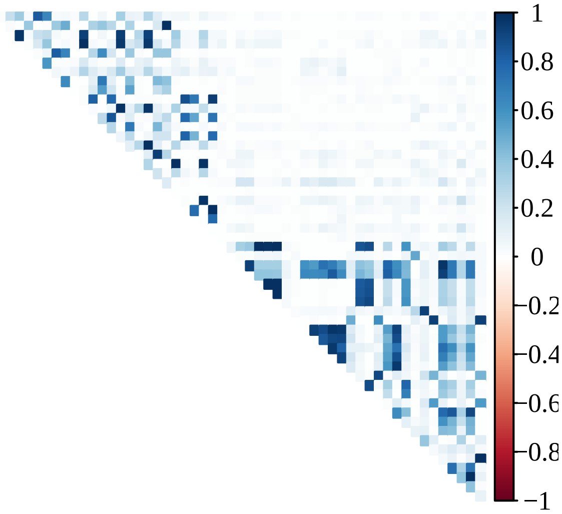


Figure S30: Heatmap of $\hat{\rho}^2_{pn}$ from Section 3.3. These *Solanum tuberosum* data come from Uitdewilligen et al. [2013].

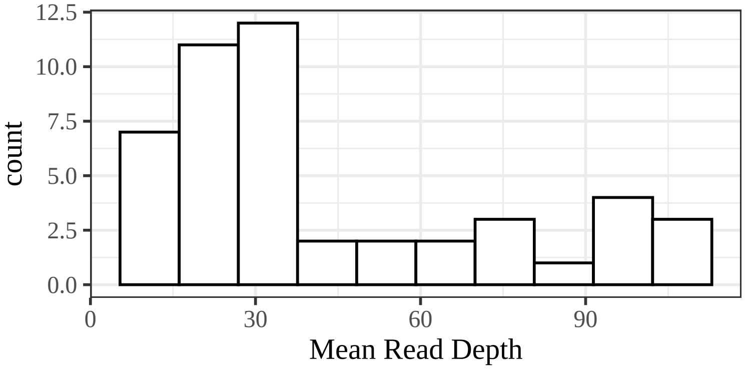


Figure S31: Histogram of mean read-depths of the SNPs used in Section 3.3 from the Uitdewilligen et al. [2013] *Solanum tuberosum* data.

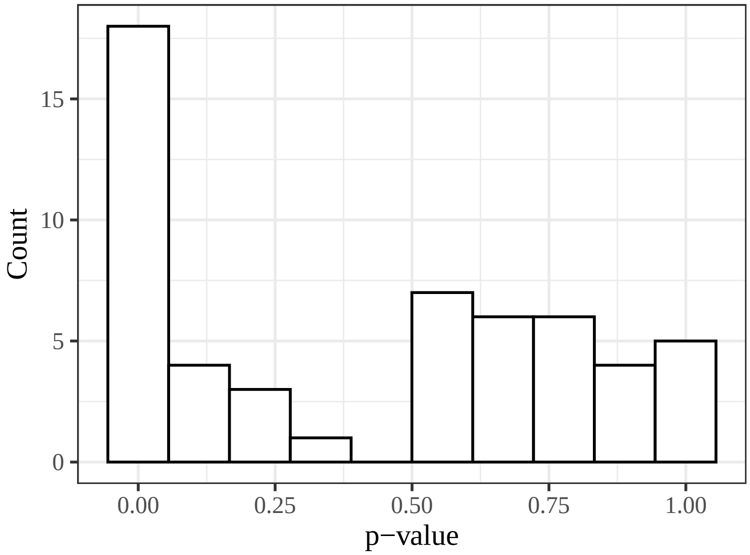


Figure S32: Histogram of $p$-values for tests of HWE using posterior mode genotypes of the SNPs used in Section 3.3 from the Uitdewilligen et al. [2013] *Solanum tuberosum* data. If most SNPs were in HWE, we would expect to see a histogram approximating a uniform distribution. The above histogram indicates that many SNPs deviate from HWE.

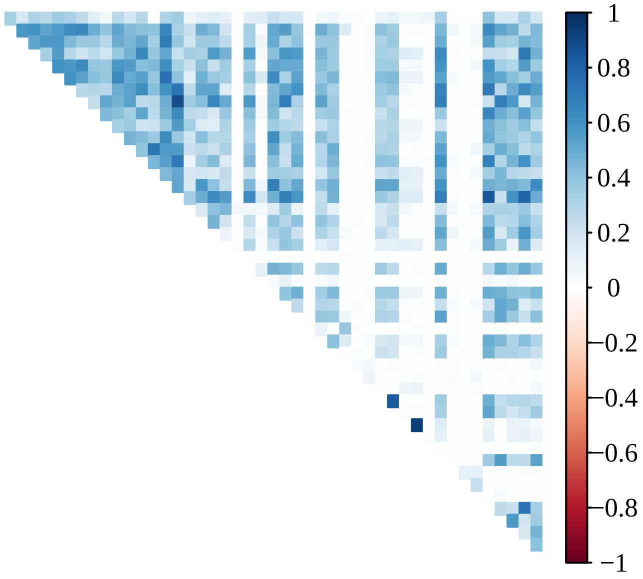


Figure S33: Heatmap of $\hat{r}_g^2$ from Section S13 using posterior mode genotypes. These *Andropogon gerardii* data come from McAllister and Miller [2016].

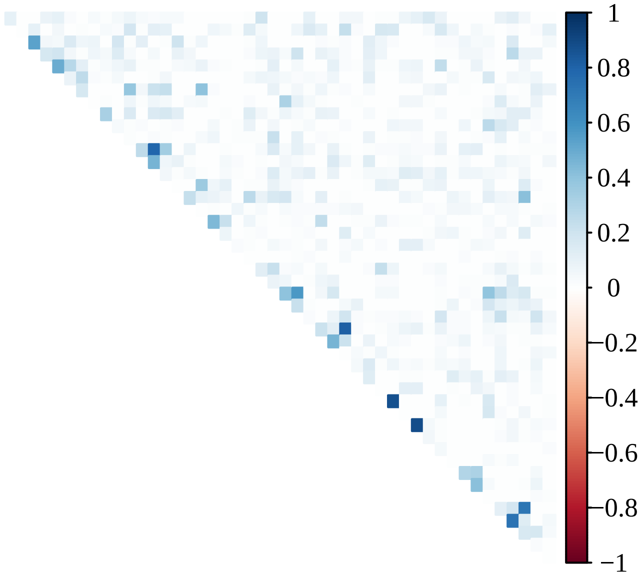


Figure S34: Heatmap of $\hat{r}^2_{gl}$ from Section S13. These *Andropogon gerardii* data come from McAllister and Miller [2016].

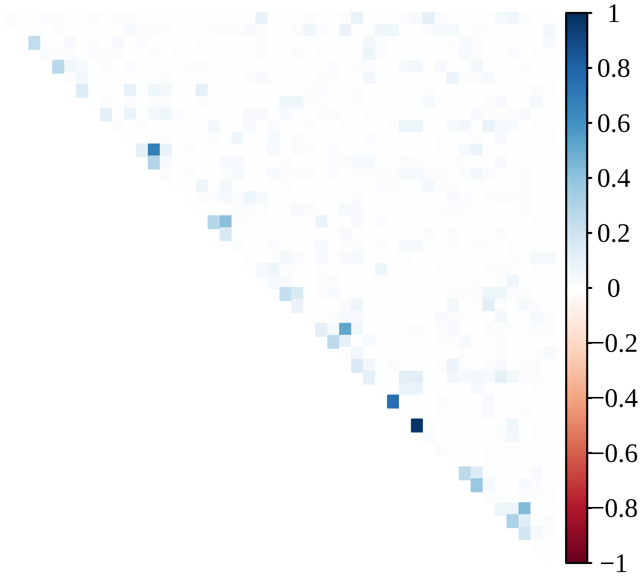


Figure S35: Heatmap of $\hat{\rho}^2_{mom}$ from Section S13 using posterior mean genotypes. These *Andropogon gerardii* data come from McAllister and Miller [2016].

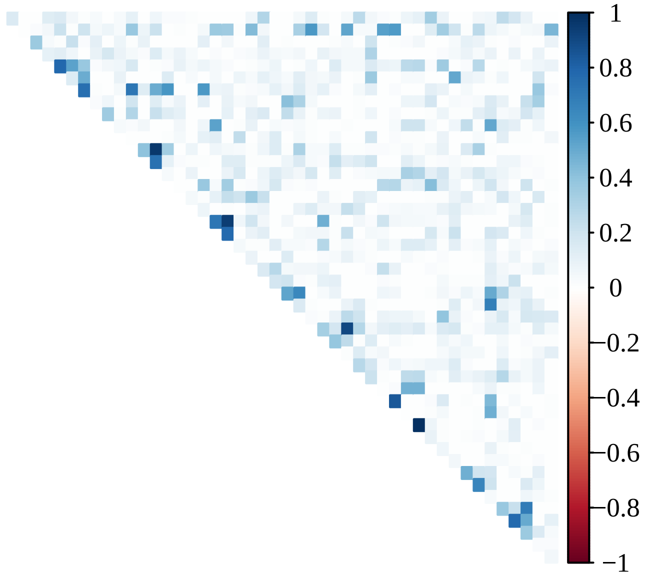


Figure S36: Heatmap of $\hat{\rho}^2_{gc}$ from Section S13. These *Andropogon gerardii* data come from McAllister and Miller [2016].

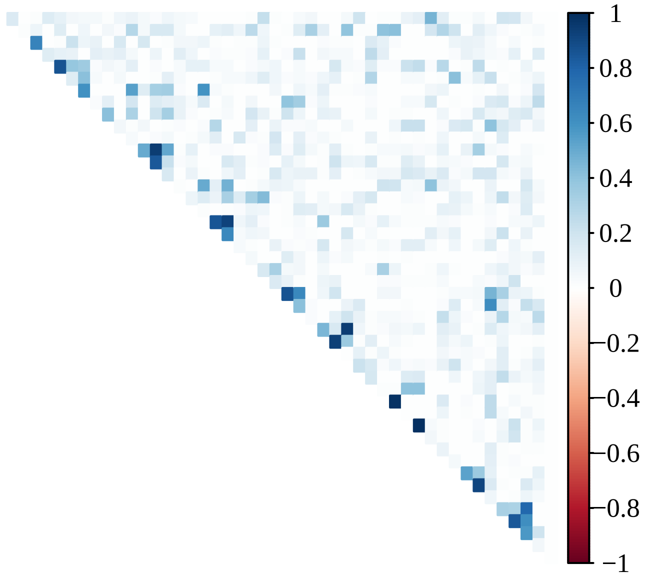


Figure S37: Heatmap of $\hat{\rho}^2_{pn}$ from Section S13. These *Andropogon gerardii* data come from McAllister and Miller [2016].

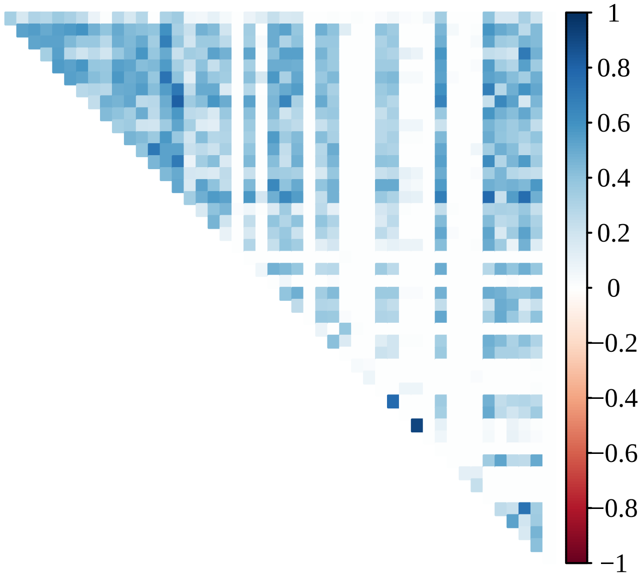


Figure S38: Heatmap of shrunken values of $\hat{r}_g^2$ from Section S13 using posterior mode genotypes. These *Andropogon gerardii* data come from McAllister and Miller [2016]. Shrinkage was done using the methods of Stephens [2016] and Dey and Stephens [2018].

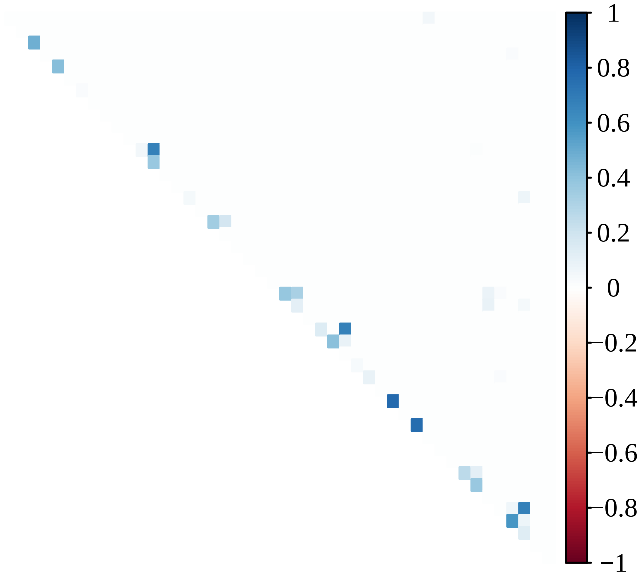


Figure S39: Heatmap of shrunken values of $\hat{r}_{gl}^2$ from Section S13. These *Andropogon gerardii* data come from McAllister and Miller [2016]. Shrinkage was done using the methods of Stephens [2016] and Dey and Stephens [2018].

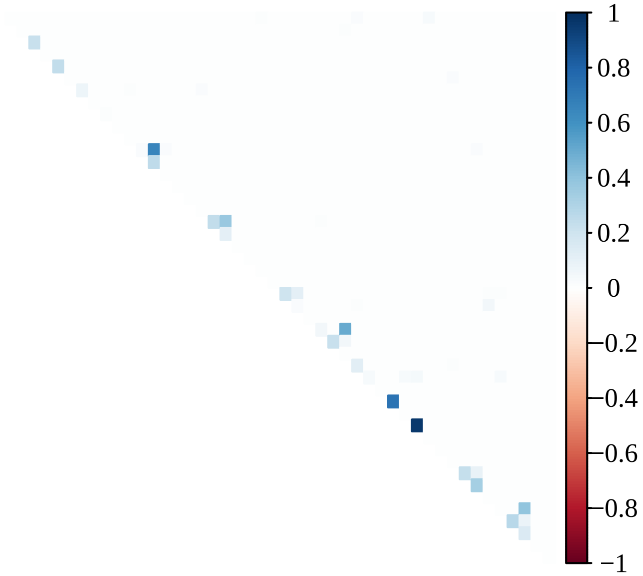


Figure S40: Heatmap of shrunken values of $\hat{\rho}^2_{mom}$ from Section S13 using posterior mean genotypes. These *Andropogon gerardii* data come from McAllister and Miller [2016]. Shrinkage was done using the methods of Stephens [2016] and Dey and Stephens [2018].

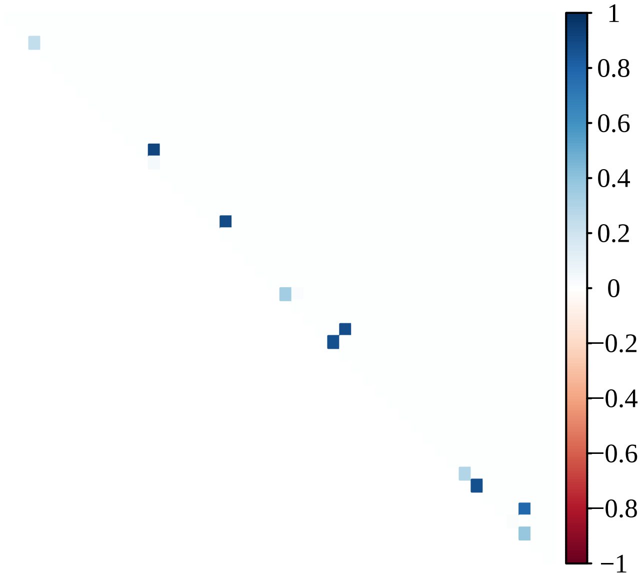


Figure S41: Heatmap of shrunken values of $\hat{\rho}^2_{pn}$ from Section S13. These *Andropogon gerardii* data come from McAllister and Miller [2016]. Shrinkage was done using the methods of Stephens [2016] and Dey and Stephens [2018].

### References


A. Agresti and B. A. Coull. Approximate is better than “exact” for interval estimation of binomial proportions. *The American Statistician*, 52(2):119–126, 1998. doi: 10.1080/00031305.1998.10480550.

M. S. Bartlett. On the theory of statistical regression. *Proceedings of the Royal Society of Edinburgh*, 53: 260–283, 1934. doi: 10.1017/S0370164600015637.

M. Betancourt. Cruising the simplex: Hamiltonian Monte Carlo and the Dirichlet distribution. *AIP Conference Proceedings*, 1443(1):157–164, 2012. doi: 10.1063/1.3703631.

C. C. Cockerham and B. S. Weir. Digenic descent measures for finite populations. *Genetical Research*, 30 (2):121–147, 1977. doi: 10.1017/S0016672300017547.

K. K. Dey and M. Stephens. CorShrink: Empirical Bayes shrinkage estimation of correlations, with applications. *bioRxiv*, 2018. doi: 10.1101/368316.

B. Efron. Bootstrap methods: Another look at the jackknife. *Ann. Statist.*, 7(1):1–26, 01 1979. doi: 10.1214/aos/1176344552.

T. S. Ferguson. *A course in large sample theory.* CRC Press, 2002. ISBN 0-412-04371-8.

R. A. Fisher. On the ‘probable error’ of a coefficient of correlation deduced from a small sample. *Metron*, 1: 3–32, 1921. URL `http://hdl.handle.net/2440/15169`.

E. A. Fox, A. E. Wright, M. Fumagalli, and F. G. Vieira. ngsLD: evaluating linkage disequilibrium using genotype likelihoods. *Bioinformatics*, 35(19):3855–3856, 03 2019. ISSN 1367-4803. doi: 10.1093/bioinformatics/btz200.

D. Gerard and L. F. V. Ferrão. Priors for genotyping polyploids. *Bioinformatics*, 36(6):1795–1800, 11 2019. ISSN 1367-4803. doi: 10.1093/bioinformatics/btz852. bioRxiv: 751784.

D. Gerard, L. F. V. Ferrão, A. A. F. Garcia, and M. Stephens. Genotyping polyploids from messy sequencing data. *Genetics*, 210(3):789–807, 2018. ISSN 0016-6731. doi: 10.1534/genetics.118.301468.

J. Haldane. Theoretical genetics of autopolyploids. *Journal of genetics*, 22(3):359–372, 1930. doi: 10.1007/BF02984197.

H. Hotelling. New light on the correlation coefficient and its transforms. *Journal of the Royal Statistical Society. Series B (Methodological)*, 15(2):193–232, 1953. ISSN 00359246. doi: 10.2307/2983768.

E. L. Lehmann and G. Casella. *Theory of point estimation.* Springer Science & Business Media, second edition, 1998. ISBN 0-387-98502-6.

S. Leonov and B. Qaqish. Correlated endpoints: simulation, modeling, and extreme correlations. *Statistical Papers*, 61(2):741–766, 2020. doi: 10.1007/s00362-017-0960-2.

H. Li. A statistical framework for SNP calling, mutation discovery, association mapping and population genetical parameter estimation from sequencing data. *Bioinformatics*, 27(21):2987, 2011. doi: 10.1093/bioinformatics/btr509.

C. A. McAllister and A. J. Miller. Single nucleotide polymorphism discovery via genotyping by sequencing to assess population genetic structure and recurrent polyploidization in *Andropogon gerardii*. *American Journal of Botany*, 103(7):1314–1325, 2016. doi: 10.3732/ajb.1600146.

C. A. McAllister and A. J. Miller. Data from: Single nucleotide polymorphism discovery via genotyping by sequencing to assess population genetic structure and recurrent polyploidization in *Andropogon gerardii*, 2017. URL `https://doi.org/10.5061/dryad.05qs7`. Dataset.

J. Morrison and T. Rajhathy. Frequency of quadrivalents in autotetraploid plants. *Nature*, 187(4736):528–530, 1960. doi: 10.1038/187528a0.

M. Stephens. False discovery rates: a new deal. *Biostatistics*, 18(2):275–294, 10 2016. ISSN 1465-4644. doi: 10.1093/biostatistics/kxw041.

J. G. A. M. L. Uitdewilligen, A.-M. A. Wolters, B. B. D'hoop, T. J. A. Borm, R. G. F. Visser, and H. J. van Eck. A next-generation sequencing method for genotyping-by-sequencing of highly heterozygous autotetraploid potato. *PLOS ONE*, 8(5):1–14, 05 2013. doi: 10.1371/journal.pone.0062355.

R. E. Voorrips and C. A. Maliepaard. The simulation of meiosis in diploid and tetraploid organisms using various genetic models. *BMC Bioinformatics*, 13(1):248, Sep 2012. ISSN 1471-2105. doi: 10.1186/1471-2105-13-248.

W. Whitt. Bivariate distributions with given marginals. *Ann. Statist.*, 4(6):1280–1289, 11 1976. doi: 10.1214/aos/1176343660.
